# Dengue virus harnesses mosquito Syntenin to load and secrete viral RNA into salivary exosomes

**DOI:** 10.1101/2025.08.11.669803

**Authors:** Florian Rachenne, Norman Schneider, Félix Rey-Cadilhac, Idalba Serrato-Pomar, Lauryne Pruvost, Laura Di Pietro, Jacques Dainat, Audrey Vernet, Chantal Cazevieille, Aurélie Ancelin, Joséphine Lai Kee Him, Sander Jansen, Elliott F. Miot, Jérémy Fraering, Laurianne Simon, Dorothée Missé, Marie Morille, Philippe Hammann, Martial Seveno, Serge Urbach, Kai Dallmeier, Éric Marois, Julien Pompon

## Abstract

Viruses exploit extracellular vesicles (EVs) to transfer infection-enhancing viral RNAs. However, mechanisms underlying viral RNA loading remain elusive. We leveraged our previous discovery that dengue virus secretes transmission-enhancing subgenomic flaviviral RNA (sfRNA) into mosquito salivary EVs to investigate viral RNA loading mechanism. We demonstrate that sfRNA alone promotes the secretion of sfRNA-containing EVs marked by the mosquito EV biogenesis protein AeSyntenin, by applying microscopy and viral genetic editing in *in vitro* and *in vivo* models. SfRNA via its stem loop structures interacts intracellularly with mosquito AeSyntenin and this interaction is selectively maintained within EVs as shown by complementary RNA-affinity chromatography and RNA immunoprecipitation, and AI-based prediction. Finally, we used systemic and salivary gland-specific protein depletion to establish a functional role for mosquito AeSyntenin in exosome production and salivary secretion of sfRNA. We propose that sfRNA binds AeSyntenin to drive its selective packaging and release into exosomes, elucidating a mechanism for viral RNA incorporation into EVs.

**Significance statement:** Viruses hijack extracellular vesicles (EVs) to enhance viral dissemination, but the mechanisms enabling selective viral RNA packaging into EVs remain unclear. Specifically, dengue virus transmission by mosquitoes relies on EV-based delivery of an immune-inhibitory subgenomic flaviviral RNA (sfRNA). Here, we uncover how dengue virus sfRNA is actively sorted into EVs from mosquito saliva. We show that sfRNA alone induces its secretion via EVs. We discover that sfRNA directly interacts with AeSyntenin intracellularly, and that this interaction persists in secreted EVs. Functional depletion studies reveal AeSyntenin’s role in salivary EV formation and sfRNA secretion. These findings establish a novel paradigm by which viral RNAs exploit vector EV pathways for dissemination.

## Introduction

Viruses exploit extracellular vesicles (EVs) to facilitate infection^1,2^. EVs are non-replicative, lipid bilayer structures secreted into the extracellular space and are broadly classified into two main categories: microvesicles (also known as ectosomes), which bud directly from the plasma membrane^3,4^, and exosomes, which originate from the endosomal pathway^3,5–7^. Exosome biogenesis involves Syntenin-1-mediated recruitment via two PDZ domains of the Endosomal Sorting Complex Required for Transport (ESCRT) to form intraluminal vesicles (ILVs) within multivesicular bodies (MVBs)^8–10^, which subsequently fuse with the plasma membrane to release exosomes^11,12^. EVs are selectively loaded with RNA and act as intercellular carriers, transferring functional RNA molecules between cells^13–16^. Viruses divert EV carriers to package and transfer viral RNAs (complete or fragment of viral genome) into other cells, activating various strategies to enhance infection^1^. Although the presence of viral RNA within EVs is well-documented for multiple virus families^1,17,18^, the mechanism(s) governing viral RNA loading into EVs has not been elucidated.

Our team discovered that mosquito-borne dengue viruses (DENV) and other Orthoflaviviruses - Flaviviridae family - secrete a subgenomic flaviviral RNA (sfRNA) inside EVs from mosquito saliva^19,20^. SfRNA is an RNA decay product, resulting from partial degradation of the viral RNA genome (gRNA) by 5’-3’ exoribonucleases which stall on secondary structures in the 3’UTR^21^. Secondary structures for DENV include two stem loops (i.e., SL1 and SL2), two dumbbells (i.e., DB1 and DB2) and a final 3’ stem loop (3’SL). SfRNA is abundant in salivary glands^22^, which produce saliva, and is deposited via salivary EVs at the mosquito bite site^19,20^. In the bitten skin, sfRNA anti-immune properties^23^ then enhance local infection and facilitate transmission^19,20^, highlighting sfRNA-containing EVs as critical enhancers of pathogen dissemination. Here, we report how viral sfRNA is loaded and secreted within exosomes. We combined cell biology, molecular biology and cutting-edge genome editing using *in vitro* and *in vivo* approaches to show that sfRNA harnesses mosquito Syntenin homolog (AeSyntenin) to induce secretion of cellular and salivary exosomes loaded with sfRNA. Our results reveal a molecular mechanism for viral RNA packaging into EVs.

## Results

### DENV infection stimulates EV secretion in mosquitoes

To evaluate the impact of DENV infection on EV production, we infected Aag2 mosquito cells with low and high multiplicity of infection (MOI), and quantified EVs at 72 hours post-infection (hpi) (Fig. 1a). Compared to the low inoculum, the high inoculum produced higher infection intensity and quantity of secreted virions, as indicated by intracellular and extracellular gRNA copies, respectively - and higher quantity of produced and secreted sfRNA, as indicated by intracellular and extracellular sfRNA quantities, respectively (Fig. S1a-d). To determine the effect on EVs, we observed that two established mosquito EV markers, AeSyntenin and the homolog of human CD63 tetraspanin (hCD63)^3,24^, increased in the cell media with infection in a dose-dependent manner (Fig. 1b-d; Fig. S2a), indicating enhanced EV marker release. In contrast, intracellular levels of these markers at the protein (Fig. 1e-g; Fig. S2b) and mRNA (Fig. S3a) levels remained unchanged. Nanoparticle tracking analysis (NTA) results confirmed an increase in secreted particles (Fig. 1h) and the average size (Fig. 1i) upon infection, with a uniform concentration increase across EV subpopulations (Fig. 1j,k), illustrated by the conserved proportions of EV subtypes (Fig. 1l,m).

**Fig. 1.**
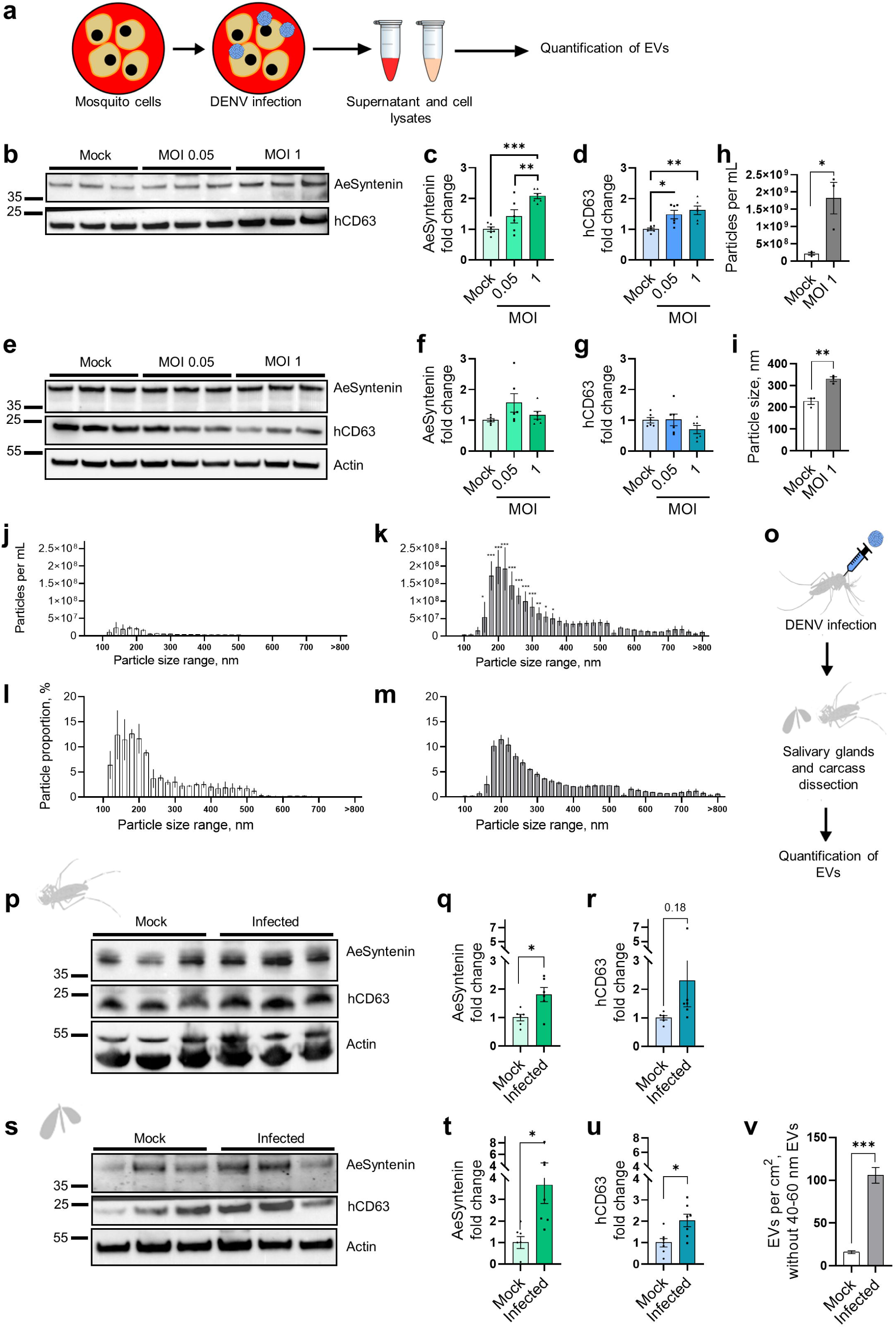
DENV infection stimulates secretion of EVs in cells and mosquitoes. **a** Assessing the effect of DENV infection on EV secretion in Aag2 mosquito cells. **b-d** Representative WB (b) and quantification of AeSyntenin (c) and hCD63 (d) in media from cells infected with MOI 0.05 or 1. Each lane corresponds to one replicate. **e-g** Representative WB (e) and quantification of AeSyntenin (f) and hCD63 (g) in cells infected with MOI 0.05 or 1. Actin was used as a loading control. Each lane corresponds to one replicate. **h,i** Particle concentration (h) and size (i) in media from cells infected with MOI 1 by NTA. **j-m** Size-range (20 nm width) particle concentration in media from cells mock-infected (j) and infected with MOI 1 (k), and size-range proportion in media from cells mock-infected (l) and infected with MOI 1 (m) by NTA. **o** Assessing the effect of DENV infection on EV secretion in mosquitoes. **p-r** Representative WB (p) and quantification of AeSyntenin (q) and hCD63 (r) in mock and infected carcasses. Actin was used as a loading control. Each lane corresponds to one replicate. **s-u** Representative WB (s) and quantification of AeSyntenin (t) and hCD63 (u) in mock and infected salivary glands. Five pairs of salivary glands were combined in each sample. Actin was used as a loading control. Each lane corresponds to one replicate. **v** EVs per cm² of pictures. N pictures, 40 and 35 for mock and infected saliva, respectively. Particles from 40-60 nm were not counted. c,d,f-m,q,r,t,u,v Bars represent mean ± s.e.m. and dots indicate repeats. j-m N, 3. c,d,f,g *, p < 0.05; **, p < 0.01; ***, p < 0.001, as determined by ANOVA post-hoc Fisher’s LSD test. h,i,q,r,t,u,v *, p < 0.05; **, p < 0.01; ***, p < 0.001, as determined by T-test. j-m *, p < 0.05; **, p < 0.01; ***, p < 0.001, as determined by ANOVA’s post hoc FDR test between Ctl. RNA and sfRNA within size range.

We examined the effect of DENV infection on EVs *in vivo* by infecting mosquitoes and analyzing tissues and saliva 10 days post infection (Fig. 1o), when infection was established and sfRNA produced in carcasses and salivary glands (Fig. S1e-h). Both AeSyntenin and hCD63 levels were elevated in infected carcasses (Fig. 1p-r; Fig. S2c), indicative of enhanced EV biogenesis, and in infected salivary glands (Fig. 1s-u; Fig. S2d), suggesting an increased production of salivary EVs. Consistent with *in vitro* findings, *AeSyntenin* mRNA levels remained unchanged in both carcasses and salivary glands (Fig. S3b, c). Finally, we used TEM to assess EVs in saliva from infected mosquitoes. Salivary EVs were mostly smaller than 40 nm (Fig. S4, S5) and to avoid biasing EV quantification by counting virions, we did not count EVs between 40 and 60 nm. TEM analysis showed that infection significantly increased EV secretion in saliva (Fig. 1v). Together, these results indicate that DENV infection enhances salivary EV production, likely by activating secretion instead of biogenesis, as intracellular EV markers remained stable.

### DENV sfRNA induces secretion of EV-packaged sfRNA

To determine whether sfRNA is required for the infection-induced EV secretion, we infected Aag2 mosquito cells with a DENV infectious clone (IC) carrying deletions in the RNA pseudo-knot 1 (ΔPk1), which confers exoribonuclease resistance^25,26^, to impair sfRNA biogenesis (Fig. 2a). The ΔPk1 IC had a reduced sfRNA concentration, as quantified by the diminished sfRNA:gRNA ratio compared to non-mutated sfRNA IC (Fig. 2b). Interestingly, infection with ΔPk1 IC reduced and abrogated the infection-induced secretion of AeSyntenin and hCD63, respectively (Fig. 2c-e; Fig. S6a). Intracellular EV marker levels remained unchanged by infection (Fig. 2f-h; Fig. S6b), consistent with prior observations upon infection with wild-type DENV. To assess whether sfRNA is sufficient to induce EV production, we transfected *in vitro*-transcribed sfRNA or a size-matched viral control RNA (Ctl. RNA) and quantified EVs 72h later (Fig. 2i). Additionally, to evaluate whether Pk1 structure was necessary for EV induction, we transfected ΔPk1 sfRNA. All RNA fragments - sfRNA, ΔPk1 sfRNA and control RNA - were homogenously delivered intracellularly after adjusting for transfection efficiency variability (Fig. S7a,b). Strikingly, sfRNA transfection increased AeSyntenin secretion without affecting hCD63 (Fig. 2j-l; Fig. S8a), while intracellular levels of both EV markers remained unchanged as compared to control RNA (Fig. 2m-o; Fig. S8b). In contrast, ΔPk1 sfRNA transfection did not alter extracellular and intracellular AeSyntenin but induced a modest increase in extracellular and intracellular hCD63 (Fig. 2j-o; Fig. S8a,b). Although NTA analysis of ultracentrifugated EVs showed no major changes in total particle number (Fig. 2p) and mean particle size (Fig. 2q) following non-mutated sfRNA transfection, size distribution analysis revealed a specific increase in the proportion of small particles (∼ 40 nm) together with a sharp decrease in EVs ∼ 60 nm (Fig. 2r,s). Collectively, these results suggest that sfRNA is sufficient to drive EV secretion, and that induction of this process depends on the integrity of the Pk1 structure.

**Fig. 2.**
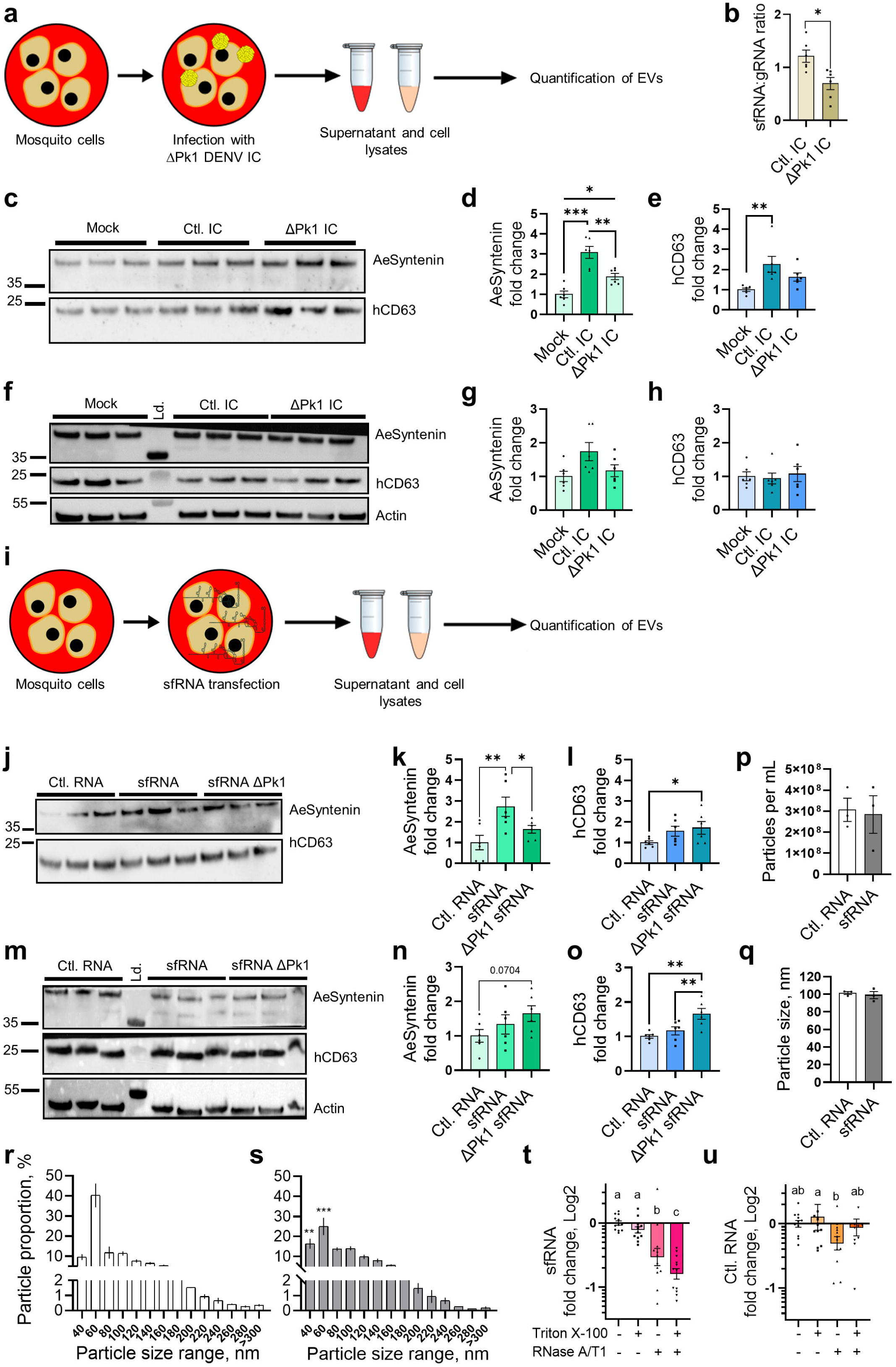
sfRNA stimulates secretion of EVs. **a** Assessing the effect of sfRNA-deficient DENV [ΔPk1 DENV infectious clone (IC)] infection on EV secretion in Aag2 mosquito cells. Control cells were infected with non-mutated sfRNA IC (Ctl. IC). **b** Levels of intracellular sfRNA:gRNA ratio. **c-e** Representative WB (c) and quantification of AeSyntenin (d) and hCD63 (e) in media from cells infected with mock, ΔPk1 sfRNA and non-mutated sfRNA DENV IC (Ctl. IC). **f-h** Representative WB (f) and quantification of AeSyntenin (g) and hCD63 (h) in cells infected with mock, ΔPk1 sfRNA and non-mutated sfRNA DENV IC. Actin was used as a loading control. Each lane corresponds to one replicate. **i** Assessing the effect of sfRNA and ΔPk1 sfRNA transfection on EV secretion in Aag2 mosquito cells. A same-size viral RNA fragment was transfected as control (Ctl. RNA). **j-l** Representative WB (j) and quantification of AeSyntenin (k) and hCD63 (l) in media from cells transfected with sfRNA, ΔPk1 sfRNA or Ctl. RNA. **m-o** Representative WB (m) and quantification of AeSyntenin (n) and hCD63 (o) in cells transfected with sfRNA, ΔPk1 sfRNA or Ctl. RNA. Actin was used as a loading control. Each lane corresponds to one replicate. **p-q** Particle concentration (p) and size (q) in EVs isolated from cell media after transfection with sfRNA and Ctl. RNA. N, 3. **r,s** Size-range (20 nm width) particle proportion in media from cells transfected with Ctl. RNA (r) or sfRNA (s) by NTA. N, 3. **t,u** Levels of secreted RNase-resistant sfRNA (t) and Ctl. RNA (u) with or without Triton X-100 pre-treatment. b,d,e,g,h,k,l,n-u Bars show mean ± s.e.m. and dots indicate repeats. Ld., protein size ladder. b *, p < 0.05 as determined by T-test. d,e,g,h,k,l,n,o *, p < 0.05; **, p <0.01; ***, p < 0.001, as determined by ANOVA post-hoc Fisher’s LSD test. r,s **, p < 0.01; ***, p < 0.001, as determined by ANOVA post hoc FDR test between Ctl. RNA and sfRNA within size range. t,u Different letters indicate significant differences as determined by ANOVA post hoc Fisher’s LDS test.

We next investigated whether transfected sfRNA is secreted inside lipid-based vesicles. We submitted sfRNA present in cell media at 72h post-transfection to a RNase resistance assay – a method previously applied to detect sfRNA packaging within EVs^19,20^. Despite multiple post-transfection washes, residual sfRNA was consistently detected in the media immediately after transfection. This may reflect rapid secretion or contamination from transfection reagents. To ensure that residual transfecting sfRNA did not bias the RNase protection evaluation for secreted sfRNA, we showed that sfRNA encapsulated in transfecting liposomes was not protected from RNase digestion (Fig. S9a). We then observed that sfRNA from transfected cell media was partially degraded by RNase, consistent with free unprotected sfRNA^19,20^. However, pre-treatment with detergent favored RNase degradation (Fig. 2t), indicating that a fraction of sfRNA was protected within lipid membranes. To test whether this packaging is specific to sfRNA, we repeated the RNase assay with Ctl. RNA, which was similarly unprotected by liposomes (Fig. S9b). In contrast to sfRNA, Ctl. RNA was similarly degraded with or without detergent (Fig. 2u), suggesting it was not vesicle-associated. Together, these results suggest that sfRNA alone is sufficient to drive the secretion of AeSyntenin-positive EVs and to be selectively packaged into EVs.

### sfRNA from DENV and other orthoflaviviruses interacts with AeSyntenin

We hypothesized that DENV sfRNA is packaged into EVs through interactions with EV-associated proteins and investigated this by performing RNA-affinity chromatography^19,27^ on EV lysates from Aag2 mosquito cells (Fig. 3a). Compared to a size-matched Ctl. RNA, MS analysis identified 50 proteins that were enriched more than 1.5-fold with DENV sfRNA, including proteins involved in EV biogenesis and previously detected within EVs (Fig. 3b; Dataset S1). Notably, AeSyntenin was enriched with sfRNA, and its interaction with sfRNA was validated by Western blot (WB) (Fig. 3c; Fig. S10a,b). Repeating the RNA-affinity chromatography on cell lysates, we also detected the sfRNA-AeSyntenin interaction (Fig. 3d; Fig. S10c,d), suggesting that the interaction is initiated intracellularly. We further confirmed this interaction using RNA-immunoprecipitation (RIP) with anti-AeSyntenin antibody (Fig. 3e) and detected sfRNA bound to AeSyntenin in both EVs (Fig. 3f,g; Fig. S10e,f) and cell lysates from DENV-infected cells (Fig. 3i,j; Fig. S10g,h). Interestingly, DENV gRNA was also enriched in AeSyntenin RIP eluates from both EVs and cells (Fig. 3h,k). To evaluate AeSyntenin binding specificity, we measured enrichment of *AeActin*, *AeRPS7*, and *AeSyntenin* mRNAs. While all three mRNAs interacted with AeSyntenin intracellularly (Fig. S11a-c), none were recovered in RIP eluates from EVs (Fig. S11d-f), suggesting a selective EV loading mechanism. AI-based prediction of protein-RNA interactions with RoseTTAFoldNA^28^ identified putative interactions between the AeSyntenin PDZ domains (aa 150-331) and the RNA stems in SL1, SL2 and 3’SL structures of DENV sfRNA (Fig. S12). To validate these binding segments, we performed RNA-affinity chromatography with DENV sfRNA fragments lacking either both DBs (ΔDBs), both SLs (ΔSLs) or 3’SL (Δ3’SL) (Fig. 3l). Compared with the full-length sfRNA, AeSyntenin enrichment was reduced in both intracellular and extracellular fractions when the SLs and 3’SL structures were absent (Fig. 3m-p; Fig. S13a,b). To extend our findings to orthoflaviviruses, we repeated the RIP assay in Aag2 cells infected with either West Nile virus (WNV) or Zika virus (ZIKV), and predicted interactions with their respective sfRNA. For both viruses, AeSyntenin interacted with sfRNA in intracellular and extracellular fractions, whereas interactions with gRNA were detected only intracellularly (Fig. S14; S15). AI-based structural predictions further supported these results by confirming the binding potential of AeSyntenin to stem-loop structures of WNV and ZIKV sfRNAs (Fig. S16; S17). Collectively, these results reveal that AeSyntenin binds stem-loop structures in sfRNA from multiple orthoflaviviruses intracellularly, as well as other RNA fragments, yet only the interaction with sfRNA persists within secreted EVs.

**Fig. 3.**
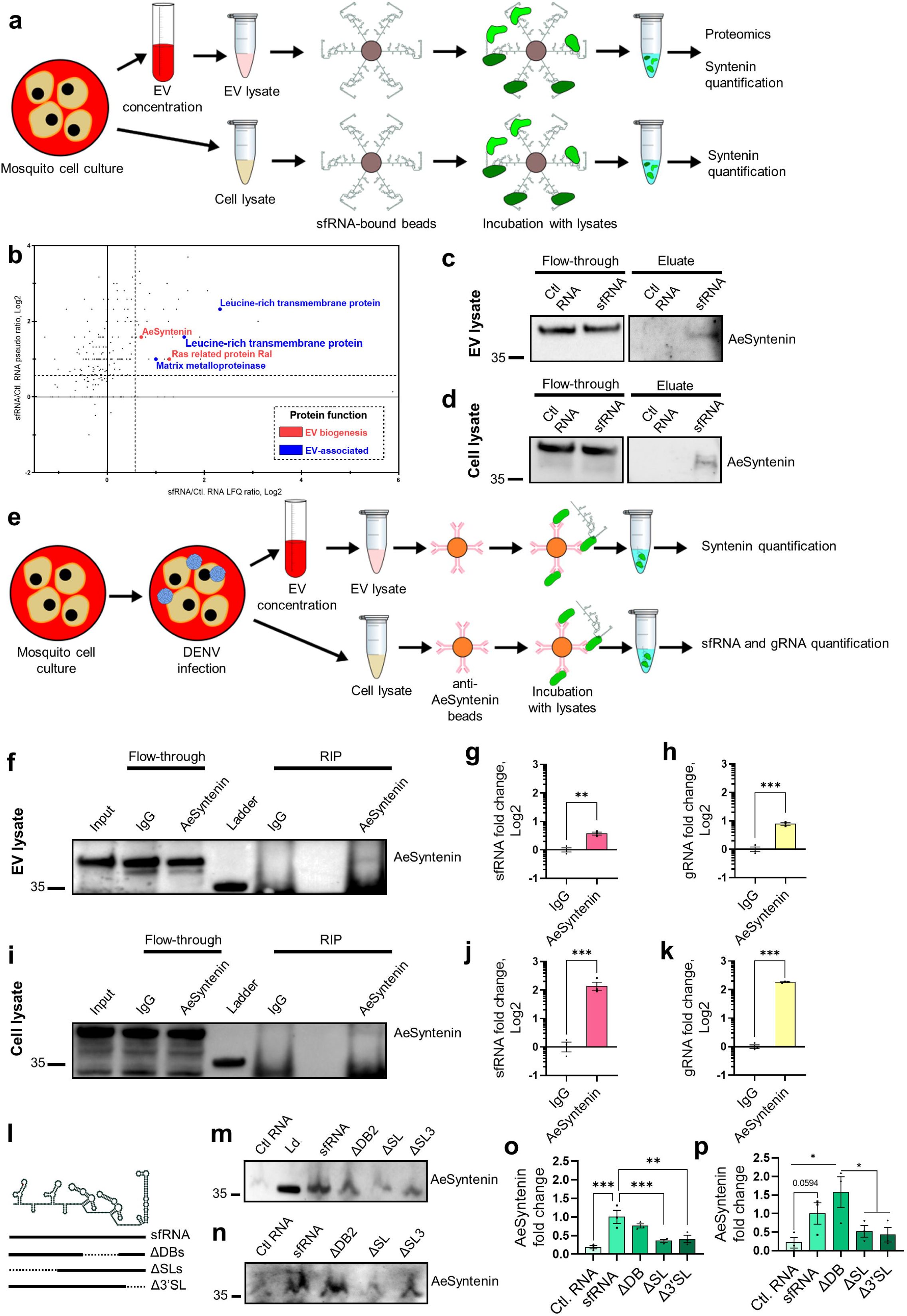
sfRNA interacts with AeSyntenin. **a** Identification of sfRNA-interacting proteins by RNA-affinity chromatography in Aag2 mosquito cells. A same-sized viral control RNA (Ctl. RNA) fragment was used as control. **b** Label-free quantification (LFQ) and pseudo ratio for sfRNA-interacting proteins as determined by MS analysis. Proteins with putative functions associated with EVs, EV biogenesis, RNA binding or Golgi trafficking are identified with different colors. N, 1. **c, d** AeSyntenin detection in flow-through and eluate from RNA-affinity chromatography conducted with EV (c) and cell (d) lysates. **e** Identification of AeSyntenin-interacting viral RNAs in EVs and cells infected with DENV by RNA immunoprecipitation. **f-h** Detection in flow-through and RNA immunoprecipitates (RIP) of AeSyntenin (f), sfRNA (g) and gRNA (h) levels precipitated with anti-AeSyntenin and IgG control in EV lysates. **i-k** Detection in flow-through and RNA immunoprecipitates (RIP) of AeSyntenin (i), sfRNA (j) and gRNA (k) levels precipitated with anti-AeSyntenin and IgG control in cell lysates. **l** Representation of the full length sfRNA, ΔSL sfRNA, ΔDB sfRNA and Δ3’SL sfRNA fragments. **m, n** AeSyntenin detection from RNA-affinity chromatography conducted with cell lysate (m) and EVs (n) from Aag2 mosquito cells. **o, p** AeSyntenin levels in RNA-affinity chromatography eluates from cell lysate (o) and EVs (p). g,h,j,k,o,p Bars represent mean ± s.e.m. Dots indicate repeats. **, p < 0.01; ***, p < 0.001, as determined by T (g,h,j,k) test or by ANOVA post-hoc FDR test (o,p).

### AeSyntenin is required for sfRNA secretion in saliva

We started by evaluating functional homology between AeSyntenin and Human Syntenin-1. Although we previously identified sequence homology in the functional PDZ domains between the two proteins^24^, we assessed structural homology by using AlphaFold^29^ and ScanProsite^30^. Blast between AeSyntenin and the Human Syntenin-1 showed a match in the globular structure corresponding to the PDZ domains, which were predicted with high confidence (Fig. S18a-c,h). PDZ domains were similarly identified in Syntenin-1 homologs from *Aedes albopictus* (AaSyntenin) (Fig. S18d,e,h) and *Culex quinquefasciatus* (CqSyntenin) (Fig. S18f-h). We further assessed AeSyntenin functional homology by identifying interacting partners in mosquito EVs using co-immunoprecipitation (co-IP) on EVs secreted from DENV-infected Aag2 mosquito cells, for relevance to AeSyntenin’s role in sfRNA loading. Among several EV-associated proteins identified (Fig. S19; Dataset S2), we detected the mosquito homolog of Human ALIX, a known partner with which Syntenin-1 induces EV biogenesis and cargo loading^9,10^. Our structural and interactome results suggest functional homology between AeSyntenin and Syntenin-1.

To assess whether AeSyntenin is required for sfRNA secretion in saliva, we silenced *AeSyntenin* in mosquitoes by injecting dsRNA, followed four days later, by intrathoracic DENV inoculation. Ten days post-infection, we quantified gRNA and sfRNA in salivary glands and saliva (Fig. 4a). While mosquito survival was unaffected at the time of inoculation, it was reduced at day 10 in the AeSyntenin-depleted, DENV-infected group (Fig. S20a,b), suggesting either a longer-term effect of AeSyntenin depletion or a combined deleterious effect of AeSyntenin depletion and infection. At the time of saliva collection, we confirmed AeSyntenin depletion in carcasses (Fig. 4b) and salivary glands (Fig. 4c,d). In salivary glands, AeSyntenin depletion reduced gRNA levels (Fig. 4e), suggesting a pro-viral function potentially linked to its interaction with gRNA (Fig. 3j,k). SfRNA levels were also reduced (Fig. 4f), consistent with lower gRNA template availability; however, the sfRNA:gRNA ratio remained unchanged (Fig. 4g), indicating that AeSyntenin does not influence sfRNA biogenesis. In contrast, in saliva, while gRNA levels were unaffected (Fig. 4h), sfRNA levels were significantly decreased following AeSyntenin depletion (Fig. 4i), resulting in a marked reduction in the sfRNA:gRNA ratio (Fig. 4j).

**Fig. 4.**
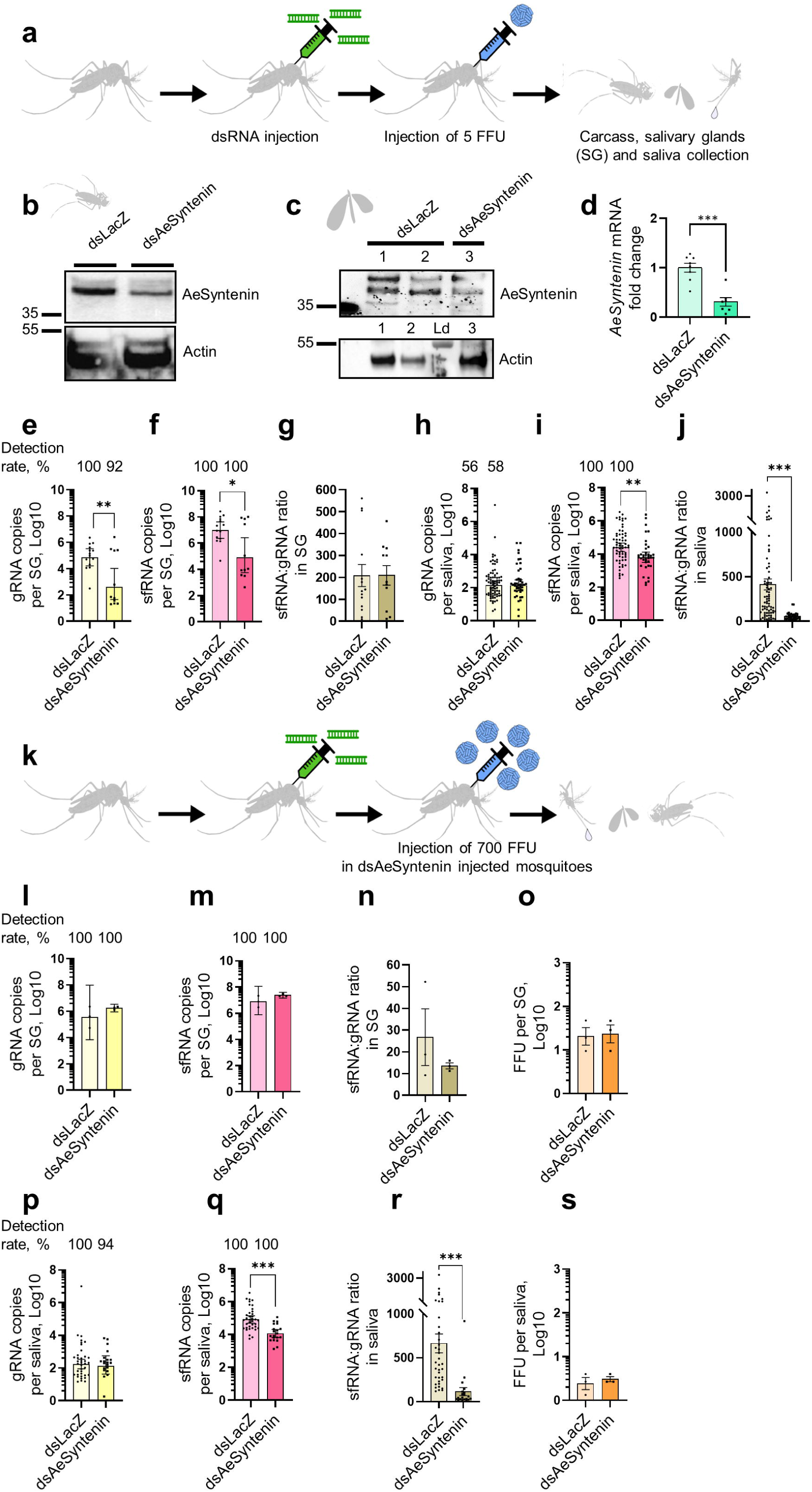
AeSyntenin depletion in mosquitoes reduces sfRNA secretion in saliva. **a** Assessing the effect of AeSyntenin depletion on gRNA and sfRNA levels in carcasses, salivary glands (SG) and saliva. Mosquitoes were injected with dsRNA against AeSyntenin or LacZ as control before inoculation with 5 FFU of DENV. **b,c** AeSyntenin levels in carcass (b) and in SG (c). Actin was used as a loading control. Each lane corresponds to one replicate. Two replicates were shown for dsLacZ SG. Ld., protein ladder. **d** *AeSyntenin* mRNA levels in SG. **e-g** Levels of gRNA (e), sfRNA (f) and sfRNA:gRNA ratio (g) in SG. **h-j** Levels of gRNA (h), sfRNA (i) and sfRNA:gRNA ratio (j) in saliva. **k** Assessing the effect of AeSyntenin depletion on gRNA and sfRNA levels in carcasses, salivary glands (SG) and saliva after compensating for AeSyntenin effect on SG infection levels. Mosquitoes injected with dsAeSyntenin were inoculated with 700 FFU of DENV, whereas mosquitoes injected with dsLacZ were injected with 5 FFU. **k-o** Levels of gRNA (k), sfRNA (m), sfRNA:gRNA ratio (n) and infectious particles (o) in SG. **p-s** Levels of gRNA (p), sfRNA (q), sfRNA:gRNA ratio (r) and infectious particles (s) in saliva. d,g,j,n,o,r,s Bars represent mean ± s.e.m. and dots indicate repeats. e,f,h,i,l,m,p,q Bars represent geometric mean ± 95% C.I. and dots indicate repeats. FFU, focus forming unit. *, p < 0.05; **, p < 0.01; ***, p < 0.001, as determined by T-test.

To account for the reduced salivary gland infection following AeSyntenin depletion - which could confound the analysis of sfRNA biogenesis and secretion - we repeated the experiment by compensating with a 140-fold higher viral inoculum (700 FFU) in *AeSyntenin*-silenced mosquitoes (Fig. 4k). Although delivered intrathoracically, the higher inoculum remains lower than the ∼1,000 infectious particles quantified in whole mosquitoes three days after infectious blood feeding when viruses escape from the midgut^31^. This approach restored infection, yielding comparable levels of gRNA, sfRNA, and sfRNA:gRNA ratios, as well as infectious particles in salivary glands (Fig. 4l-o) and whole mosquitoes (Fig. S21), irrespective of AeSyntenin status. As observed previously (Fig. 4h), salivary gRNA levels remained unaffected (Fig. 4p), whereas salivary sfRNA levels were reduced upon AeSyntenin depletion (Fig. 4q), resulting in a significantly decreased sfRNA:gRNA ratio (Fig. 4r). Supporting a lack of AeSyntenin involvement in viral particle secretion, AeSyntenin depletion had no impact on the quantity of infectious particles in saliva (Fig. 4s). Altogether, these results demonstrate that AeSyntenin plays a key role in secretion of salivary sfRNA.

### AeSyntenin in salivary glands is required for EV production and sfRNA secretion in saliva

To further investigate AeSyntenin function, we genetically abrogated its expression in mosquitoes’ salivary glands. Female mosquitoes expressing Cas9 under a salivary gland-specific promoter^32^ were crossed with male mosquitoes carrying multiple guide RNAs under ubiquitous promoters (Fig. 5a). We generated three female lines - each derived from a different gene insertion event and labeled with different fluorophores (i.e., RFP, EGFP and YFP) - to cross with the same male line, yielding three different progenies. Compared to the maternal Cas9 lines, AeSyntenin levels were successfully reduced in salivary glands but not in carcasses (Fig. 5b,c; Fig. S22a-e). Partial AeSyntenin deletion may result from the promoter’s restricted expression to the distal lateral lobes of salivary glands^32^. TEM revealed that salivary gland-specific AeSyntenin deletion significantly lowered EV concentrations in saliva, as estimated by EV counts per image area (Fig. 5d-f; Fig. S23, S24a). Notably, EVs in the 30–40 nm range were selectively depleted (Fig. 5g,h; Fig. S24b,c).

**Fig. 5.**
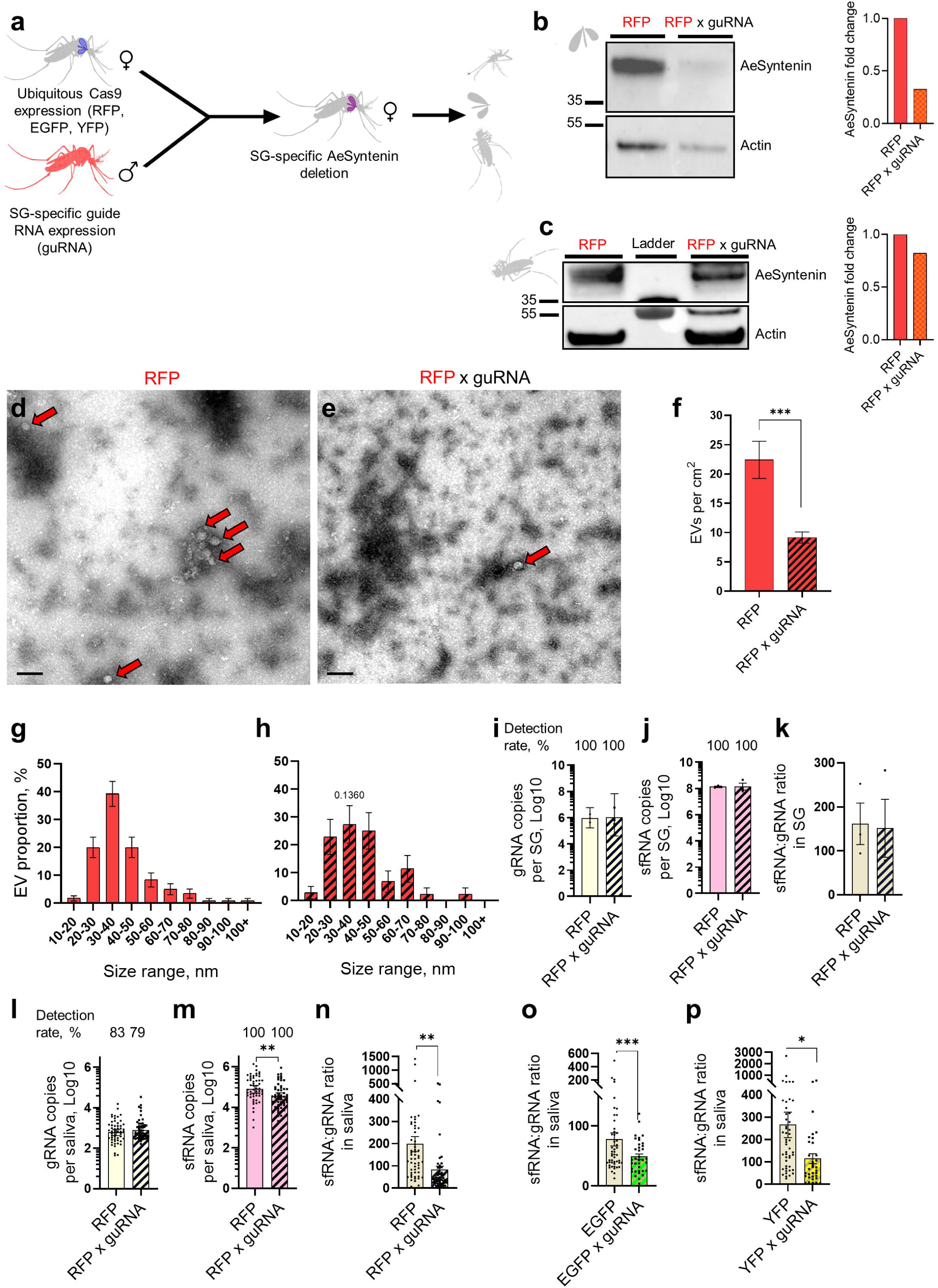
AeSyntenin genetic deletion in salivary glands reduces EVs and sfRNA in saliva. **a** Production of SG-specific AeSyntenin deletion in mosquitoes. **b,c** AeSyntenin detection in SG (b) and carcass (c) from control RFP and RFP x guide RNA (guRNA) progeny. Bar graphs show AeSyntenin fold change normalized to Actin. Each lane corresponds to one replicate. **d-f** Representative TEM pictures of saliva from RFP (d) and RFP x guRNA progeny (e), and EVs per cm² of pictures (f). Scale bar, 100 nm. Red arrows indicate EVs. EV concentration was calculated from 34 and 35 pictures for RFP and RFP x guRNA, respectively. **g,h** Distribution of EVs per size range in saliva from RFP (g) and RFP x guRNA progeny (h). N EVs for g, 120; and for h, 44. **i-k** Levels of gRNA (i), sfRNA (j) and sfRNA:gRNA ratio (k) in SG from infected RFP and RFP x guRNA progeny. **l-n** Levels of gRNA (l), sfRNA (m) and sfRNA:gRNA ratio (n) in saliva from infected RFP and RFP x guRNA progeny. **o,p** Levels of sfRNA:gRNA ratio in saliva from infected EGFP and EGFP x guRNA progeny (o) and YFP and YFP x guRNA progeny (p). f,k,n,o,p Bars represent mean ± s.e.m. and dots indicate repeats. g,h Bars show proportion ± SD. i,j,l,m Bars represent geometric mean ± 95% C.I. and dots indicate repeats. f,m-p *, p < 0.05; *, p < 0.01; ***, p < 0.001 as determined by T-test. h *, p < 0.05 as determined by Z test with same size range in RFP (g).

To evaluate the impact of salivary gland-specific AeSyntenin deletion on sfRNA salivary secretion, we infected transgenic mosquitoes by injecting the high DENV inoculum as previously and quantified gRNA and sfRNA 10 days post-infection. Unlike systemic RNAi, tissue-specific genetic deletion of AeSyntenin did not affect survival (Fig. S25a-c), gRNA or sfRNA levels in salivary glands (Fig. 5i,j; Fig. S26a-d), resulting in comparable sfRNA:gRNA ratios across Cas9 parental lines and most progeny (Fig. 5k; Fig. S26e) – with the exception of one progeny (derived from the YFP Cas9 line) which had a reduced infection and increased sfRNA:gRNA ratio (Fig. S26f), highlighting variability among transgenic lines. In saliva, infection levels remained unchanged in two progeny (Fig. 5l; Fig. S26h) and moderately increased in one (Fig. S26g), correlating with slightly higher salivary gland infection (p = 0.07; Fig. S26a). Importantly, despite progeny variation, most lines exhibited lower sfRNA levels (Fig. 5m; Fig. S26i,j) and all had reduced sfRNA:gRNA ratios (Fig. 5n-p) in saliva. Together, these findings demonstrate that AeSyntenin in salivary glands specifically contributes to EV-associated sfRNA secretion in saliva.

## Discussion

Our study elucidates the first mechanism for selective loading and secretion of viral RNA into exosomes. First, we demonstrate that infection-induced secretion of salivary EVs is partially dependent on the production of sfRNA. Second, we confirm that sfRNA actively promotes EV secretion by showing that sfRNA alone is sufficient to drive its own packaging into small EVs. Third, we provide evidence that sfRNA stem-loops specifically bind both intracellularly and extracellularly to AeSyntenin, which we identified as a key mediator of small EV production in mosquito saliva. Finally, we establish that AeSyntenin in salivary glands is required for the secretion of sfRNA-containing exosomes in saliva. Collectively, these findings support a model in which sfRNA interacts with AeSyntenin to mediate its loading into exosomes, inducing their secretion in mosquito saliva.

Although mechanisms underlying viral RNA loading into EVs remain poorly defined, several pathways have been proposed for the selective incorporation of host RNAs into EVs^33,34^. One well-supported model involves RNA-binding proteins (RBPs) that recognize specific RNA motifs and subsequently interact with EV biogenesis proteins to mediate cargo sorting^35,36^. The involvement of RBPs is supported by the detection of multiple RBPs within EVs, along with the enrichment of their inferred RNA targets. For example, hnRNPA2B1, which contains four RNA-binding domains,^37^ and the La protein^38^ have been shown to bind specific microRNAs and facilitate their inclusion into EVs. In addition, ALG-2-interacting protein X (ALIX)**—**a key regulator of EV biogenesis—interacts with RBPs such as Dicer-2 to control the selective loading of miRNAs^39^. Alternatively, direct electrostatic interactions between RNA and membrane lipids^40^ or indirect interactions through lipid-binding RBPs^41^ can drive RNA incorporation within EV membranes. In this study, we discover that the interaction between sfRNA and AeSyntenin drives sfRNA secretion into EVs. This represents an unexpected function for AeSyntenin. The absence of exomere and supermere proteins^42^ among the sfRNA-interacting proteins (Dataset S1) does not support sfRNA packaging within these small non-vesicular particles. Based on the small size of AeSyntenin-dependent EVs and the structural and functional homologies with human Syntenin-1^9^, we suggest that the sfRNA-containing EVs are exosomes. While our previous work identified sfRNA-interacting RBPs using Aag2 mosquito cell lysates^19^, the AeSyntenin co-IP in the current study did not yield RBPs. This suggests that AeSyntenin-mediated loading of sfRNA into EVs occurs independently of RBPs. Supported by the capacity of Syntenin-1 to load host and viral proteins into exosomes^43,44^, our results show that viral RNA harnesses EV biogenesis proteins for secretion.

Overall, we used a mosquito vector and a global viral pathogen as models to delineate the first mechanism of viral RNA loading into exosomes. We expanded these findings to two other Orthoflaviviruses, suggesting a conserved function of AeSyntenin in viral RNA loading. Leveraging homology in EV biogenesis, this knowledge could help understand how other virus family exploit EVs in multiple animals. In the context of DENV transmission, our results suggest that AeSyntenin deficiency would reduce sfRNA-mediated immune inhibition at the bite site^19,20^.

## Limitations of the study

We leveraged our expertise in mosquito-borne viral diseases to study EV loading. However, our study system suffers from several technical limitations. The nanoliter-size volumes of mosquito saliva preclude the use of biochemical methods usually applied for EV isolation. Additionally, although we generated and previously characterized^24^ a mosquito-specific antibody against EV markers, we did not have access to multiple EV-targeted antibodies, reducing our capacity to characterize mosquito EVs. Moreover, sfRNA sequence identity with gRNA did not allow its direct quantification, and instead we relied on a well-supported^19,20,22^ indirect subtractive sfRNA quantification.

## Acknowledgements

We are grateful to the VectoPole team in Montpellier, particularly Bethsabée Scheid and Carole Ginibre for providing mosquito eggs. We thank Sarah Debaveye for help with generating virus constructs. The authors acknowledge the ISO 9001 certified IRD i-Trop HPC (member of the South Green Platform) at IRD Montpellier for providing HPC resources that have contributed to the research results reported within this paper (https://bioinfo.ird.fr/, http://www.southgreen.fr).

## Material and methods

### Cells and viruses

*Aedes aegypti* Aag2, *Aedes albopictus* C6/36 (ATCC - CRL1660) and Hamster kidney BHK-21 cells (ATCC – CL10) were used. All experiments on mosquito cells were performed with Aag2, whereas C6/36 was exclusively used for virus amplification. More information available in SI Appendix, SI methods.

DENV-2 New Guinea C strain (NGC) was obtained from ATCC (VR-1584). ΔPk1 infectious clone was generated by modifying NGC DENV2 infectious molecular clone pShuttle-DV2/mCherry^45^ with yellow fluorescent protein (mCitrine) using standard molecular biology techniques, i.e. side-directed PCR mutagenesis (KapaHF HotStart ReadyMix, Roche) and homologous recombination in yeast^46^. To abolish sfRNA expression in ΔPk1 infectious clone, a three-nt base exchange was introduced in the DENV2 3’-UTR cDNA impairing the intramolecular base-paring between the top of stem-loop (SL I) and its proximal downstream RNA required for the formation of pseudoknot structure 1 (Pk1)^21^. Recombinant virus was rescued by plasmid DNA transfection of BHK21 cells using TransIT-LT1 transfection reagent (MirusBio), amplified by passage in C6/36 mosquito cells and quantified by plaque titration as previously described^46^. The mCherry non-mutated sfRNA NGC infectious clone was similarly produced from the initial infectious molecular clone^45^ and used as control for ΔPk1 infectious clone.

WNV strain IS-98-ST1 (or Stork 98) was isolated from a stork in Israel in 1998 and obtained from Dr. Philippe Desprès, Centre de Ressources Biologiques, Institut Pasteur, Paris^47^. ZIKV strain H/PF13 was collected from human serum in French Polynesia in 2013 and obtained from the European Virus Archive-Global (EVAg)^48^.

### Mosquitoes

*Aedes aegypti* mosquitoes from the Bora Bora colony^49^ were used. More information available in SI Appendix, SI methods.

### Cell infection

For IP, 1.8 x 10^7^ Aag2 cells were incubated with DENV, WNV and ZIKV at a MOI of 1 in 5 ml of FBS-free cell media for 1 h. For WB and RT-qPCR analyses, 2.5 x 10^5^ Aag2 cells were incubated with DENV at a MOI of 0.05 or 1 in 150 µl of FBS-free cell media for 1 h. For NTA, 8 x 10^6^ Aag2 cells were incubated with DENV at a MOI of 0.05 or 1 for 1 h in 2 ml of FBS-free cell media for 1 h. After removing the inoculum, cells were incubated with complete media containing 2% EV-free FBS for 72 h.

### Quantification of gRNA, sfRNA, ΔPk1 sfRNA and Ctl. RNA

Total RNA was extracted using E.Z.N.A. Total RNA Kit I and eluted in DEPC-treated water. For DENV, gRNA, non-mutated sfRNA/3’UTR, Pk1 mutated sfRNA/3’UTR and Ctl. RNA were quantified by RT-qPCR using the iTaq SYBR GREEN One-Step Kit (Bio-Rad) with primers detailed in Table S1. For WNV and ZIKV, gRNA and sfRNA were quantified with optimized RT-qPCR protocols, as detailed^20^. Absolute quantification was achieved using standard equation. More information available in SI Appendix, SI methods.

SfRNA copies were calculated by subtracting gRNA copies (quantified using primers targeting the envelop) from the combined sfRNA and 3’UTR copy numbers (quantified using sfRNA/3’UTR primers), as previously developed^20,22^. For samples containing detectable levels of sfRNA, sfRNA:gRNA ratio was calculated by dividing sfRNA to gRNA copy numbers. Detection for gRNA was calculated by dividing the number of samples with detectable amount of gRNA over the total number of samples analysed. Detection rate for sfRNA was calculated by dividing the number of samples with detectable amount of sfRNA over the number of samples with gRNA.

### Western Blot (WB)

WB analyses were conducted on eluates from RNA affinity chromatography and IP, and on lysates from Aag2 cells, cell media, whole mosquitoes and salivary glands. For protein blotting, antibodies included 1:500 rabbit anti-human CD63 (Ab134045,clone EPR5702, Abcam), 1:500 mouse anti-human pan-actin (MA5-11869, clone ACTN05, Thermo Scientific), and 1:500 rabbit custom-made anti-AeSyntenin^50^. Intracellular AeSyntenin and hCD63 signals were normalized to actin level. More information available in SI Appendix, SI methods.

### Nanoparticle Tracking Assay (NTA)

Cell media was diluted by half in PBS and analyzed using a NanoSight NS300 instrument (Malvern) equipped with a 488 nm laser. More information available in SI Appendix, SI methods.

### Infection of mosquitoes

Three-to-five day-old female mosquitoes were intrathoracically microinjected with 5 FFU of wild-type DENV in a single injection of 69 nl using Nanoject II (Drummond). An equivalent volume of RPMI was injected as control. DsRNA-injected female mosquitoes were microinjected with 5 or 700 FFU of wild-type DENV at 4 days post dsRNA injection. Mosquitoes were maintained on a 10% sugar solution for an additional 10 days before analysis.

### Extracellular vesicle (EV) isolation

EV-free FBS was produced by ultracentrifugating heat-inactivated FBS at 100,000 × g for 18 h using an S58-A rotor (k-factor 50, Thermo Scientific) in a Sorvall MX Plus Series Floor Model Micro-Ultracentrifuge (Thermo Scientific). EV-free FBS was collected by leaving approximately 1 ml of residual volume at the bottom of the 8 ml tube, filtered through a 0.22 µm filter (Sartorius), and stored at 4°C. 1.8 x 10^6^ Aag2 cells per T175 flask were maintained in complete medium supplemented with 2% EV-free FBS for 3 days. Cell media was then centrifugated at 3,500 × g for 5 min to remove cell debris. The clarified supernatant was ultracentrifugated at 100,000 × g for 3 h to obtain an EV-concentrated pellet. For NTA, the EV-concentrated pellet was washed in PBS and ultracentrifugated one more time at 100,000 x g for 3h before resuspension in PBS.

### Collection of mosquito saliva and tissues

Mosquitoes were starved for 24h, cold-anesthetized, and immobilized by removing their wings and legs. The proboscis of each mosquito was individually inserted into a 20 µl pipette tip containing 10 µl of pre-warmed DMEM supplemented with 0.5 mM ATP (Invitrogen) and 1% Erioglaucine (25 mM, Sigma-Aldrich) for 1 h^20^. Mosquitoes with blue-colored abdomens, indicating ingestion of the dye, were considered as having salivated. For TEM observation, saliva was collected in bulk. Hundreds of mosquitoes were let to feed on artificial feeding system (Hemotek) containing 3 ml of DMEM supplemented with 0.5 mM ATP and 1% Erioglaucine covered with a silicone membrane for 2 h. The DMEM media was then precleared by centrifugation at 1,500 g for 10 min at 4°C, ultracentrifugated at 100,000 g for 3 h at 4°C and resuspended in 30 μl of PBS. Number of salivating mosquitoes was estimated by counting the number of blue-abdomen mosquitoes, indicating salivation and quantities of EVs were normalized per mosquitoes between infected and non-infected samples. For RNA quantification, individual saliva samples were transferred into 200 µl of TRK lysis buffer. For FFU assays, pooled saliva samples from 10 mosquitoes was combined with 100 µl of RPMI medium supplemented with 1% P/S and 1% antibiotic-antimycotic solution (Gibco). Following saliva collection, salivary glands were dissected using fine forceps. For RNA quantification, 5 salivary glands were pooled in 200 µl of TRK lysis buffer. For FFU assays, 5 salivary glands were pooled in 150 µl of RPMI 1% P/S and 1% antibiotic-antimycotic. For WB, 5 or 10 salivary glands were pooled in 150 µl of RIPA buffer IV. Remaining carcasses were collected and processed similarly by pools of five. Tissues and saliva samples were homogenized using silica beads with a BeadBeater homogenizer. RNA was extracted using the EZNA Total RNA Kit I. For FFU assays, homogenized tissues and saliva were sterilized by filtration through a 0.22 µm filter (Millex-GV). For WB, lysates from salivary glands and carcasses were pre-cleared by centrifugation at 3,500 × g for 10 min.

### Transmission electron microscopy (TEM)

For uninfected and infected EV saliva, 3 µl of ultracentrifugated EV solution, and for EV saliva from transgenic mosquitoes, 7 µl of ultracentrifugated salivary EV solution was analysed. More information available in SI Appendix, SI methods.

### Production of sfRNA, ΔPk1 sfRNA and control RNA (Ctl. RNA)

sfRNA, ΔPk1 sfRNA and Ctl. RNA, corresponding to a DENV NS2 fragment^27^, were amplified from viral cDNA using T7 promoter-tagged forward primers (Table S1), transcribed using MEGAscript T7 Kit, purified with the Total RNA Kit I, monophosphorylated with RNA 5’ polyphosphatase (Biosearch Technologies), re-purified with Total RNA kit I, quantified as above and folded by incubating at 95°C for 5 min and gradually cooling to 4°C for folding. More information available in SI Appendix, SI methods.

### Transfection of sfRNA, ΔPk1 sfRNA and Ctl. RNA

10^10^, 10^14^ and 10^12^ copies of monophosphorylated folded sfRNA, ΔPk1 sfRNA and Ctl. RNA, respectively, were transfected for 1 h using TransIT mRNA Transfection Kit (Euromedex) into 2.5 x 10^5^ Aag2 cells for WB analysis or into 1.2 x 10^6^ Aag2 cells for NTA analysis. Difference between sfRNA, ΔPk1 sfRNA and Ctl. RNA amounts was to compensate for a lower transfection efficiency of Ctl. RNA and ΔPk1 sfRNA (Fig. S7a). Transfection media was removed and cells were washed three times with pre-warmed RPMI medium. After three days of incubation with 2% EV-free FBS 1% P/S 1X NEAA RPMI, media was collected, pre-cleared by centrifugation at 3,500 x g for 10 min and lysed in RIPA buffer IV. For NTA analysis, pre-cleared cell media was ultracentrifugated at 100,000 × g for 3 h, the pellet washed, ultracentrifugated at 100,000 x g for 3 h, and resuspended in 500 µl PBS. Aag2 cells were lysed in TRK lysis buffer (EZNA Total RNA Kit I) for RNA quantification or in 100 µl of RIPA buffer IV for WB.

### RNAse resistance assay

Three days post sfRNA and Ctl. RNA transfection, 1 ml of cell media was pre-cleared at 3,500 x g for 10 min, divided into equal volumes of 4 x 200 µl and treated with different conditions. Samples were treated with 2.5 µl of 0.1 % triton X 100 (Sigma) or PBS as control and incubated at 4°C at 30 min. Then, 5 µl of RNase A/T1 (Thermofisher Scientific) or PBS as control was added and samples were incubated at 37°C for 30 min before RNA extraction using E.Z.N.A. Total RNA extraction Kit I. Twelve replicates per condition were performed in four series of three replicates each. To account for between-experiment variability, fold changes were normalized over the mean of the control replicates within each experiment.

### RNA affinity chromatography

RNA affinity chromatography was performed as previously described^27^ with minor modifications. Beads bound with DENV sfRNA or a same-size control RNA fragment were incubated with lyzed Aag2 cells or isolated EVs. Protein eluates were analysed by MS and WB. More information available in SI Appendix, SI methods.

### Mass spectrometry (MS)

Eluates from RNA affinity chromatography were analyzed using the facilities of the Montpellier Proteomics Platform (PPM, BioCampus Montpellier), a member of the national Proteomics French Infrastructure (ProFI UAR 2048). More information available in SI Appendix, SI methods.

### Immunoprecipitation (IP) and co-immunoprecipitation (co-IP)

Lysed infected Aag2 cells and EVs from infected Aag2 cells were subjected to IP and co-IP with using beads decorated with anti-AeSyntenin antibody^50^. Beads decorated with anti-rabbit IgG antibody (Invitrogen) were used as control. More information available in SI Appendix, SI methods.

Affinity of the anti-AeSyntenin antibody for sfRNA was assessed by coupling SureBeads Protein G Magnetic Beads with either anti-rabbit IgG or rabbit anti-AeSyntenin antibody as above and incubating them with 10¹ copies of *in vitro*-transcribed sfRNA for 2 h at 4°C on a disk rotator. Elution for RNA extraction was performed as above.

### RoseTTAFoldNA prediction

Interaction predictions between the 3D structures of SL1, SL2, SL1-SL2, DB1, DB2, D1-D2, 3’SL for DENV, and SL1, SL2, 3’SL for WNV and ZIKV, and the AeSyntenin was performed on RoseTTAFoldNA version 0.2^28^. Molecular visualization was performed with Molstar^51^.

### Quantification of mRNA

Total RNA was extracted using the EZNA Total RNA Kit I, cleared of genomic DNA using DNAse following protocol from gDNA Clear cDNA Synthesis kit (Biorad), quantified with a NanoDrop, normalized, and reverse transcribed using iScript gDNA Clear cDNA Synthesis Kit (Bio-Rad). *AeSyntenin* mRNA was quantified using qPCR. *Actin* and *RPS7* mRNA were quantified as housekeeping genes. More information available in SI Appendix, SI methods.

### RNAi-mediated silencing

Target sequences were amplified from Aag2 cDNA using GoTaq Master Mix (Promega) and 400 nM T7-flanked primer pairs (Table S1), transcribed overnight with the MEGAscript T7 Kit, extracted with the EZNA Total RNA Kit I, adjusted to 14 µg RNA/µl, and annealed by heating at 95°C for 5 min followed by gradual cooling. Negative control dsRNA targeting LacZ^52^ was similarly produced (Table S1). Four-day-old female mosquitoes were cold-anesthetized and injected with 138 nl of dsRNA using Nanoject II microinjector (Drummond Scientific). Mosquitoes were maintained on a 10 % sugar solution. More information available in SI Appendix, SI methods.

### Focus forming unit (FFU) assay

FFU was quantified in BHK-21 cells. More information available in SI Appendix, SI methods.

### Salivary gland-specific abrogation of AeSyntenin

Golden Gate Cloning was used to construct a first piggyBac transgenesis vector with Cas9 under the control of the 30Ka promoter^32^. In a second piggyBac transgenesis vector, different guide RNAs targeting *AeSyntenin* were placed under the control of four different U6 promoters: protospacer GACGAGTCCTTAACGATGG(CGG) under control of U6 promoter AAEL017774; GTCACCAACGGAATTCGAG(AGG) under U6 AAEL017763; GATACTTCCGGTTGGTAAG(CGG) under U6 AAEL017774, and GGGATTGGAGCTCAGCCGTG(AGG) under U6 AAEL017702. The sequences of the transgenesis plasmids are provided in Fig. S27 and DataSet S3 and S4. Transgenic mosquitoes were obtained by injecting embryos of the Liverpool strain with 240 ng/l of piggyBac plasmid mixed with 80 ng/µl of transposase helper plasmid^53^. Injected mosquitoes reaching adulthood and showing transient expression of the fluorescence marker were backcrossed to wild-types. In the next generation (G1), transgenic larvae were selected based on expression of the plasmid’s fluorescence markers^54^. Single G1 females were crossed to wild-type males. Transgenes showing about 50% inheritance, indicative of probable single insertions, were retained and the G2 positives of each progeny were self-crossed to establish lines. In the G3 generation, homozygous larvae were selected by their stronger fluorescence to establish stable lines. Three lines with Cas9 expression were produced each with a different selection fluorescent marker (i.e., RFP, EGFP, YFP), while one line with guRNA was produced. From the guide RNA lines, seven homozygous males were mass-crossed with twenty virgin Cas9-expressing females. The resulting eggs were allowed to develop for 3 to 5 days before hatching. Individual containers were provided with blood meals. Blood-fed females were then transferred to separate boxes for oviposition. After egg-laying, the egg papers were removed, and the eggs were allowed to mature for at least five days before hatching the cohorts selected for experiments.

### Statistics

Statistical analyses were conducted using Prism v8 (GraphPad). More information available in SI Appendix, SI methods.

