## Supplemental material for "Dengue virus harnesses mosquito Syntenin to load and secrete viral RNA into salivary exosomes"

### ****Material and methods****

Cells and viruses**.** *Aedes aegypti* Aag2 and *Aedes albopictus* C6/36 (ATCC - CRL1660) were grown in Roswell Park Memorial Institute (RPMI) medium (Gibco) and Dulbecco’s Modified Eagle Medium (DMEM) (Gibco), respectively, with 10% heat-inactivated fetal bovine serum (FBS) (Gibco), 1X non-essential amino acids (Gibco), and 1% penicillin/streptomycin (Gibco) at 28°C with 5% CO₂. Hamster kidney BHK-21 cells (ATCC – CL10) were grown in RPMI medium with 5% heat-inactivated FBS and 1% penicillin/streptomycin at 37°C, 5% CO₂. All experiments were performed with Aag2, whereas C6/36 was exclusively used for virus amplification.

Mosquitoes**.** *Aedes aegypti* mosquitoes from the Bora Bora colony^1^ were maintained in BugDorm cages (BugDorm) at 27 ± 1°C, with 70 ± 5% relative humidity, under a 12 h:12 h light:dark cycle, and provided with *ad libitum* access to a 10% sucrose solution. Larvae were reared at 28°C in Milli-Q water supplemented with fish food flakes (TetraMin) until pupation.

Quantification of gRNA, sfRNA, ΔPk1 sfRNA and Ctl. RNA**.** Total RNA was extracted using E.Z.N.A. Total RNA Kit I and eluted in DEPC-treated water. For DENV, gRNA, non-mutated sfRNA/3’UTR, Pk1 mutated sfRNA/3’UTR and Ctl. RNA were quantified by RT-qPCR using the iTaq SYBR GREEN One-Step Kit (Bio-Rad) with primers detailed in Table S1. A total reaction volume of 10 µl containing 0.3 µl of 300 nM of forward primer, 0.3 µl of 300 nM reverse primer, 5 µl of iScript master mix, 2.275 µl of water nuclease-free water and 2 µl of RNA extract was amplified using LightCycler 96 (Roche) with the following thermal profile: 50°C for 15 min, 95°C for 1 min, followed by 40 cycles of 95°C for 10 sec and 60°C for 15 sec, with a final melting curve analysis.

For WNV, gRNA was quantified by RT-qPCR in LighCycler 96 thermocycler (Roche) with the following thermal profile: 50°C for 10 min, 95°C for 2 min, and 40 cycles at 95°C for 15s, 60°C for 15s and 72°C for 20s. SfRNA was quantified with the following thermal profile : 50°C for 20 min, 95°C for 1 min, and 40 cycles at 95°C for 10s and 60°C for 25s with a total reaction volume of 10 µl containing 0.3 µl of 300 nM of forward primer, 0.3 µl of 300 nM reverse primer, 5 µl of iScript master mix, 2.275 µl of water nuclease-free water and 2 µl of RNA extract, as detailed^2^.

For ZIKV, gRNA was quantified using iTaq Universal Probes one-step RT-qPCR kit (Bio-Rad) in LighCycler 96 (Roche) with the following thermal profile: 50°C for 15 min, 95°C for 2 min, and 45 cycles at 95°C for 10s and 60°C for 30s, with a total reaction volume of 10 µl containing 0.3 µl of 300 nM of forward primer, 0.3 µl of 300 nM reverse primer, 5 µl of iScript Universal probe master mix, 2.075 µl of water nuclease-free water, 0.2 µl of 200 nM probe and 2 µl of RNA extract. SfRNA was quantified using a two-step RT-qPCR. RT was conducted with the reverse primer and M-MLV enzyme (Promega) with a denaturation/annealing step at 70°C and following manufacturer’s conditions with RT at 37°C. qPCR was then performed in AriaMx thermocycler (Agilent) using EvaGreen qPCR MixPlus (Euromedex) with the following thermal profile: 90°C for 15 min, and 40 cycles at 95°C for 15s, 62°C for 30s and 72°C for 20s, with a total reaction volume of 10 µl containing 0.3 µl of 300 nM of forward primer, 0.3 µl of 300 nM reverse primer, 2 µl of EvaGreen master mix, 5 µl of water nuclease-free water and 2 µl of DNA extract, as detailed^2^.

Absolute quantification was achieved using standard equation. by quantifying qPCR templates using forward T7-tagged primers detailed in Table S1. Templates were transcribed using the MegaScript T7 Kit and purified with the E.Z.N.A. Total RNA Extraction Kit I. RNA concentrations were measured using a NanoDrop spectrophotometer (Thermo Fisher Scientific) and serial dilutions were used to generate absolute standard curves.

Western Blot (WB)**.** WB analyses were conducted on eluates from RNA affinity chromatography and IP, and on lysates from Aag2 cells, cell media, whole mosquitoes and salivary glands. Cell lysate was obtained by adding RIPA buffer IV (Bio Basic), centrifugating the lysate at 20,000 × g for 20 min and collecting the supernatant. Cell media was pre-cleared by centrifugation at 3,500 x g for 10 min and lysed in RIPA buffer IV. Mosquito and salivary gland lysates were obtained by homogenizing 5 carcasses and 10 salivary glands in 250 or 150 µl of RIPA buffer IV, respectively, with silica beads (BioSpec) using a BeadBeater homogenizer (MP Biomedicals) and collecting supernatants after centrifugation at 14,000 rpm for 10 min at 4°C. Proteins from eluates and lysates were quantified using Qubit protein assay (ThermoFisherScientific). Homogenized quantities of proteins were diluted in reducing Laemmli SDS buffer, heated to 95°C for 10 min, separated on NuPAGE 4-12% acrylamide gel (Invitrogen) and transferred onto a 0.2 µm nitrocellulose membrane (Bio-Rad) using Trans-Blot turbo Transfer System (Bio-Rad). Membranes were washed with 0.1% Tween 20 in PBS, blocked with 5% milk (Régilait) for 1 h at RT, incubated overnight at 4°C with primary antibodies diluted in 0.1% Tween 20 in PBS. Antibodies included 1:500 rabbit anti-human CD63 (Ab134045,clone EPR5702, Abcam), 1:500 mouse anti-human pan-actin (MA5-11869, clone ACTN05, Thermo Scientific), and 1:500 rabbit custom-made anti-AeSyntenin^3^. Secondary antibody staining was performed with 1:2,000 HRP-conjugated anti-rabbit IgG (7074, Cell Signaling Technology) or 1:2,000 HRP-conjugated anti-mouse IgG (7076, Cell Signaling Technology) in 0.1% Tween 20 5% milk in PBS for 1 h at RT. Blots were revealed with SuperSignal West Pico PLUS Chemiluminescent Substrate (Thermo Scientific) or SuperSignal West Femto Chemiluminescent Substrate (Thermo Scientific) using a ChemiDoc MP Imaging System (Bio-Rad) and analyzed with ImageLab Software (Bio-Rad). Blot intensity was estimated with ImageLab. Intracellular AeSyntenin and hCD63 signals were normalized to actin level. For each gel, protein fold changes were calculated relative to the mean value of the control replicates loaded on the same gel, thereby accounting for between-gel variability.

Nanoparticle Tracking Assay (NTA)**.** Cell media was diluted by half in PBS and analyzed using a NanoSight NS300 instrument (Malvern) equipped with a 488 nm laser. The capture settings were as follows: camera type sCMOS, laser type Blue488, camera level 12, slider shutter 1206, slider gain 366, frame rate (FPS) 25.0, number of frames 1498, temperature 25.0 °C, viscosity 0.9 cP (for water), and syringe pump speed 40. Five 1-minute videos were recorded, and particle analysis was conducted using NanoSight NTA software version 3.4 (build 3.4.4, Malvern). NTA analyses were performed at the Charles Gerhardt Institute, Montpellier, France.

Transmission electron microscopy (TEM). For uninfected and infected EV saliva, 3 µl of ultracentrifugated EV solution was absorbed on a glow-discharged 300 mesh formvar carbon-coated grid (Electron Microscopy Science) and stained with 1% uranyl acetate in two steps. Grids were observed using a JEOL 1400 Flash transmission electron microscope at 120kV. Micrographs were recorded using a One View camera (Gatan Inc.). Experiment was performed at the Center of Structural Biology of Montpellier, CNRS, France. For EV saliva from transgenic mosquitoes, 7 µl of ultracentrifugated salivary EV solution was loaded onto a formvar-coated copper grid 100 mesh (EMS FCF100-Cu-50), left to settle for 2 min, and dried with a blotting paper. The grid was stained with 1% uranyl acetate and analyzed using a TEMF20 operating at 120 KV (Tecnai) with a Veleta numeric camera (Olympus) at the Neuroscience institute of Montpellier, INSERM U1298, France.

Production of sfRNA, ΔPk1 sfRNA and control RNA (Ctl. RNA)**.** sfRNA, ΔPk1 sfRNA and Ctl. RNA, corresponding to a DENV NS2 fragment^4^, were amplified from viral cDNA using T7 promoter-tagged forward primers. As template, we used NGC virus for sfRNA and the ΔPk1 infectious clone for ΔPk1 sfRNA. sfRNA, ΔPk1 sfRNA and Ctl. RNA primers are detailed in Table S1. PCR products were size-validated by agarose gel electrophoresis, purified using the QIAquick PCR Purification Kit, and *in vitro* transcribed using MEGAscript T7 Kit. Transcripts were purified with the E.Z.N.A. Total RNA Kit I and quantified via NanoDrop. RNA fragments were treated with RNA 5’ polyphosphatase (Biosearch Technologies), and re-purified with Total RNA kit I and quantified as above. Samples were incubated at 95 °C for 5 min and gradually cooled to 4 °C for folding.

RNA affinity chromatography**.** RNA affinity chromatography was performed as previously described^4^ with minor modifications. 1 ml of NHS Mag Sepharose bead (Cytiva) flurry was washed four times with 1 ml of 1 mM HCl (Carlo Erba) and incubated in 950 µl of coupling buffer [0.2 M NaHCO₃ (Sigma), 0.5 M NaCl (Sigma), pH 8.3] with 50 µl of 5 mM tobramycin (Sigma) overnight at 4°C under gentle rotation. Tobramycin beads were blocked in 1 ml of blocking buffer [20 mM Tris pH 7.0 (Sigma), 1 mM CaCl_2_ (Sigma), 1 mM MgCl_2_ (Sigma), 300 mM KCl (Sigma), 0.1 mg/ml of tRNA (Ambion), 0.5 mg/ml BSA (Sigma), 0.2 mM Dithiothreitol (Biorad), 0.01% NP-40 (MP Biomedical)] for 1 h at 4°C, washed three times with PBS and resuspended in 500 µl of PBS. DENV sfRNA and a same-size control RNA fragment both flanked by a tobramycin aptamer sequence were amplified from plasmids^4,5^ using T7-tagged forward primers detailed in Table S1. ∆SL sfRNA, ∆DB sfRNA and ∆3’SL sfRNA were amplified from the plasmid and linked to the aptamer sequence using splice overlap extension PCR, with the following program: 95°C for 2 min, and 30 cycles at 95°C for 30s, 55°C for 30s and 72°C for 30s, and 72°C for 10min. After size validation on agarose gel, PCR products were purified using the QIAquick PCR Purification Kit (Qiagen) and transcribed using MEGAscript T7 Kit (Thermo Fisher). Transcribed RNA was extracted with EZNA Total RNA Kit I (Omega) and quantified using Nanodrop. 30 µg of *in vitro*-transcribed RNA in 200 µl of RNA binding buffer (20 mM Tris pH 7.0, 1 mM CaCl₂, 1 mM MgCl₂, 75 mM NaCl, 145 mM KCl, 0.1 mg/ml yeast tRNA, 0.2 mM Dithiothreitol) was heated to 95°C for 5 min and slowly cooled to RT to enable RNA folding. The RNA solution was incubated with 30 µl of tobramycin-coupled beads for 1 h at 4°C under end-over-end rotation. Beads were washed once with 250 µl of RNA washing buffer (20 mM Tris pH 7.0, 1 mM CaCl₂, 1 mM MgCl₂, 75 mM NaCl, 145 mM KCl, 0.1% NP-40, 0.2 mM Dithiothreitol).

1.8 x 10^7^ uninfected Aag2 cells or isolated EVs were lysed in 1 ml and 500 µl of lysis buffer [50 mM Tris pH 7.5 (Sigma), 150 mM NaCl, 1% NP-40, 1 X protease and phosphatase inhibitor (Sigma)], respectively. 417 µl of cell lysate in 500 µl of 1 X modified protein buffer (2 mM CaCl_2_, 2 mM MgCl_2_, 290 mM KCl, 4 mM Dithiothreitol) or 210 µl of EV lysate in 300 µl of 1 X modified protein buffer were pre-cleared with 30 µl of tobramycin bead flurry during 1 h at 4 °C and then incubated with beads from 30 µl of RNA-bead flurry for 2 h at 4°C with end-over-end rotation. Beads were washed four times in 500 µl of 145 mM protein washing buffer (20 mM Tris pH 7.0, 1 mM CaCl₂, 1 mM MgCl₂, 75 mM NaCl, 145 mM KCl, 0.5% NP-40). Elution for MS analysis was performed by incubating beads in 20 µl of elution buffer (20 mM Tris pH 7.0, 1 mM CaCl₂, 3 mM MgCl₂, 145 mM KCl, 5 mM tobramycin, 0.2 mM Dithiothreitol) for 5 min on ice and elution for WB analyses was performed in 25 µl of Laemmli SDS buffer (Thermo Scientific) and heated to 95 °C for 5 min.

Mass spectrometry (MS)**.** Eluates from RNA affinity chromatography were analyzed using the facilities of the Montpellier Proteomics Platform (PPM, BioCampus Montpellier), a member of the national Proteomics French Infrastructure (ProFI UAR 2048). Protein digestion was performed on S-TrapTM micro columns (Protifi) following the manufacturer’s instructions. In brief, protein extracts were diluted in 40 µL of final 5% SDS / 50 mM triethylammonium bicarbonate (TEAB), reduced with 20 mM dithiothreitol (DTT) and held for 10 min at 95°C. Samples were cooled to room temperature and alkylated with 40 mM iodoacetamide (IAA) for 30 min at room temperature in dark. Samples were acidified with phosphoric acid at final concentration of 1.2% and diluted 6 times in S-Trap binding buffer (90% methanol / 100 mM TEAB). The resulting protein suspension was transferred to the S-Trap filter via centrifugation at 4000g for 1 min. Trapped proteins were washed three times with 150 µL S-Trap binding buffer. 1 µg of trypsin (Trypsin Glod, Promega) in 50 mM TEAB were added to the filter surface and incubated for 2 hours at 47°C. Tryptic peptides were eluted sequentially with 40 µl of 50 mM TEAB, 50 mM TEAB / 0.2% aqueous formic acid, and then with 50% of acetonitrinile (ACN) *via* centrifugation at 4000g. Eluted peptides were vacuum-dried. Peptide samples were reconstituted in 10 µl of buffer A (0.1% formic acid) and injected into nano-flow HPLC system (Ultimate 3000 RSLC, Thermo Fisher Scientific) coupled to a mass spectrometer equipped with a nanospray source (Q-Exactive HF, Thermo Fisher Scientific). Peptides were separated on a capillary column (0.075 mm × 500 mm, Acclaim PepMap 100, reverse-phase C18, NanoViper, Thermo Fisher Scientific) using a gradient of 2-40% B over 120 minutes with A = 0.1% formic acid and B = 0.1% formic acid in 80% acetonitrile, at a flow rate of 300 nl/min (total runtime: 150 min). Spectra were recorded using Xcalibur 4.1 software (Thermo Fisher Scientific). Raw spectra were processed using the MaxQuant environment (Cox and Mann, 2008, v1.6.10.43) and Andromeda for database search with Label-Free Quantification (LFQ), match between runs and the iBAQ algorithm enabled^6^. The MS/MS spectra were matched against the UniProt Reference proteome of Aedes_aegypti (Proteome ID UP000008820 ; <https://www.uniprot.org/>), VectorBase-52_AaegyptiLVP_AGWG_AnnotatedProteins and a homemade contaminant database. Different release versions were used, depending on the date of analysis. Enzyme specificity was set to trypsin/P, and the search included cysteine carbamidomethylation as a fixed modification and oxidation of methionine, and acetylation (protein N-term) as variable modifications. Up to two missed cleavages were allowed for protease digestion. FDR was set at 0.01 for peptides and proteins and the minimal peptide length at 7. A representative protein ID in each protein group was automatically selected using an in-house developed bioinformatics tool (Leading tool v3.4) as the best annotated protein in UniProtKB (reviewed entries rather than automatic ones)^7^. Proteins with more than 1.5 fold enrichment with LFQ ratio and 2 fold enrichment with MS/MS pseudoRatio were considered significantly enriched. The mass spectrometry proteomics data have been deposited to the ProteomeXchange Consortium via the PRIDE^8^ partner repository with the dataset identifier PXD065152.

Eluates from AeSyntenin co-IP were analyzed by the Proteomic Platform of Strasbourg-Esplanade (PPSE) at the Institut de Biologie Moléculaire et Cellulaire (IBMC). Briefly, proteins were first precipitated overnight using 0.1 M ammonium acetate in 100% methanol. After centrifugation at 12,000 x g and 4°C for 15 min, the pellets were washed twice with 0.1 M ammonium acetate in 80% methanol. This precipitation step was repeated once more. Then, proteins were dried using a Speed-Vac concentrator. The dried pellets were resuspended in 100 µL of 50 mM ammonium bicarbonate, reduced with 5 mM dithiothreitol at 95°C for 10 minutes, and alkylated with 10 mM iodoacetamide at room temperature for 30 min in the dark. Samples were digested with 150 ng sequencing-grade trypsin (Promega) at 37°C overnight. Peptides were resuspended in 0.1% formic acid, and 3/64 of the sample was loaded for nLC-MS/MS analysis. Samples were analyzed using a timsTOF Pro 2 coupled with a NanoElute 2 (Bruker Daltonik GmbH) in Data-Independent Acquisition and Parallel Accumulation Serial Fragmentation (DIA-PASEF) mode. Peptides were separated on an IonOpticks Aurora Elite column (25 cm × 75 μm, 1.7 μm particle size, and 120 Å pore size; AUR3–15075C18-CSI) with a 45-minute elution gradient. Data were searched against the *Aedes aegypti* database from UniProt reference proteome (release march 2021, 33 998 sequences) using DIANN 1.9 software with a library-free approach keeping most of the default parameters. One missed cleavage and one variable modification were allowed (M-Ox, Ac-Nterm), and match between runs was enabled. Prostar 1.34.5 was used for statistical analyses. Imputation of missing values was based on the 1% detection quantile, and a LIMMA statistical test was used for each comparison. The mass spectrometry proteomics data from the immunoprecipitations have been deposited to the ProteomeXchange Consortium via the PRIDE partner repository with the dataset identifier PXD065851.

Immunoprecipitation (IP) and co-immunoprecipitation (co-IP)**.** Infected Aag2 cells and EVs from infected Aag2 cells were lysed in 500 µl of NT2 buffer [200 mM KCl, 20 mM HEPES pH 7.2, 2% N-dodecyl-β-D-maltoside (Thermofisher Scientific), 1% Igepal (Sigma), 100 U/mL Murine RNase inhibitor (Biolabs)] and supernatant was collected after centrifugation at 13,000 × g for 13 min at 4°C. 100 µl of SureBeads Protein G Magnetic Beads (Bio-Rad) were washed with 500 µl of 0.01% Tween 20 in PBS, blocked with 50 µl of 1% BSA 0.01% Tween 20 in PBS for 10 min at 4°C, washed with PBS and incubated for 1 h at 4°C with 500 µl of 0.01% Tween 20 in PBS containing 2.5 µg of custom-made anti-AeSyntenin antibody^3^ on a disk rotator. Beads were similarly incubated with anti-rabbit IgG antibody (Invitrogen) as control. After three washes with 500 µl of 0.01% Tween 20 in PBS, beads were incubated with homogenized volumes of lysed proteins for 2 h at 4°C on a rotator. 1 % of initial input was kept for analysis by WB as control. After washes, beads diluted in 500 µl of 0.01% Tween 20 in PBS were separated into aliquots of 100 µl for WB analysis and 400 µl for RNA quantification. WB samples were resuspended in 30 µl of Laemmli SDS buffer and warmed at 95°C for 5 min. RNA quantification samples were resuspended in 350 µl of TRK lysis buffer before extraction with EZNA Total RNA extraction Kit I. For co-IP, all beads were resuspended in 50 µl of Laemmli SDS buffer and heated at 95°C for 5 min. Elution was repeated once with 50 µl of Laemmli SDS buffer to obtain a final elution volume of 100 µl.

Quantification of mRNA**.** Total RNA was extracted using the EZNA Total RNA Kit I, cleared of genomic DNA using DNAse following protocol from gDNA Clear cDNA Synthesis kit (Biorad), quantified with a NanoDrop, normalized, and reverse transcribed using iScript gDNA Clear cDNA Synthesis Kit (Bio-Rad). *AeSyntenin* mRNA was quantified in a total reaction of 10 µl, containing 2 µl of Hot FirePol SolisGreen qPCR mix (Euromedex), 0.3 µl of 300 nM forward, 0.3 µl of 300 nM reverse primers (Table S1), 5.4 µl of nuclease-free water and 2 µl of cDNA. qPCR thermal profile consisted of 95°C for 5 min, followed by 40 cycles of 95°C for 15 sec, 60°C for 20 sec, and 72°C for 20 sec, with a final melting curve analysis on a LightCycler 96 (Roche). *Actin* and *RPS7* mRNA were quantified as housekeeping genes and the geometric mean of their Ct values was used for normalization using the delta-delta Ct method.

RNAi-mediated silencing**.** Target sequences were amplified from Aag2 cDNA using GoTaq Master Mix (Promega) and 400 nM T7-flanked primer pairs (Table S1) designed via the E-RNAi website^9^ with the following thermal profile: 95°C for 5 min; 45 cycles of 95°C for 15 sec., 55°C for 15 sec., and 72°C for 45 sec.; 72°C for 10 min. After product size confirmation on agarose gel, PCR products were purified using the QIAquick PCR Purification Kit and transcribed overnight with the MEGAscript T7 Kit. RNA was extracted with the EZNA Total RNA Kit I, adjusted to 14 µg/µl, and annealed by heating at 95°C for 5 min followed by gradual cooling. Negative control dsRNA targeting LacZ^10^ was similarly produced (Table S1). Four-day-old female mosquitoes were cold-anesthetized and injected with 138 nl of dsRNA using Nanoject II microinjector (Drummond Scientific). Mosquitoes were maintained on a 10 % sugar solution.

Focus forming unit (FFU) assay**.** Samples were subjected to a four-fold serial dilution in RPMI medium and 150 µl of each dilution was incubated with 1.5 × 10^5^ BHK-21 cells for 1 h on a rocking platform. After removing the inoculum, cells were overlaid with 500 µl of 1% carboxymethyl cellulose (CMC) (Sigma) in 2% FBS (Gibco) RPMI medium. After 5 days, cells were fixed with 4% paraformaldehyde (VWR) in PBS, permeabilized with 0.3% Triton X-100 (Euromedex), blocked with 1% FBS in PBS, and stained using a 1:200 mouse monoclonal anti-envelope antibody (4G2, provided by Dr Subhash Vasudevan, Duke-NUS Medical School, Singapore) and 1:500 secondary anti-mouse IgG Alexa Fluor 488 (Invitrogen). FFU were counted using the EVOS m5000 microscope (Invitrogen) in three replicates per dilution, and average FFU per sample was calculated per ml.

Statistics. One-tailed T-test was applied to test differences in log-transformed gRNA, sfRNA copy numbers and fold changes, in non-transformed sfRNA:gRNA ratio, AeSyntenin and hCD63 protein quantities. Variations in concentration and proportion of particles within size ranges were tested using ANOVA test with post-hoc FDR adjustment or Z-test. Differences in RNA quantities after RNase assay and in AeSyntenin and hCD63 protein quantities were assessed using ANOVA followed by post-hoc Fisher’s LSD test. Statistical analyses were conducted using Prism v8 (GraphPad).

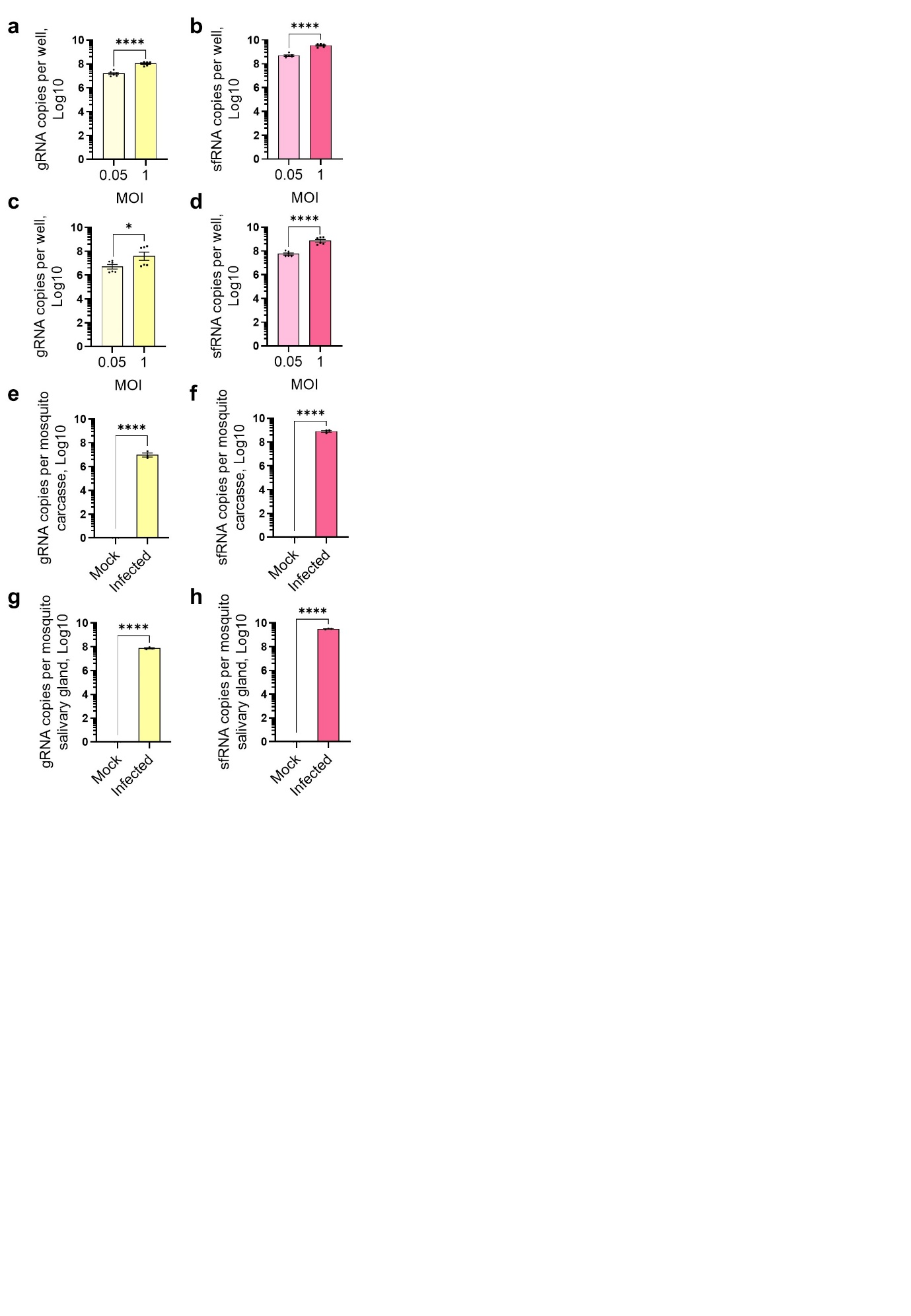

Fig. S1 | Quantification of gRNA and sfRNA in infected and mock-infected Aag2 mosquito cells and mosquitoes, related to Fig. 1. **a,b** Levels of gRNA (a) and sfRNA (b) in cells infected with MOI 0.05 or 1 at 72 hpi. **c,d** Levels of gRNA (c) and sfRNA (d) in cell media from cells infected with MOI 0.05 or 1 at 72 hpi. **e,f** Levels of gRNA (e) and sfRNA (f) in carcasses from mock and infected mosquitoes at 10 dpi. **g-h** Levels of gRNA (g) and sfRNA (h) in salivary glands from mock and infected mosquitoes at 10 dpi. a-h Bars indicate means ± s.e.m. Dots indicate repeats. n, 3. *, p < 0.05; ****, p < 0.0001, as determined by T-test.

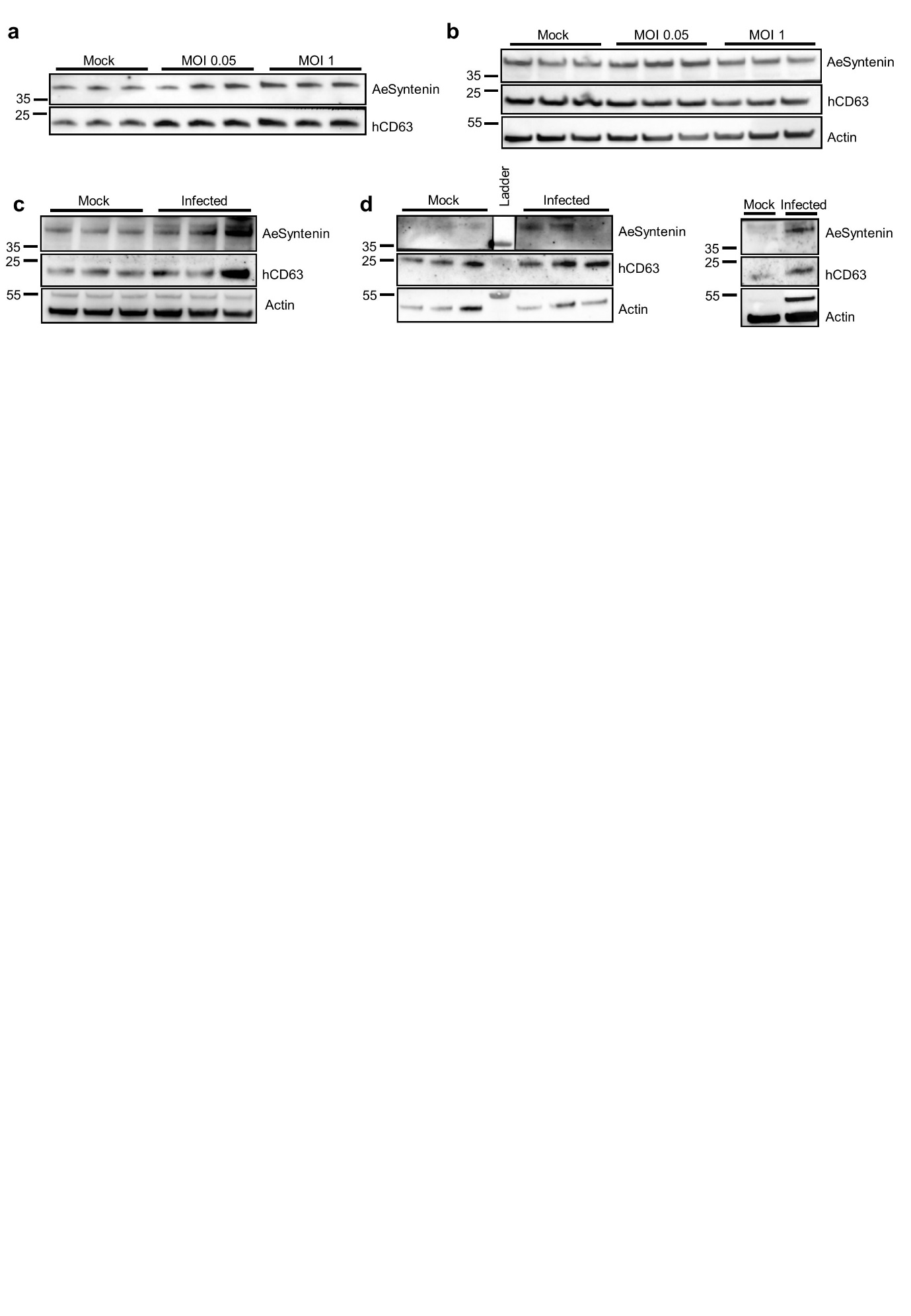

Fig. S2 | Repeats of the quantification of AeSyntenin and hCD63 in infected and mock-infected Aag2 mosquito cells and mosquitoes, related to Fig. 1. **a,b** WB of AeSyntenin and hCD63 in supernatant (a) and cells (b) infected with MOI 0.05 or 1 at 72 hpi. **c,d** WB of AeSyntenin and hCD63 in carcasses (c) and salivary glands (d) infected mosquitoes at 10 dpi.

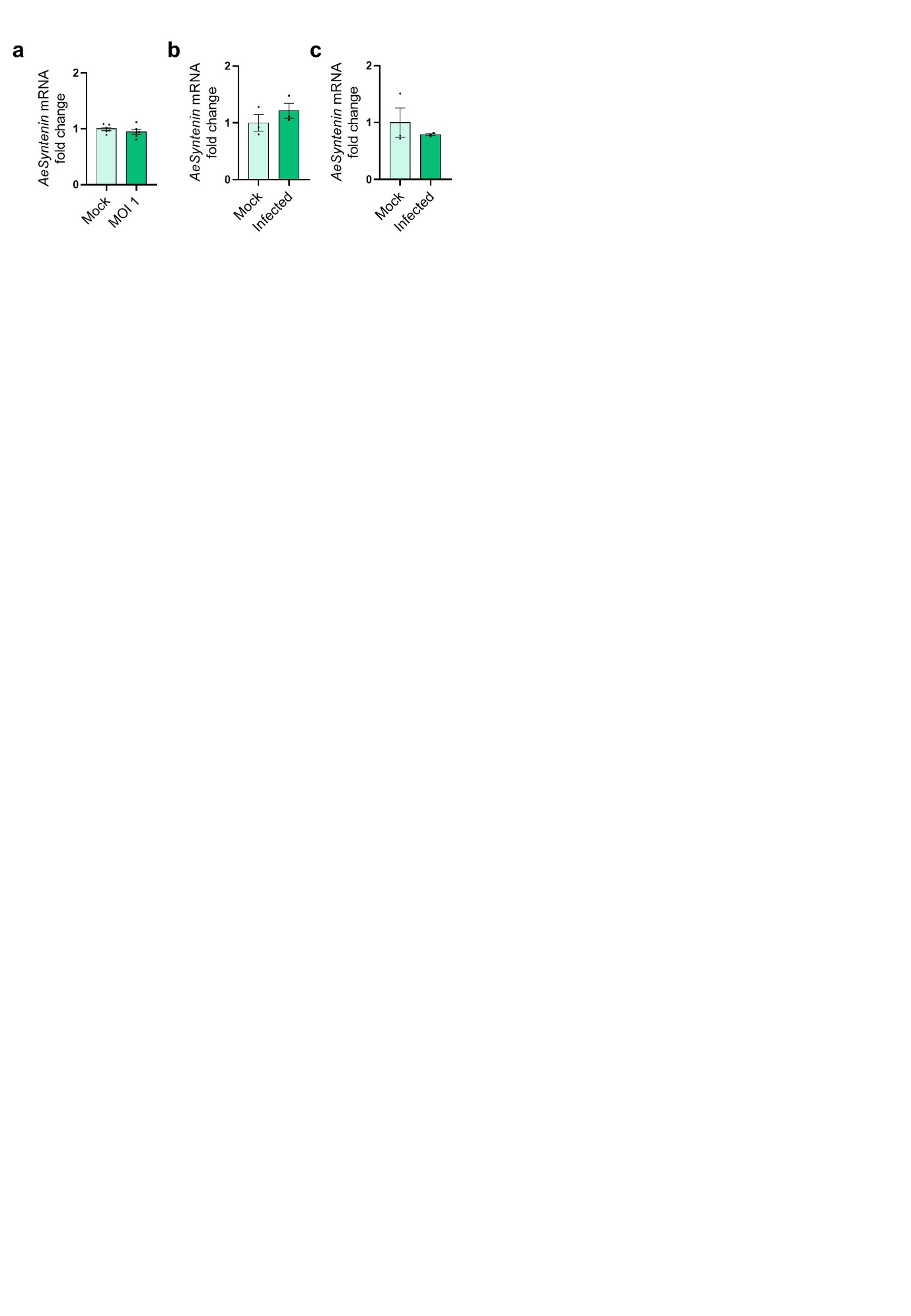

Fig. S3 | Quantification of *AeSyntenin* expression in infected and mock-infected Aag2 mosquito cells and mosquitoes, related to Fig. 1. **a-c** *AeSyntenin* mRNA levels in cells (a), carcasses (b) and salivary glands (c) infected with DENV at 10 dpi. a-c Bars indicate means ± s.e.m. Dots indicate repeats. n for a, 6. n for b and c, 3.

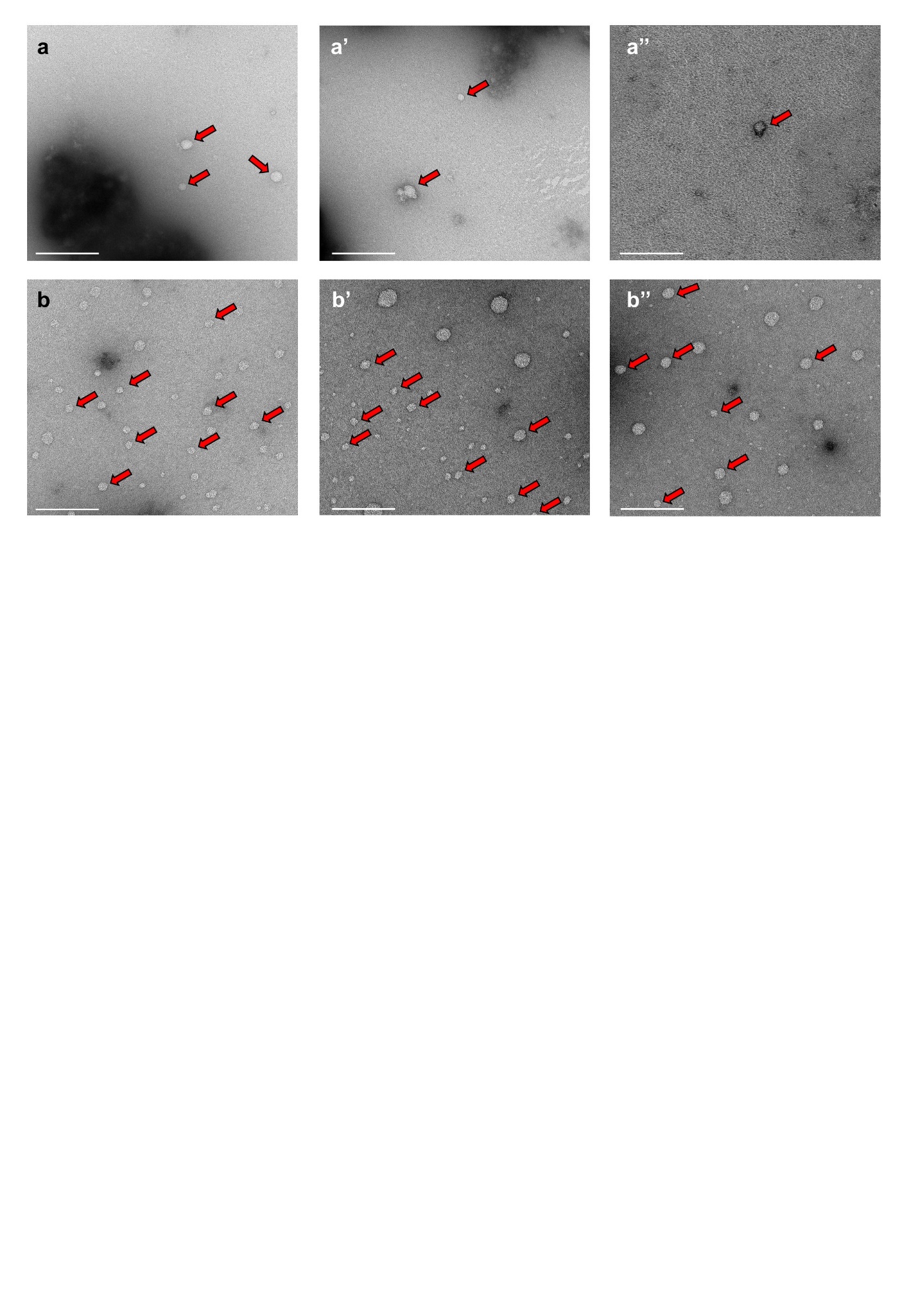

Fig. S4 | Visualization of salivary EVs from infected mosquitoes, related to Fig. 1. **a-b’’** Representative TEM pictures of saliva from mock (a-a'’) and from DENV infected mosquitoes (b-b'’). Scale bar, 200 nm. Red arrows indicate EVs.

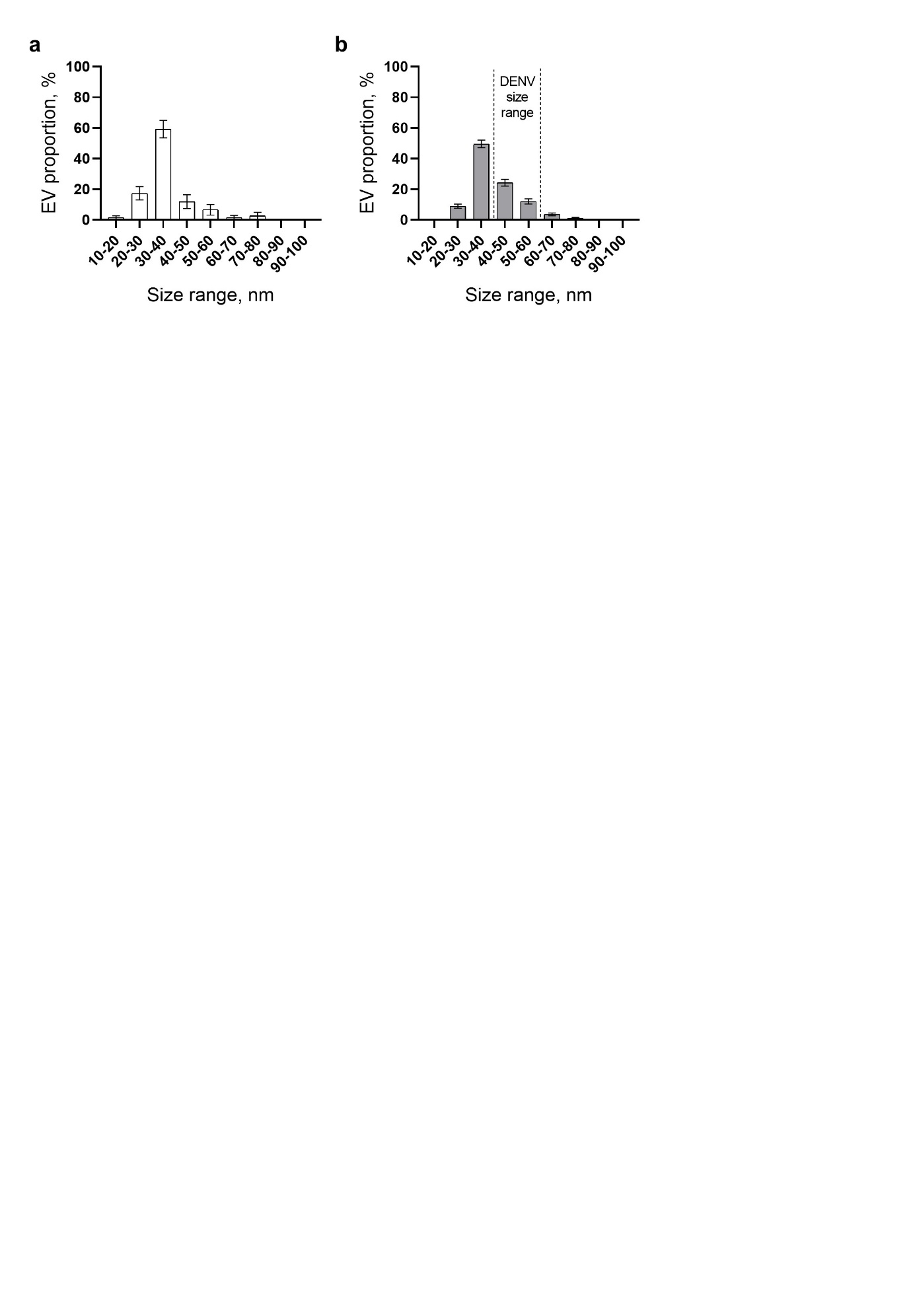

Fig. S5 | Size distribution of EVs in saliva from infected mosquitoes, related to Fig. 1. **a,b** Distribution of EVs per size range in mock (a) and infected mosquitoes (b) according to TEM. Bars show proportion ± SD. n EVs for a, 75; and for b, 450.

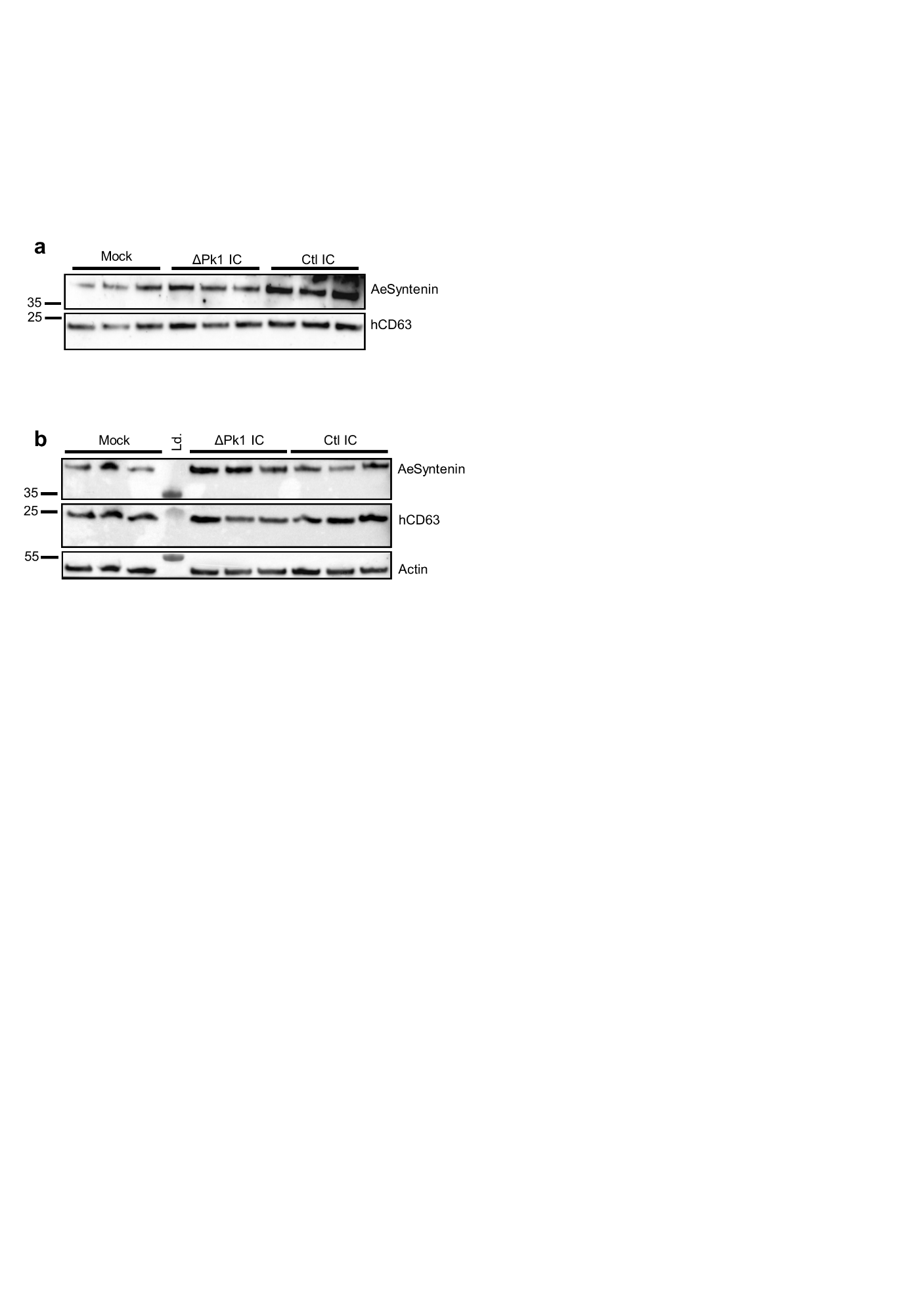

Fig. S6 | Repeats of the quantification of AeSyntenin and hCD63 in ΔPk1-infected Aag2 cells and supernatant, related to Fig. 2. **a,b** WB of AeSyntenin and hCD63 in cell media (a) and cells (b) infected with mock, ΔPk1 IC and control non-mutated DENV IC. Actin was used as a loading control.

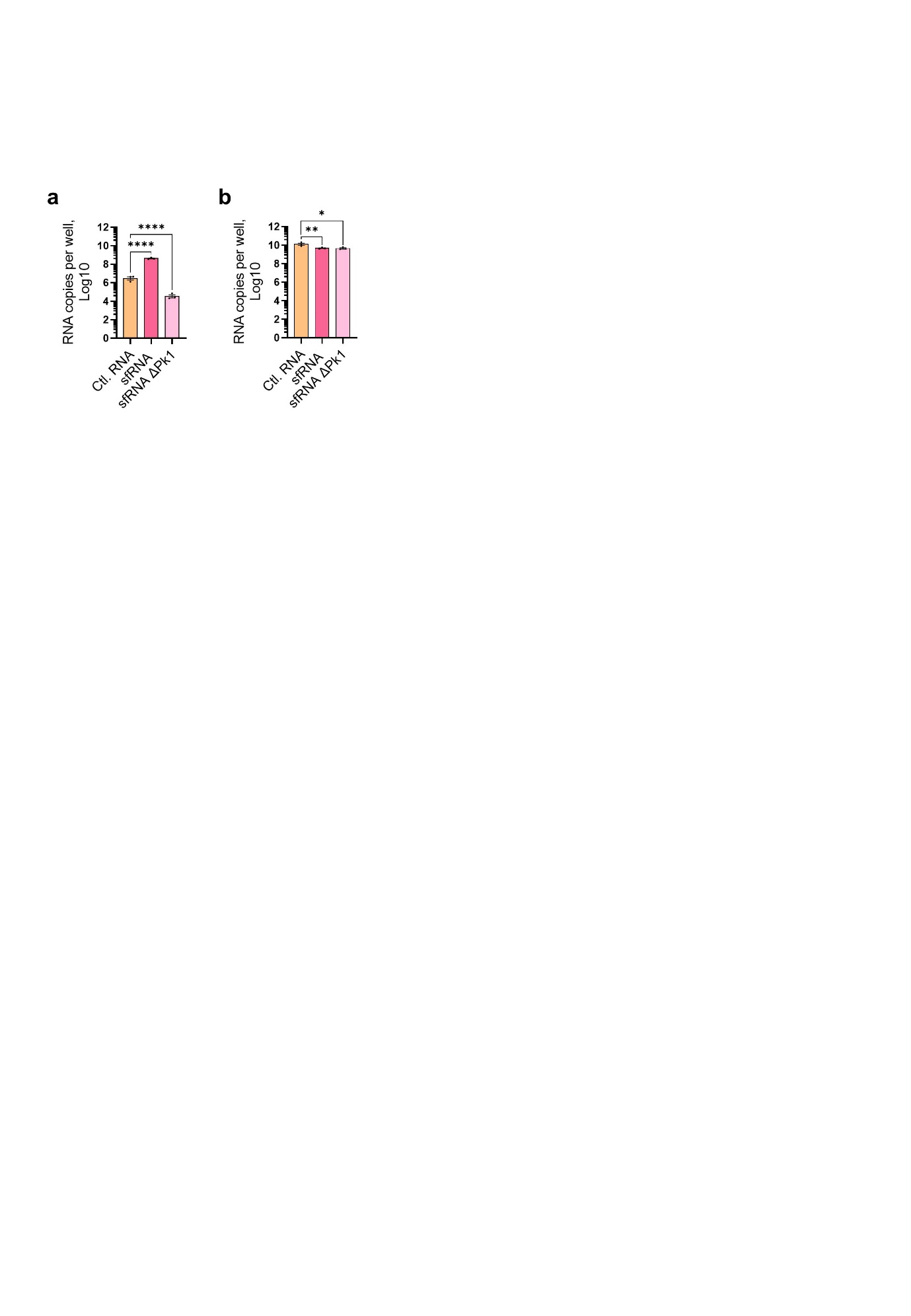

Fig. S7 | Quantification of transfected RNA fragments, related to Fig. 2. **a** Levels of intracellular sfRNA, ΔPk1 sfRNA and Ctl. RNA just after transfection with 10^10^ copies/µl. Bars show geometric means ± 95% C.I. **b** Levels of intracellular sfRNA, ΔPk1 sfRNA and Ctl. RNA just after transfection with 100 more copies of Ctl. RNA and 14,000 more copies of ΔPk1 sfRNA. Bars show means ± s.e.m. Dots indicate biological replicates. n, 3. *, p < 0.05; **, p < 0.01; ****, p < 0.0001, as determined by Fisher’s LSD test.

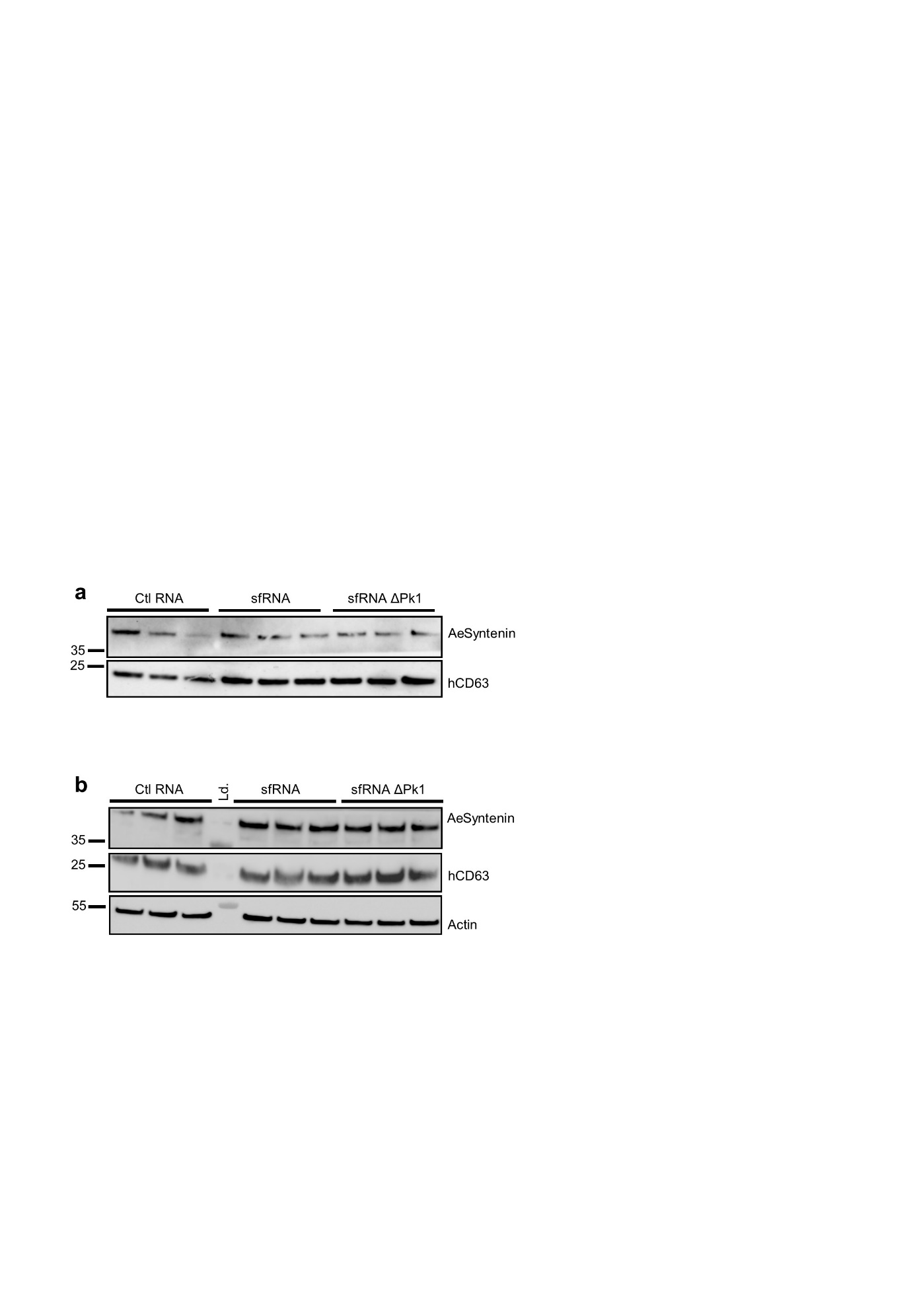

Fig. S8 | Repeats of the quantification of AeSyntenin and hCD63 in sfRNA-transfected Aag2 cells and supernatant, related to Fig. 2. **a,b** WB of AeSyntenin and hCD63 in cell media (a) and in cells (b) after transfection with sfRNA, ΔPk1 sfRNA or Ctl. RNA. Actin was used as a loading control.

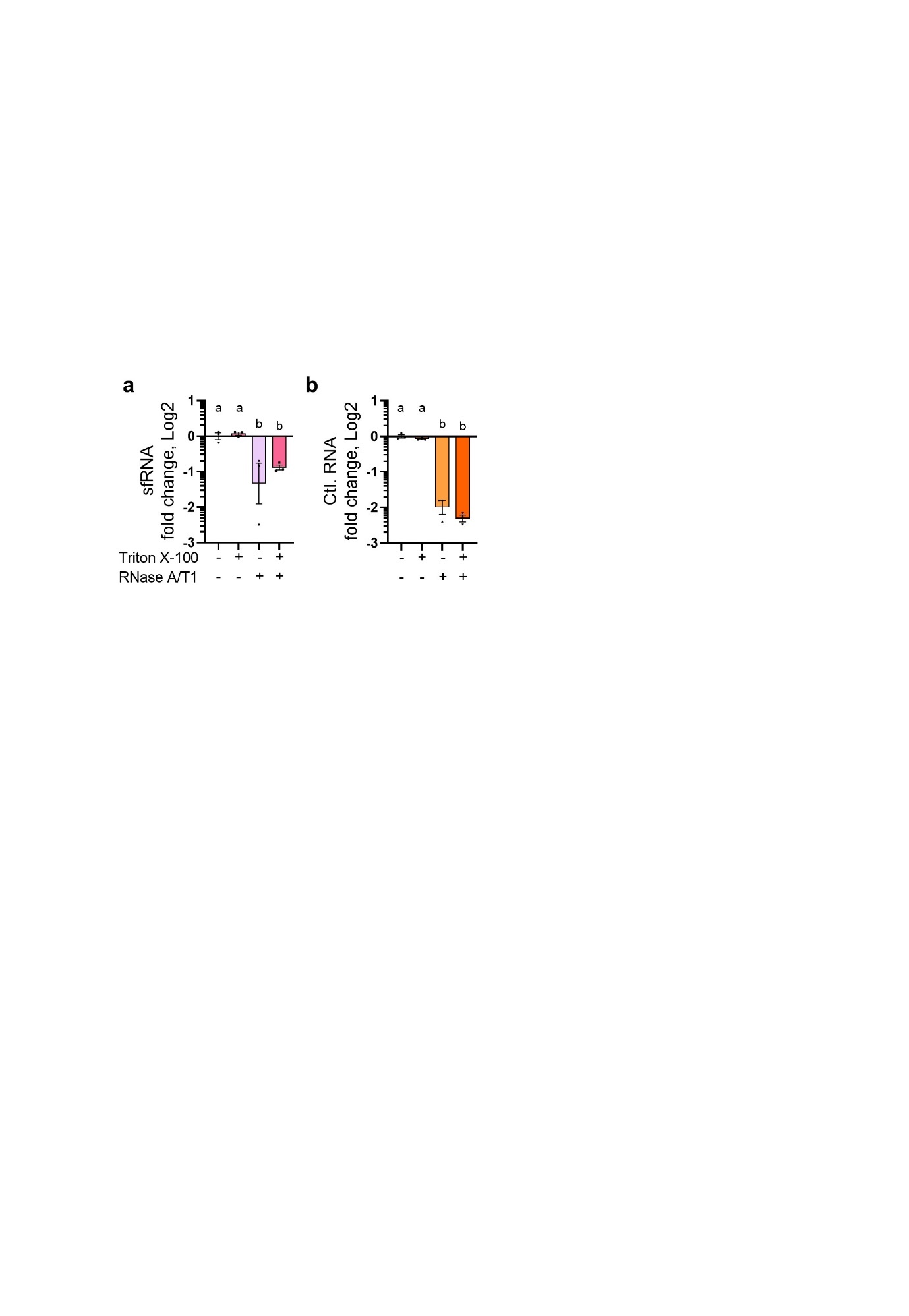

Fig. S9 | Resistance to RNase for *in vitro*-transcribed Ctl. RNA and sfRNA, related to Fig. 2. *In vitro*-transcribed RNA fragments were mixed with cell media and treated with Triton X-100 and RNase A/T1. **a,b** Levels of sfRNA (a) and Ctl. RNA (b) after RNase treatment with or without pre-treatment with Triton X-100. Different letters indicate significant differences as determined by Fisher’s LSD test. Dots indicate biological repeats. n, 3.

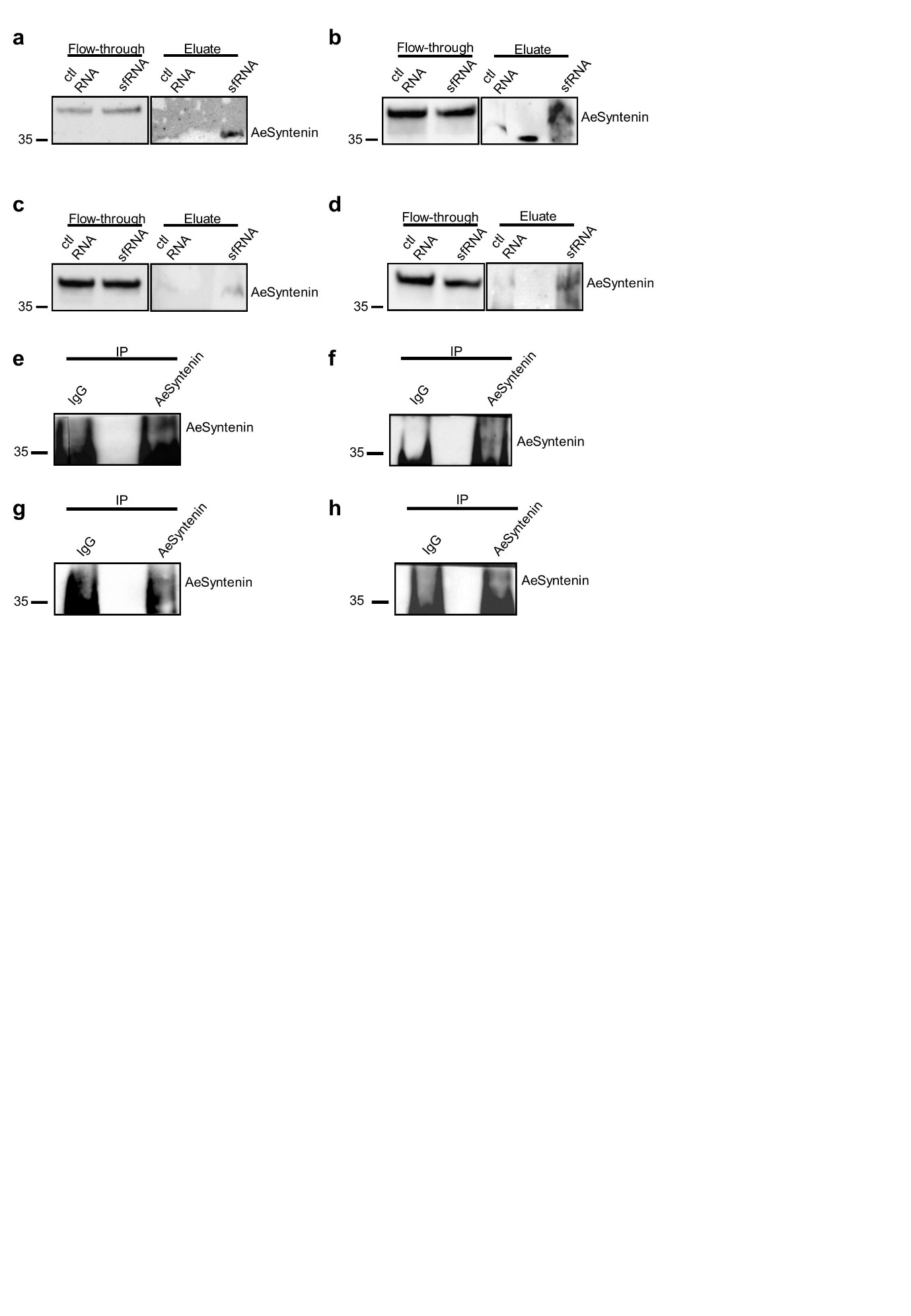

Fig. S10 | Repeats for the detection of AeSyntenin in RNA-affinity chromatography eluates and RIP, related to Fig. 3. **a-d** Repeats of AeSyntenin detection in flow-through and eluates from RNA-affinity chromatography conducted with EV (a,b) and cell (c,d) lysates. **e-h** Repeats of AeSyntenin detection in flow-through and RNA immunoprecipitates (RIP) from EV (e,f) and cell (g,h) infected with DENV.

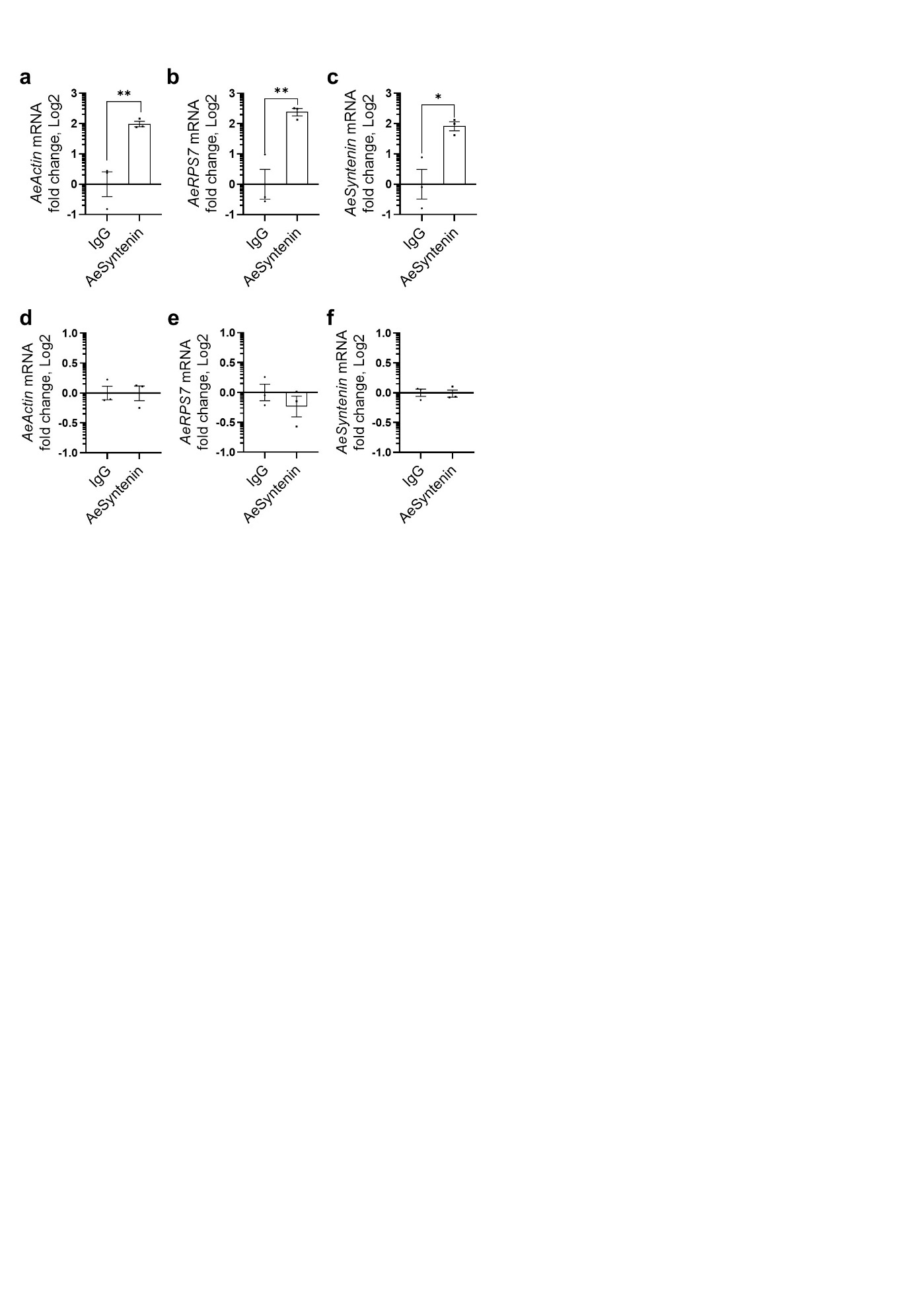

Fig. S11 | Quantification of mRNAs in RIP with anti-AeSyntenin, related to Fig. 3. **a-c** *AeActin* (a), *AeRPS7* (b) and *AeSyntenin* (c) mRNA in RIP with anti-AeSyntenin and IgG control in infected-cell lysates. **d-f** *AeActin* (d), *AeRPS7* (e) and *AeSyntenin* (f) mRNA in RIP with anti-AeSyntenin and IgG control in EV lysates from infected cells. a-f Bars indicate means ± s.e.m. Dots indicate repeats. n, 3. *, p < 0.05; **, p < 0.01, as determined by T-test.

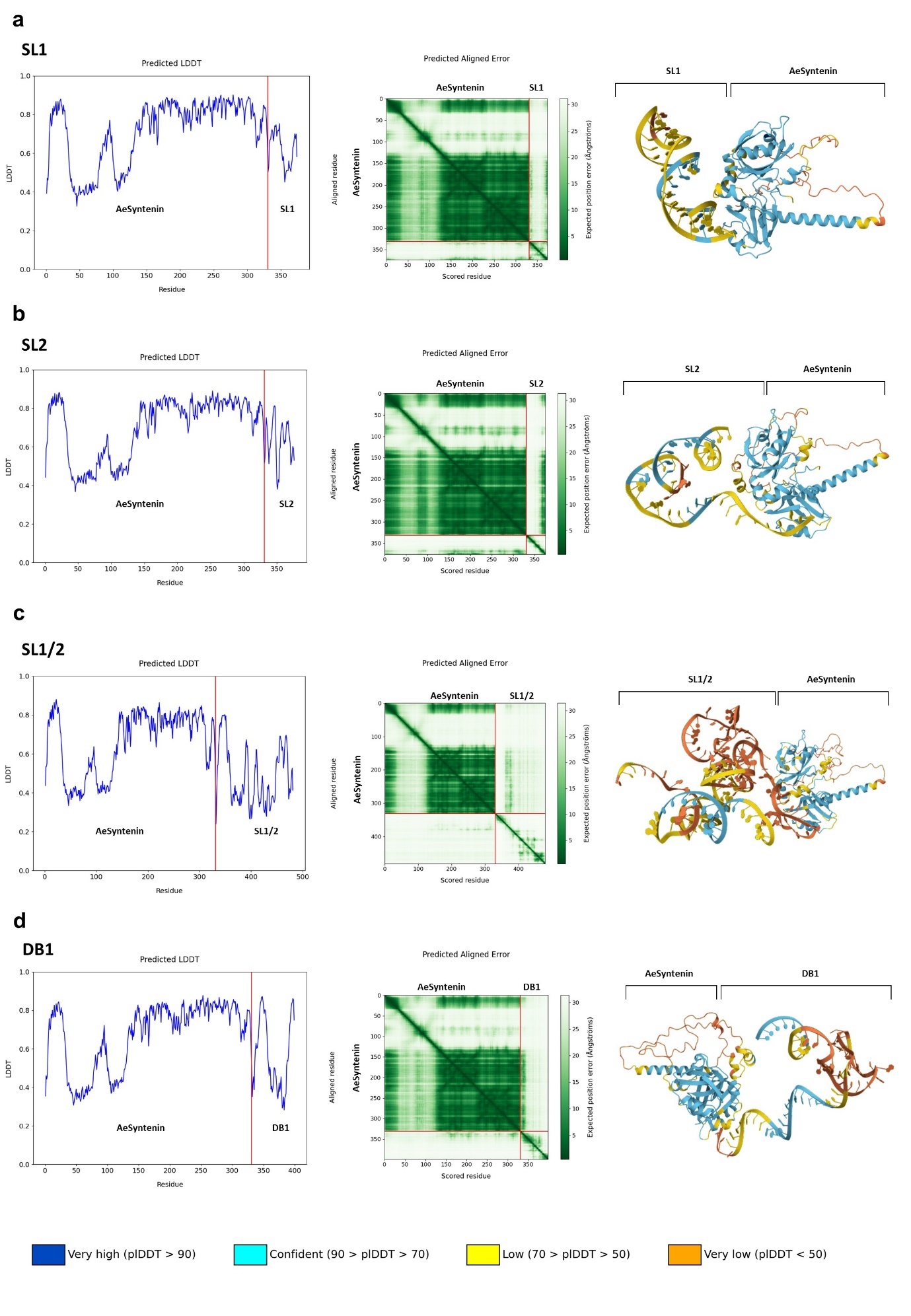

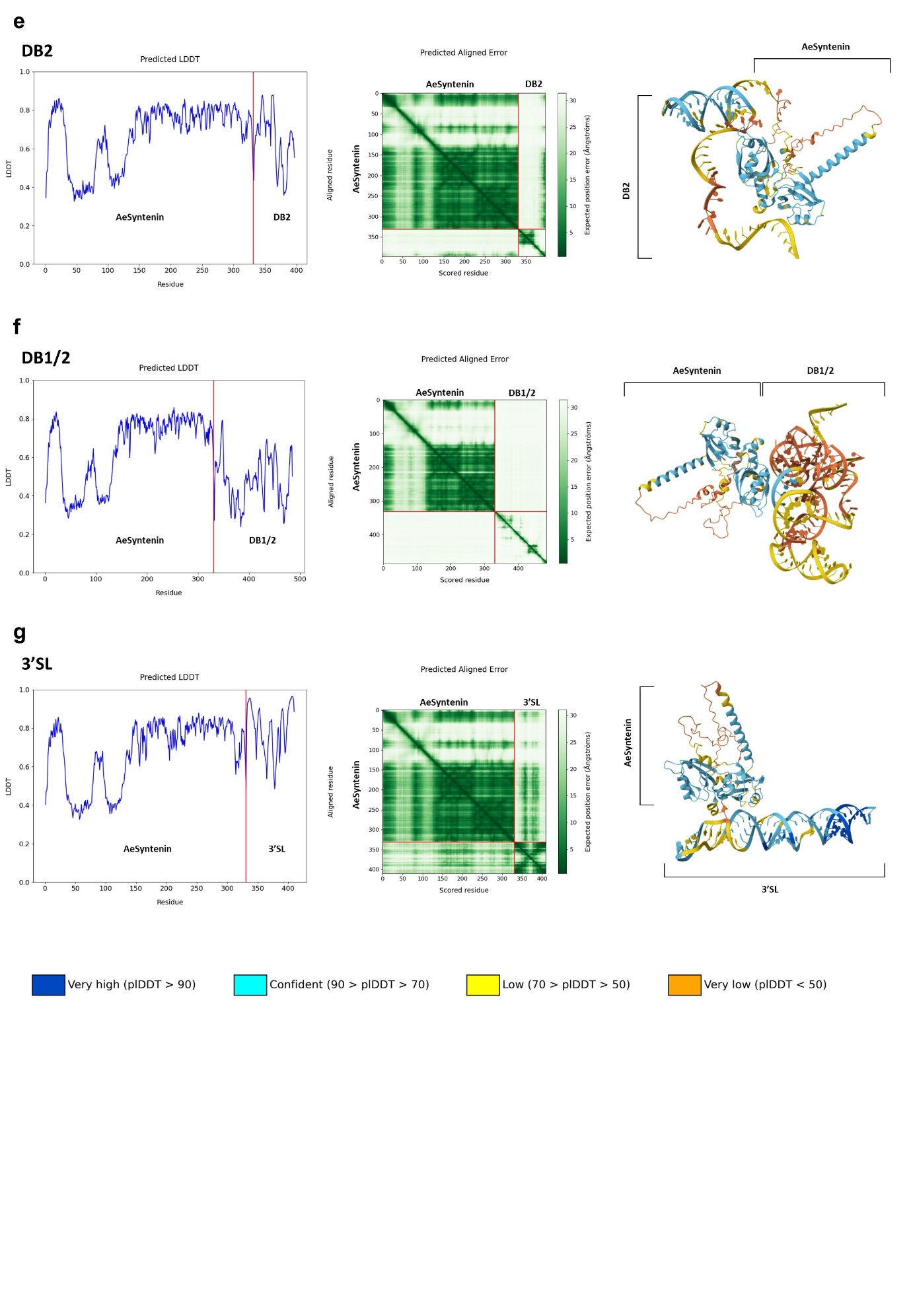

Fig. S12 | Interaction predictions between DENV sfRNA structures and AeSyntenin, related to Fig 3. **a-g** Prediction of interactions between AeSyntenin and SL1 (a), SL2 (b), SL1/2 (c), DB1 (d), DB2 (e), DB1/2 (f), and 3’ SL structures (g). For each interaction, (i) the predicted Local Distance Difference Test (LDDT) is shown to indicate the confidence level in the predicted structure per residue, (ii) the Predicted Aligned Error (PAE) plot is shown to estimate the expected positional distance between two residues in Ångströms (Å) - A lower PAE value (dark green) for a pair of residues provide confidence in their interaction, whereas a higher PAE value (light green) suggests that the residues do not interact; and (iii) the 3D predicted protein-RNA structure is shown with a color code reflecting the pLDDT score - low confidence is indicated in red and high confidence in blue.

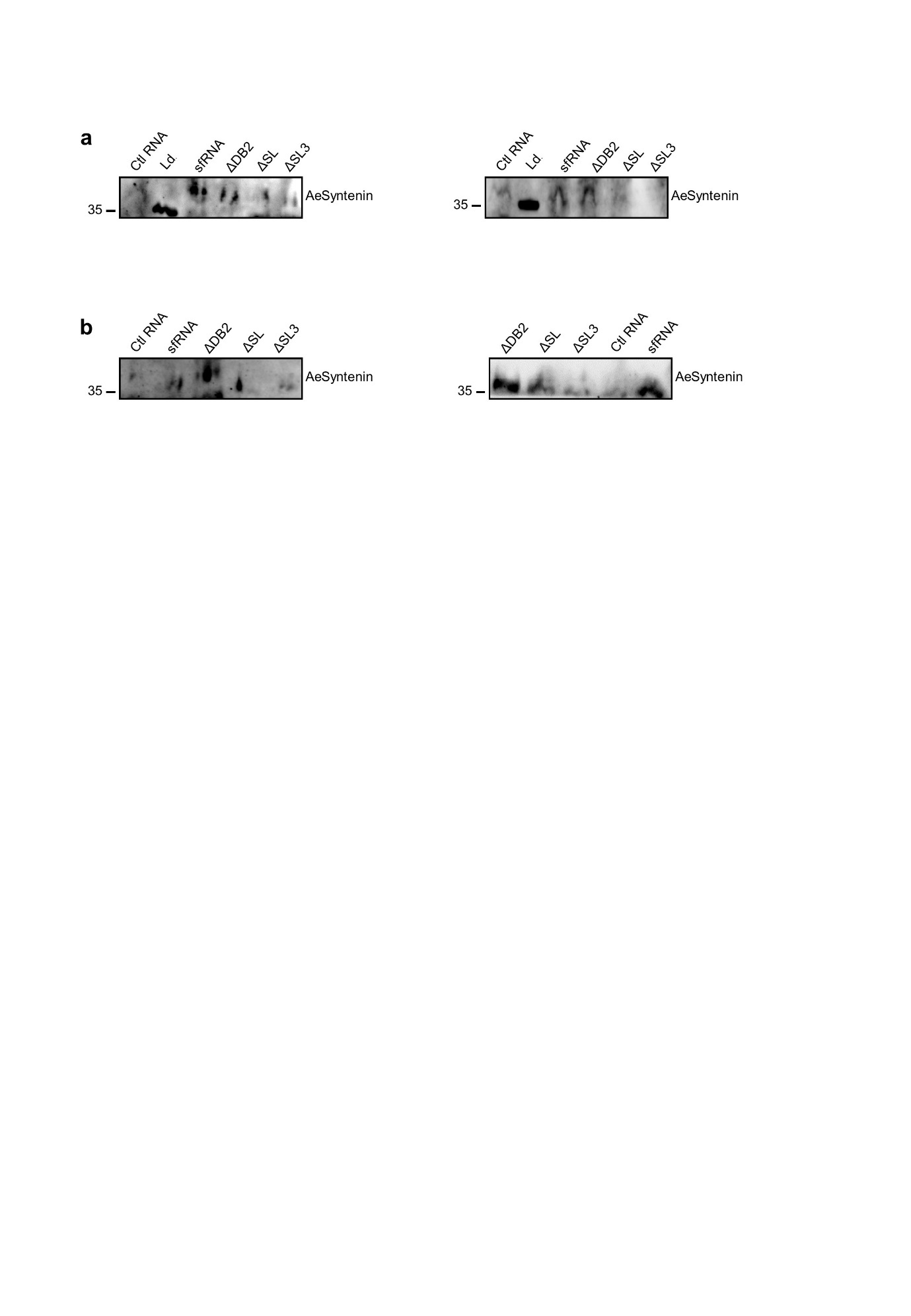

Fig. S13 | Repeats for the detection of AeSyntenin in RNA-affinity chromatography eluates with sfRNA fragments, related to Fig. 3. **a-d** Repeats of AeSyntenin detection in flow-through and eluates from RNA-affinity chromatography conducted with EV (a,b) and cell (c,d) lysates. **e-h** Repeats of AeSyntenin detection in flow-through and RNA immunoprecipitates (RIP) from EV (e,f) and cell (g,h) infected with DENV.

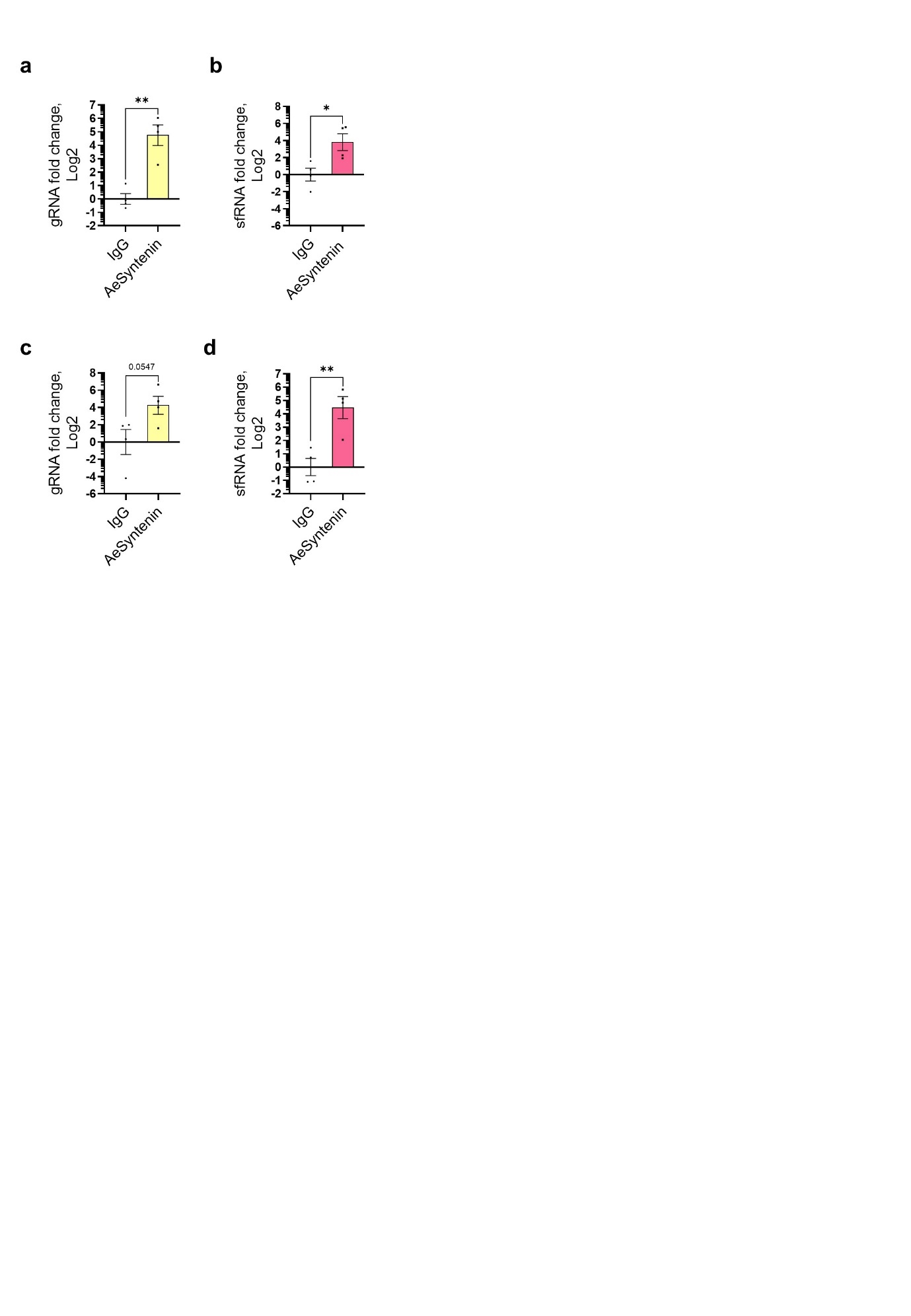

Fig. S14 | AeSyntenin IP and WNV gRNA, sfRNA quantification related to Fig. 3. **a,b** gRNA level (a) and sfRNA level (b) in AeSyntenin RIP eluates from WNV-infected cells. **c,d** gRNA level (a) and sfRNA level (b) in AeSyntenin RIP eluates from EVs from WNV-infected cells. Bars indicate means ± s.e.m. Dots indicate repeats. n, 4. *, p < 0.05; **, p < 0.01, as determined by T-test.

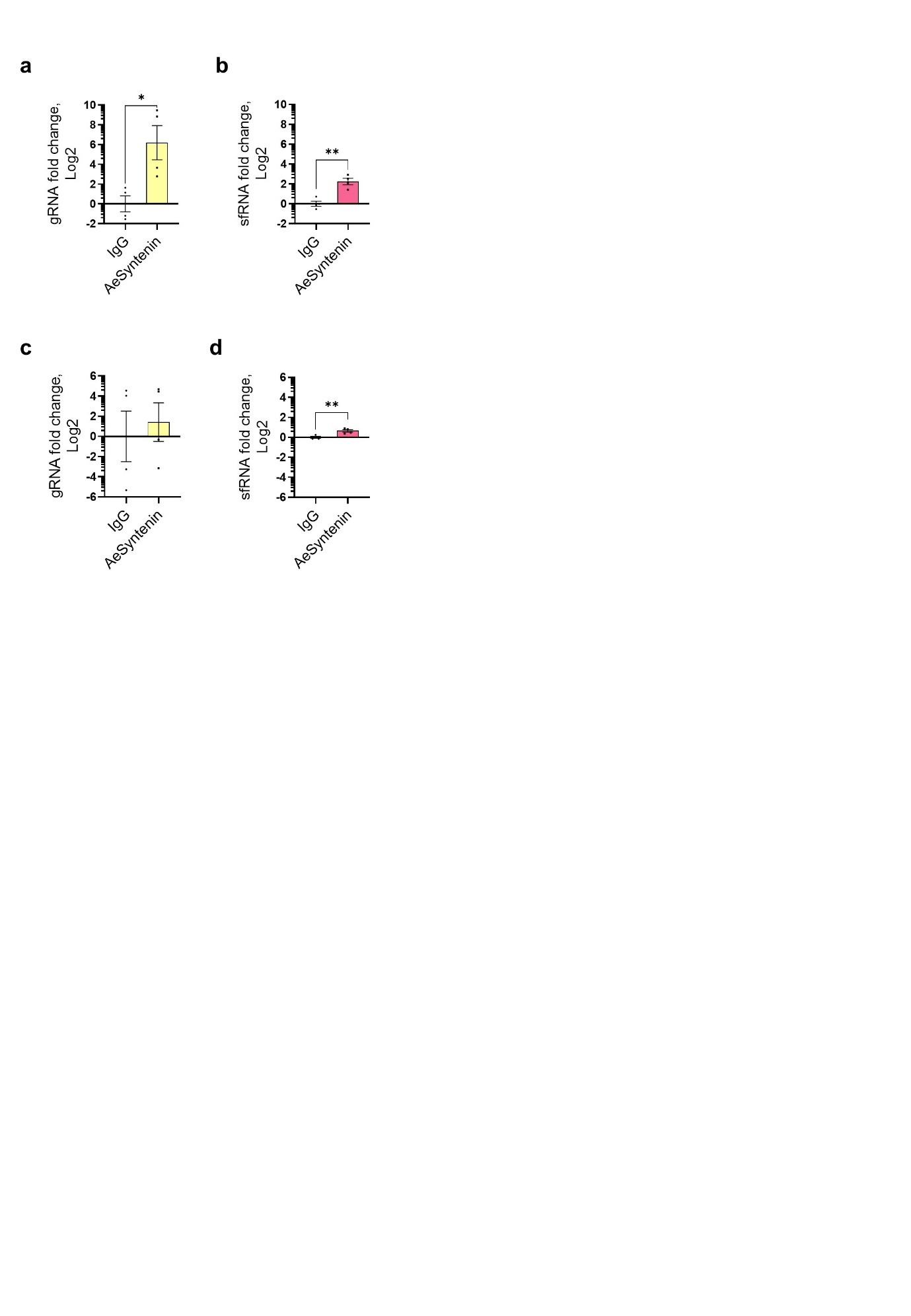

Fig. S15 | AeSyntenin IP and ZIKV gRNA, sfRNA quantification related to Fig. 3. **a,b** gRNA level (a) and sfRNA level (b) in AeSyntenin RIP eluates from ZIKV-infected cells. **c,d** gRNA level (a) and sfRNA level (b) in AeSyntenin RIP eluates from EVs from ZIKV-infected cells. Bars indicate means ± s.e.m. Dots indicate repeats. n, 4. *, p < 0.05; **, p < 0.01, as determined by T-test.

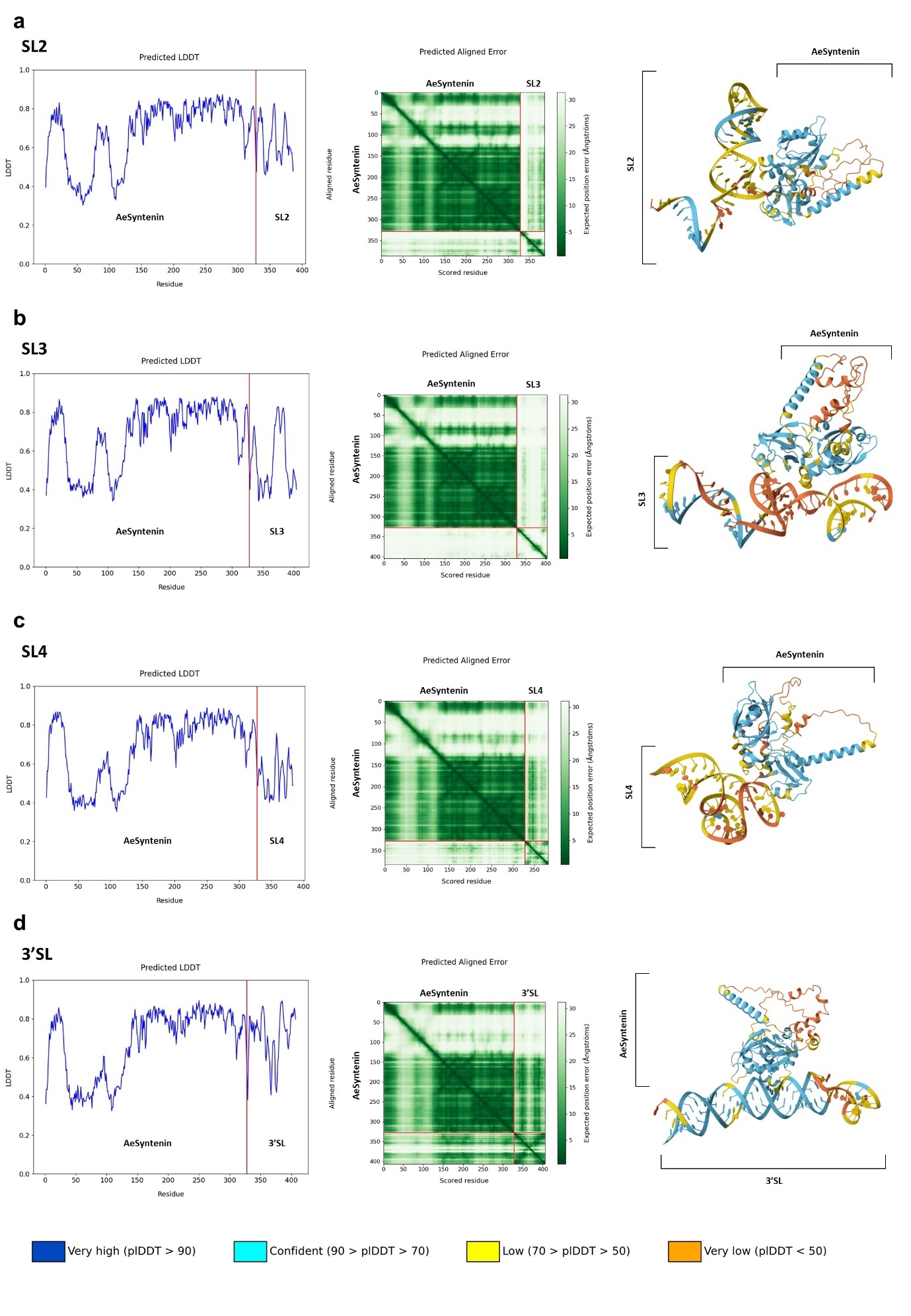

Fig. S16 | Interaction predictions between WNV sfRNA structures and AeSyntenin, related to Fig 3. **a-d** Prediction of interactions between AeSyntenin and SL2 (a), SL3 (b), SL4 (c) and 3’ SL structures (d). For each interaction, (i) the predicted Local Distance Difference Test (LDDT) is shown to indicate the confidence level in the predicted structure per residue, (ii) the Predicted Aligned Error (PAE) plot is shown to estimate the expected positional distance between two residues in Ångströms (Å) - A lower PAE value (dark green) for a pair of residues provide confidence in their interaction, whereas a higher PAE value (light green) suggests that the residues do not interact; and (iii) the 3D predicted protein-RNA structure is shown with a color code reflecting the pLDDT score - low confidence is indicated in red and high confidence in blue.

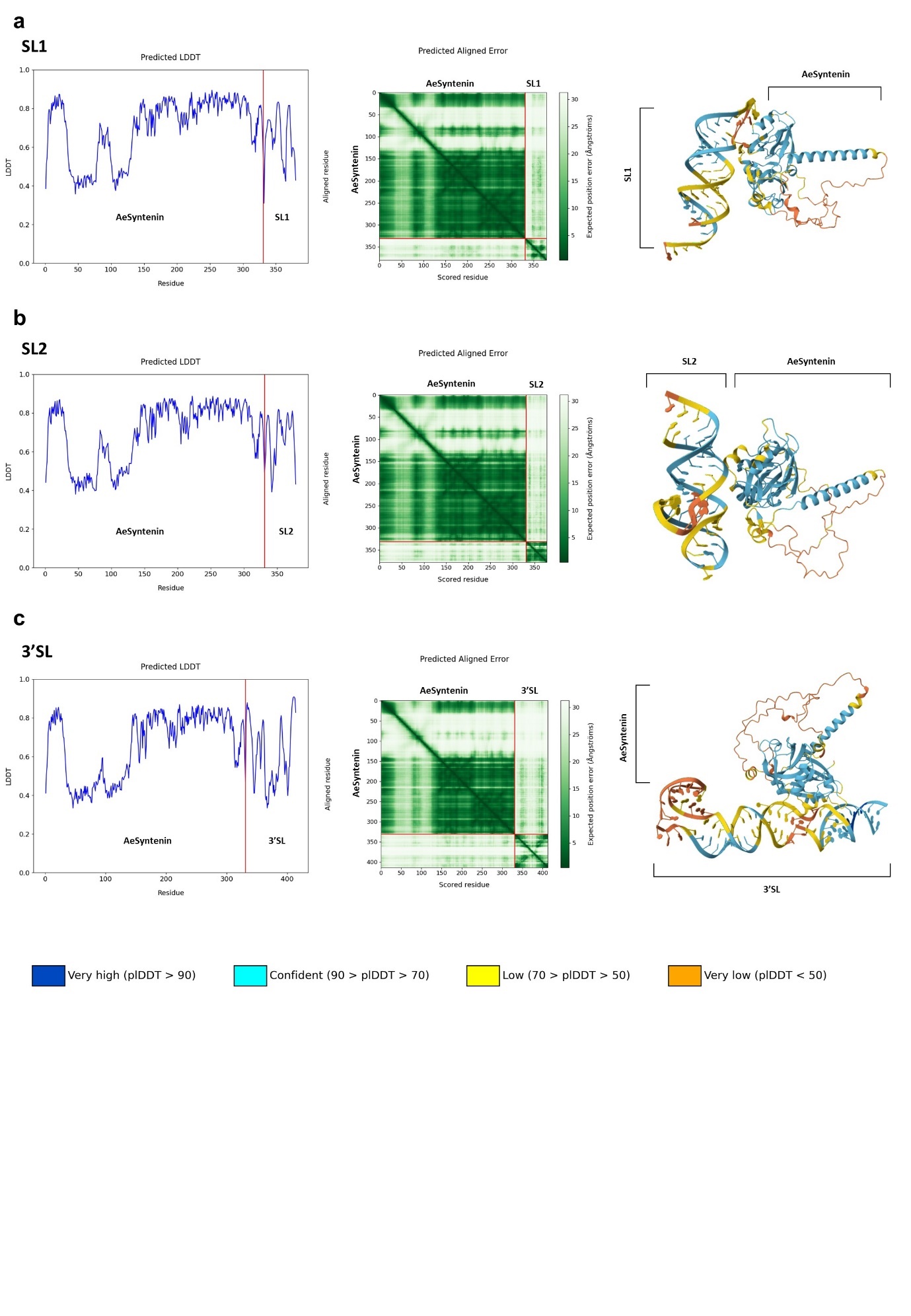

Fig. S17 | Interaction predictions between ZIKV sfRNA structures and AeSyntenin, related to Fig 3. **a-c** Prediction of interactions between AeSyntenin and SL1 (a), SL2 (b) and 3’ SL structures (d). For each interaction, (i) the predicted Local Distance Difference Test (LDDT) is shown to indicate the confidence level in the predicted structure per residue, (ii) the Predicted Aligned Error (PAE) plot is shown to estimate the expected positional distance between two residues in Ångströms (Å) - A lower PAE value (dark green) for a pair of residues provide confidence in their interaction, whereas a higher PAE value (light green) suggests that the residues do not interact; and (iii) the 3D predicted protein-RNA structure is shown with a color code reflecting the pLDDT score - low confidence is indicated in red and high confidence in blue.

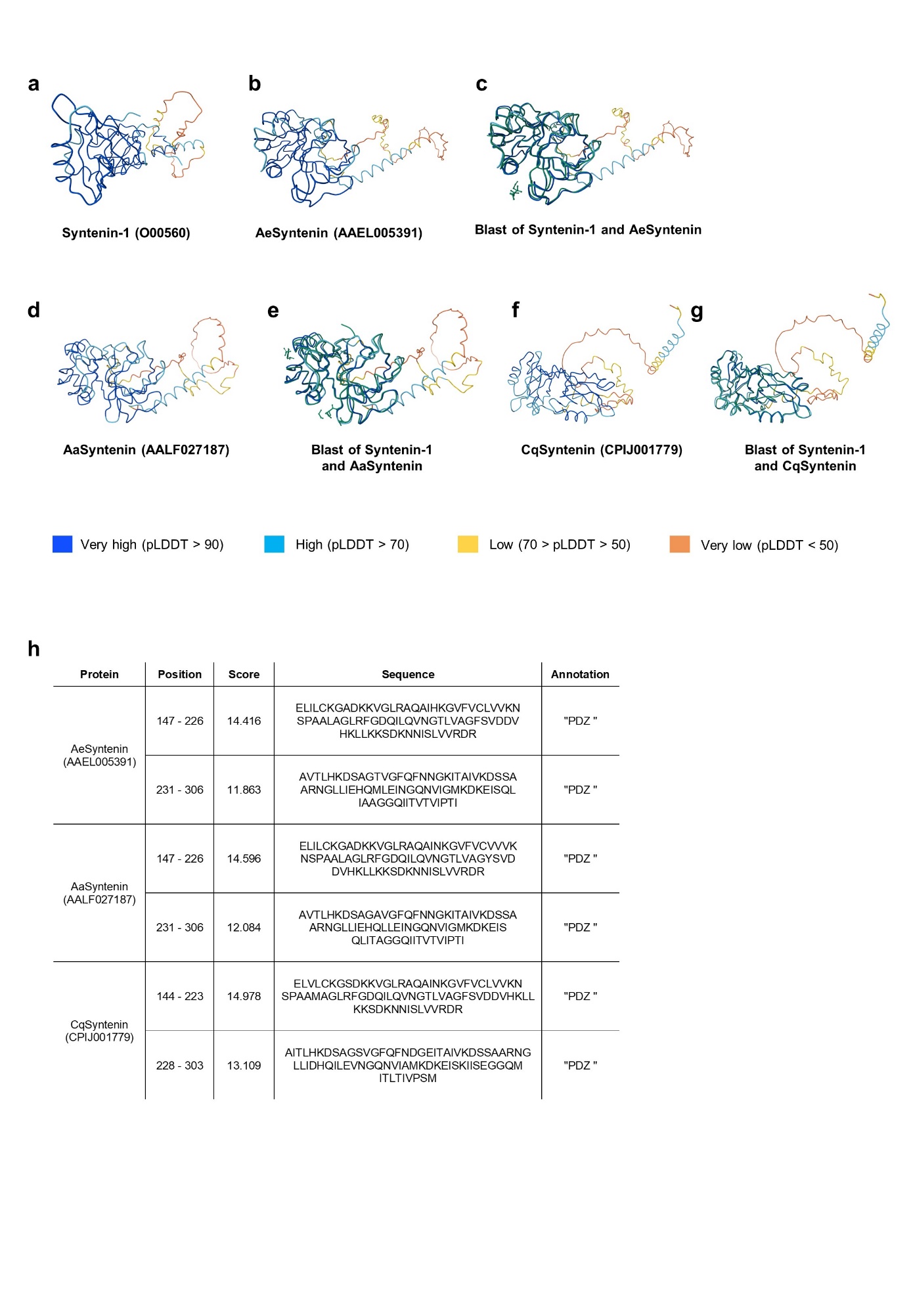

Fig. S18 | Structural comparison between Human syntenin-1 and mosquito *Ae. aegypti* syntenin (AeSyntenin), *Ae. albobictus* syntenin (AaSyntenin) and *Cx. quiquefasciatus* syntenin (CqSyntenin) homologs, related to Fig. 4. **a-c** Predicted 3D structures of Human Syntenin-1 (a), AeSyntenin (b), and structure alignment (BLAST) between the two. **d,e** Predicted 3D structures of AaSyntenin (d), and structure alignment (BLAST) between AaSyntenin and Syntenin-1 (e). **f,g** Predicted 3D structures of CqSyntenin (f), and structure alignment (BLAST) between CqSyntenin and Syntenin-1 (g). **h** Predicted PDZ domains in AeSyntenin, AaSyntenin and CqSyntenin as identified by ScanProsite. A score higher than 11 is indicative of strong confidence in the identification. pLDDT, predicted Local Distance Difference Test.

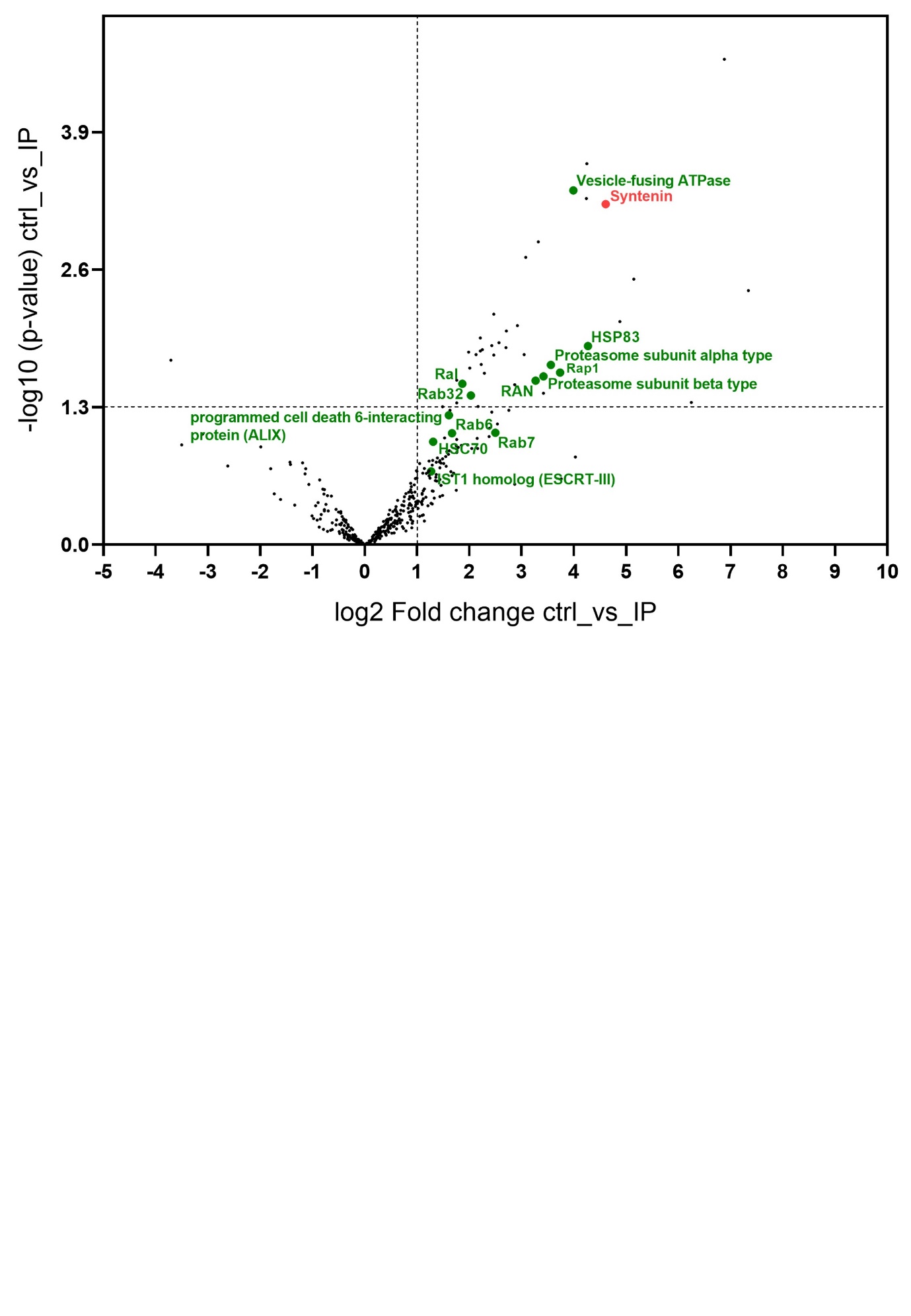

Fig. S19 | Identification of AeSyntenin partners in mosquitoes through co-IP, related to Fig. 4. Volcano plot showing the proteins interacting with AeSyntenin in EVs from DENV-infected Aag2 mosquito cells. Protein names in green indicate proteins usually associated with EVs. Significantly interacting proteins were selected based on Log2 FC > 1 and –log10 (p-value) > 1.3. N, 3.

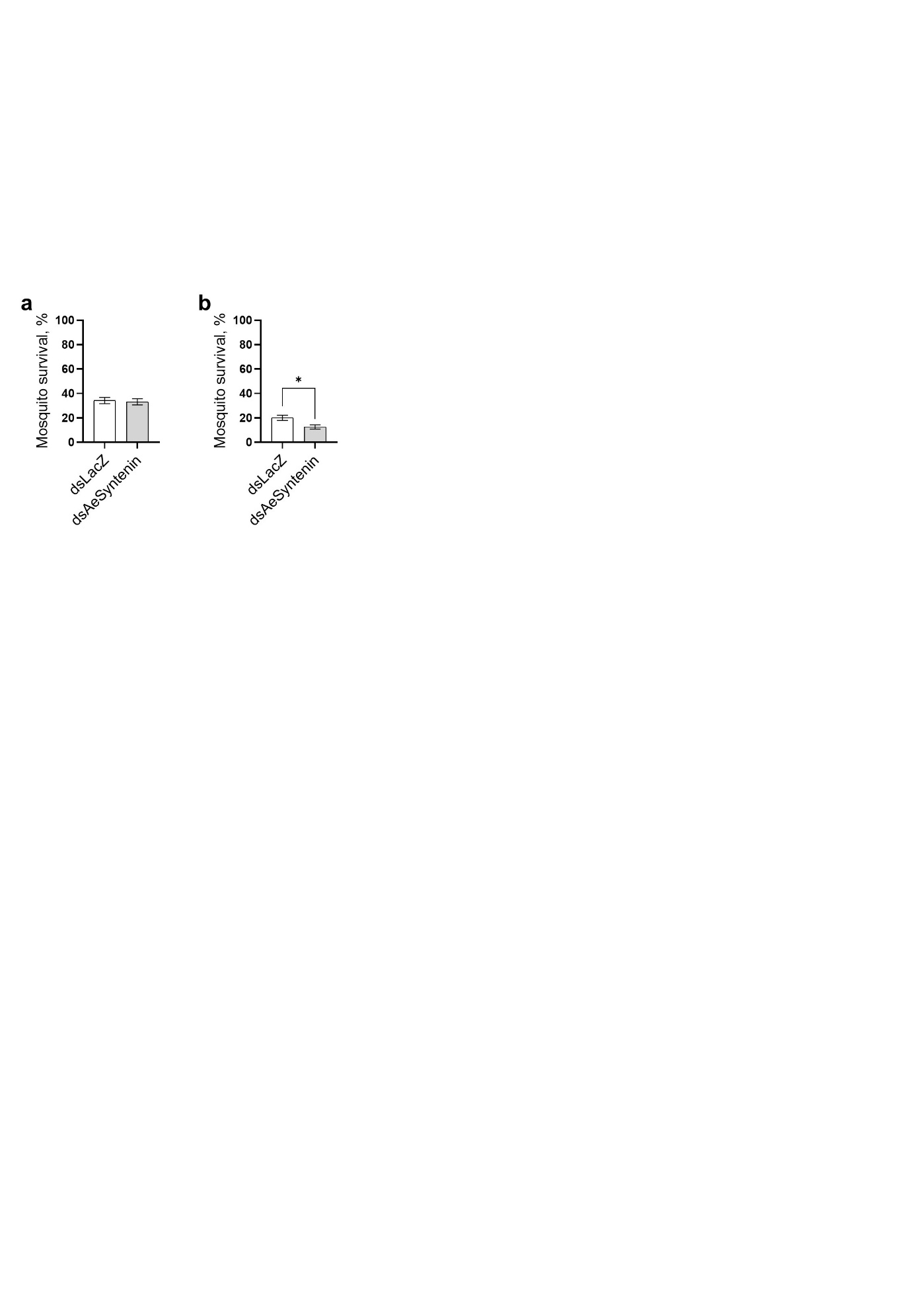

Fig. S20 | Effect of AeSyntenin depletion and infection on mosquito survival, related to Fig. 4. **a,b** Mosquito survival at 4 days post dsRNA injection (on the day of DENV inoculation) (a) and at 10 days post inoculation (on the day of tissue and saliva collection) (b). Bars show percentage ± s.d. n for a, 150. N for b, 100. *, p-value < 0.05 as determined by χ2 test.

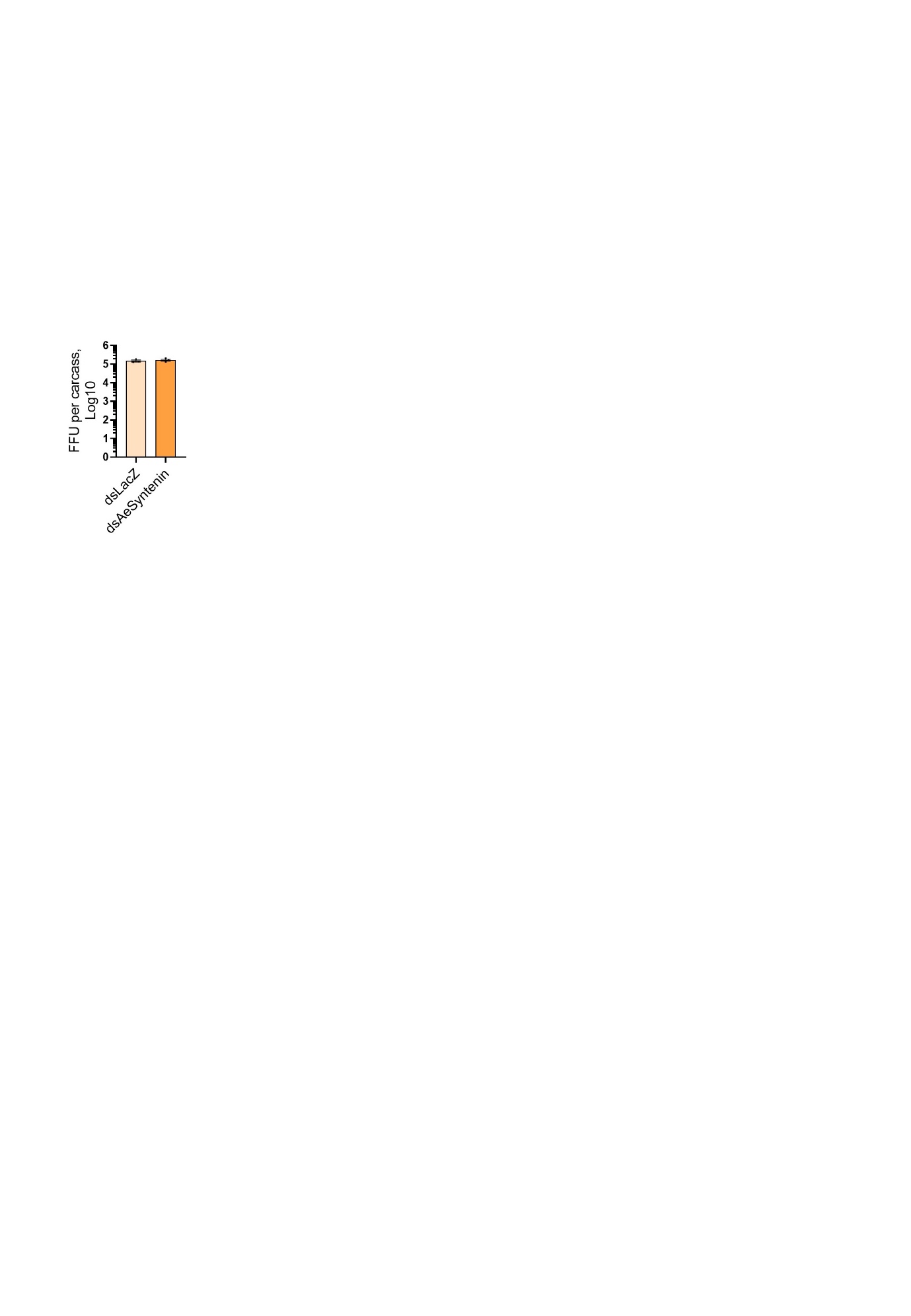

Fig. S21 | Effects of AeSyntenin depletion on infectious particles in mosquitoes, related to Fig.4. Levels of FFU (focus forming unit) in carcasses from mosquitoes injected with dsRNA against AeSyntenin or LacZ (Control). Bars show geometric means ± 95% C.I. n, 3.

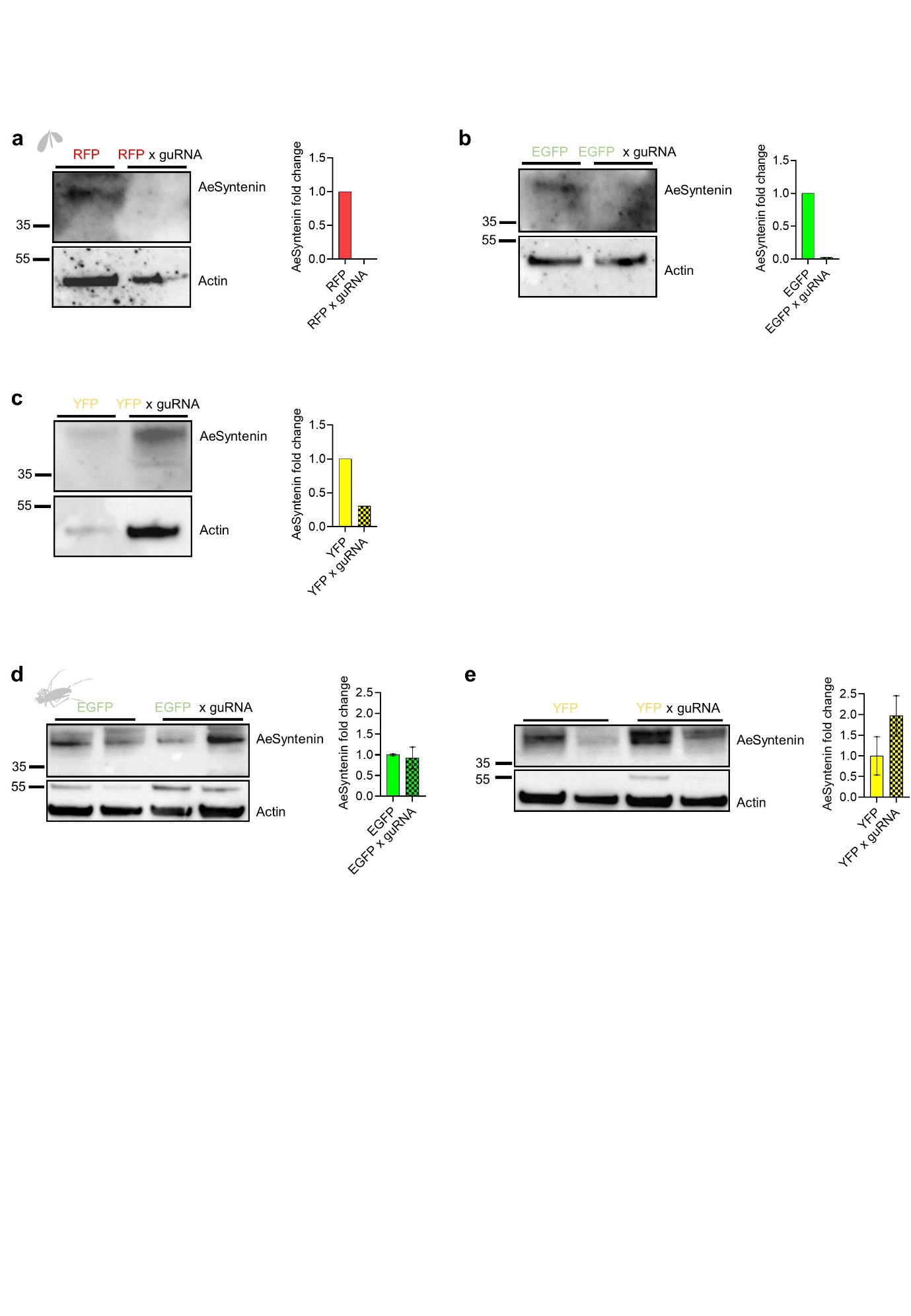

Fig. S22 | AeSyntenin in transgenic mosquitoes, related to Fig. 5. **a-e** AeSyntenin expression in SG (a-c) and carcass (d,e) from control RFP and RFP x guRNA progeny, EGFP and EGFP x guRNA progeny, and YFP and YFP x guRNA progeny. Bar graphs show AeSyntenin fold change normalized to Actin expression.

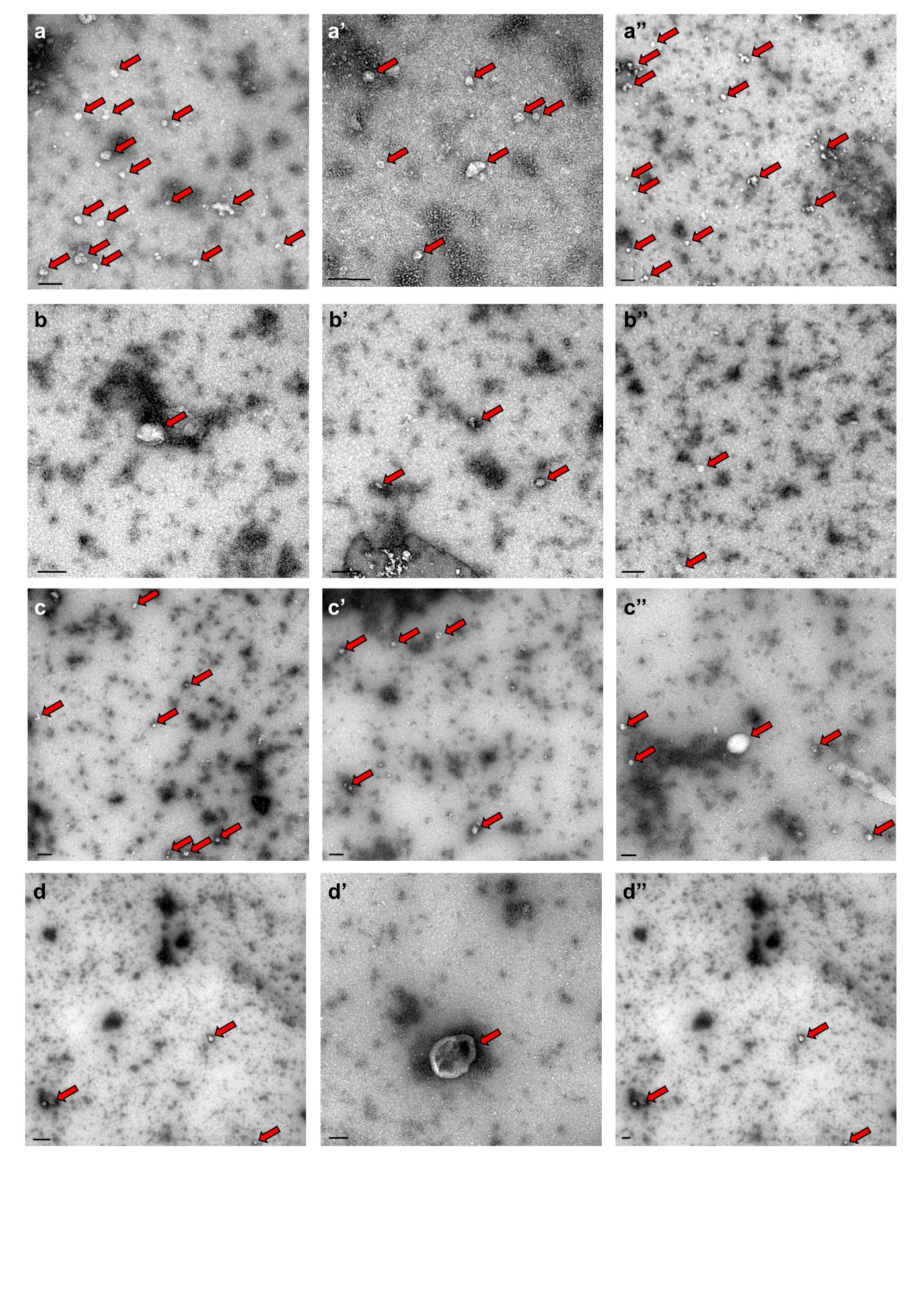

Fig. S23 | Visualization of salivary EVs from SG-deleted AeSyntenin mosquitoes, related to Fig. 5. **a-d'’** Representative TEM pictures of saliva from RFP (a-a'’) and RFP x guRNA progeny (b-b'’), and from EGFP (c-c'’) and EGFP x guRNA progeny (c-c'’). Scale bar, 100 nm. Red arrows indicate EVs.

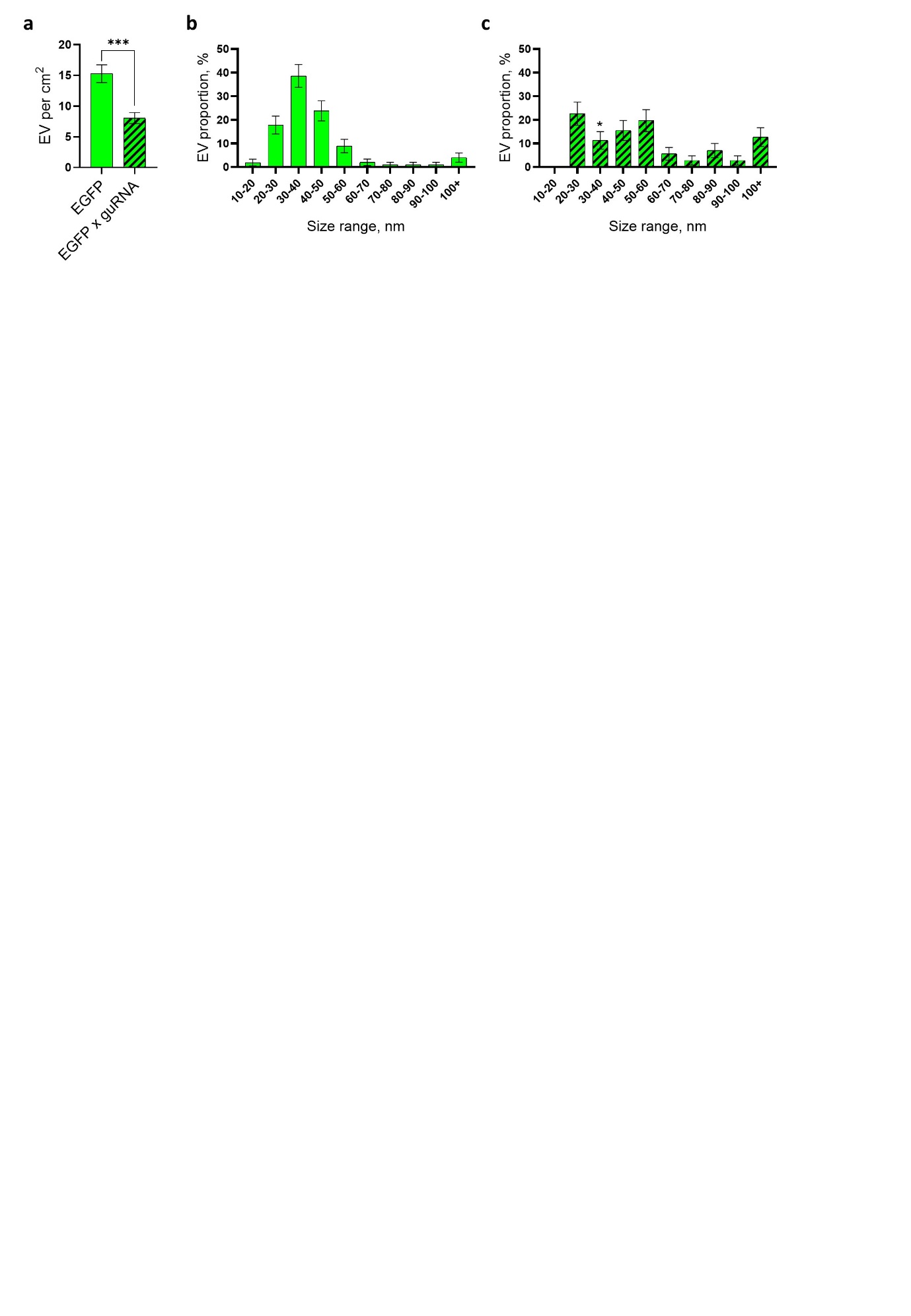

Fig. S24 | Quantification and distribution of EVs in saliva from EGFP x guRNA progeny, related to Fig. 5. **a** EV concentration per cm² of TEM pictures. Bars show mean ± s.e.m. from 43 and 46 pictures for EGFP and EGFP x guRNA, respectively. ***, p < 0.001 as determined by T-test. **b,c** Distribution of EVs per size range in EGFP (b) and EGFP x guRNA progeny (c) according to TEM. Bars show proportion ± SD. n EVs for b, 101; and for c, 71. *, p < 0.05 as determined by Z test same size range in EGFP.

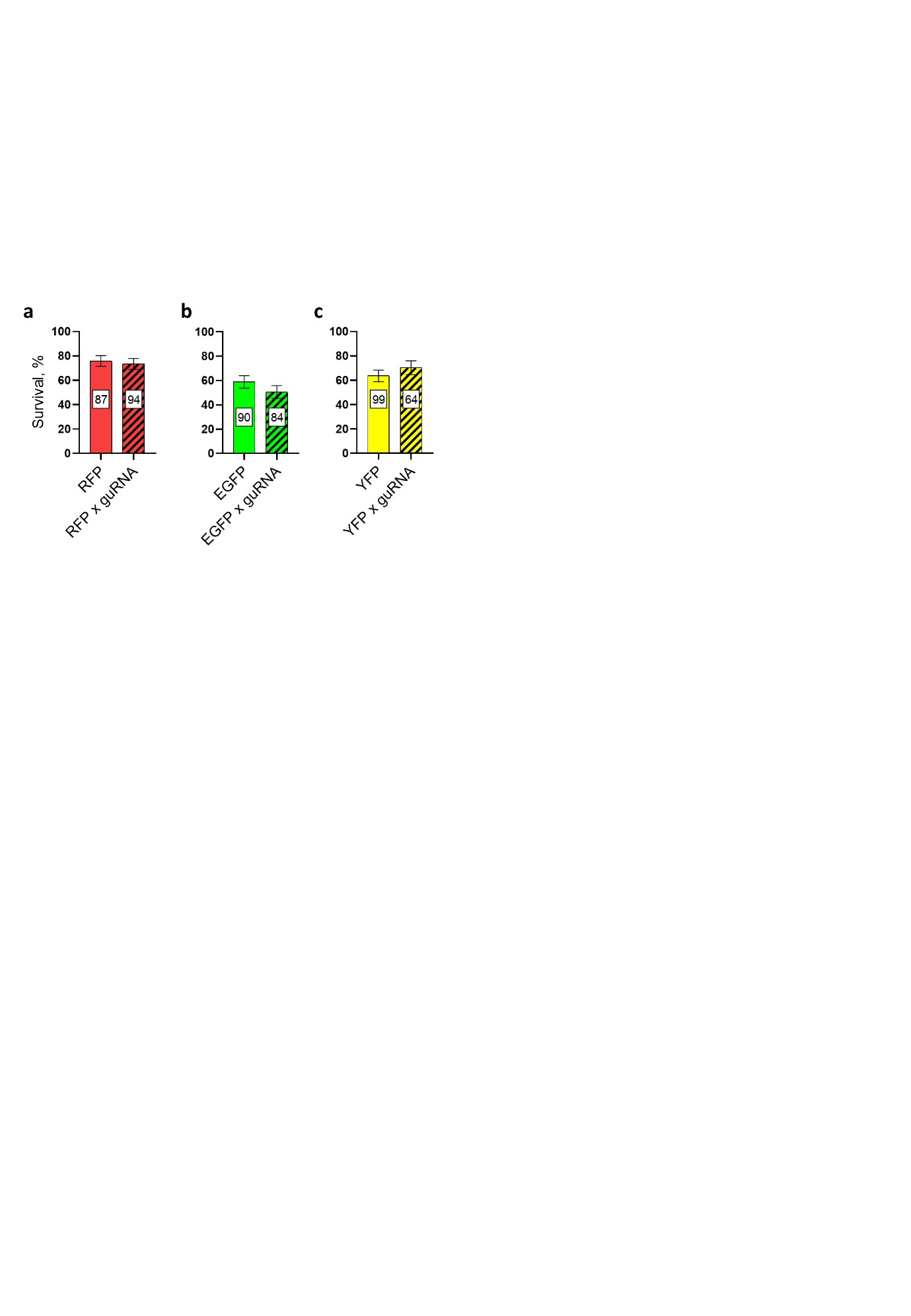

Fig. S25 | Survival of infected SG-AeSyntenin deleted mosquitoes, related to Fig. 5. **a-c** Survival rate at 10 days post DENV infection for RFP and RFP x guRNA (a), EGFP and EGFP x guRNA (b), and YFP and YFP x guRNA (c) mosquitoes. Bars show mean ± s.e.m. from 3 experiments with n indicated within bars.

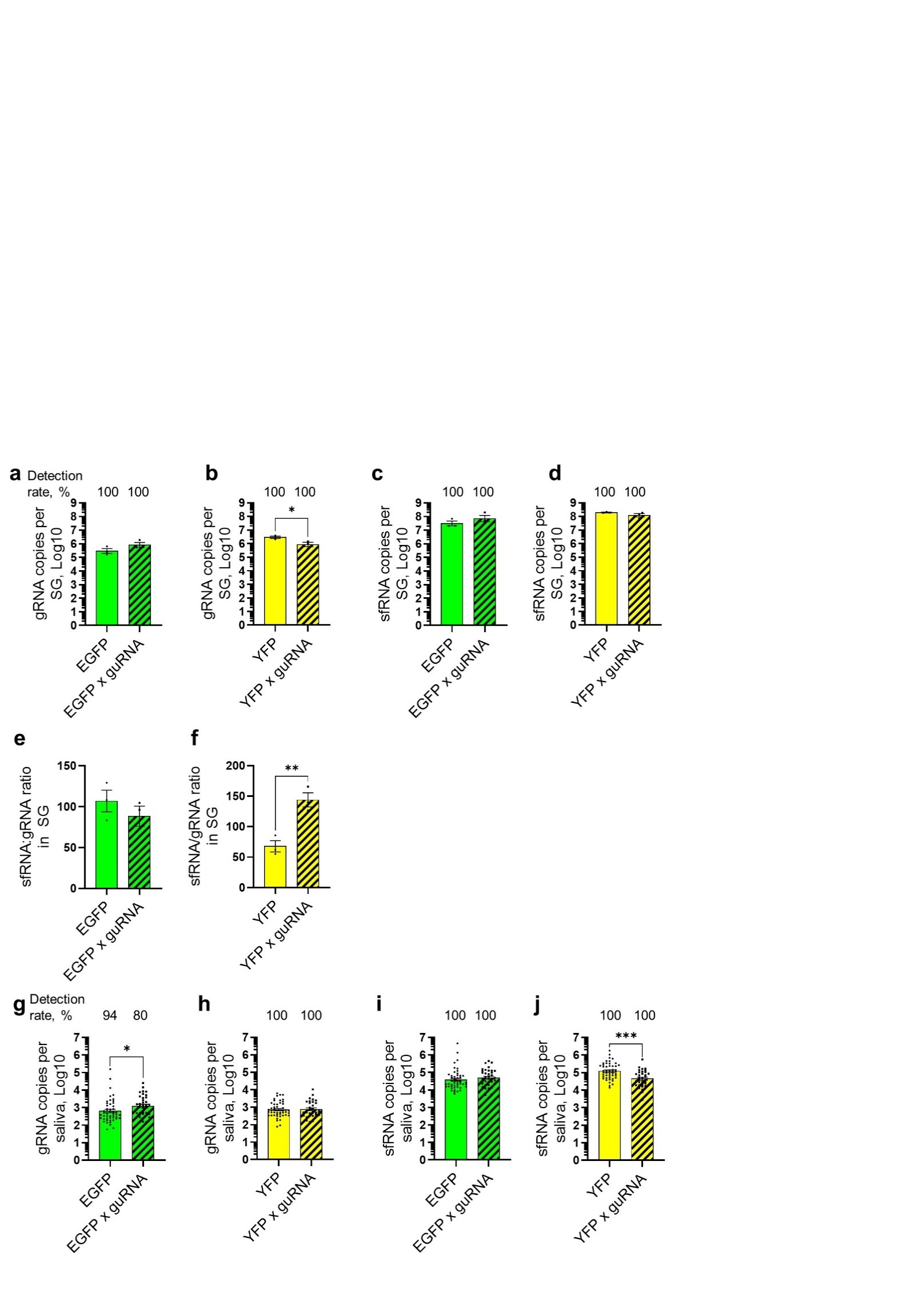

Fig. S26 | Levels of gRNA and sfRNA in salivary glands and saliva from three mosquito transgenic lines with AeSyntenin deletion in SG, related to Fig. 5. **a-f** Levels in SG of gRNA from EGFP and EGFP x guRNA progeny (a) and YFP and YFP x guRNA progeny (b), of sfRNA from EGFP and EGFP x guRNA progeny (c) and YFP and YFP x guRNA progeny (d), and of sfRNA:gRNA ratio from EGFP and EGFP x guRNA progeny (e) and YFP and YFP x guRNA progeny (f). **g-j** Levels in saliva of gRNA from EGFP and EGFP x guRNA progeny (g) and YFP and YFP x guRNA progeny (h), and of sfRNA from EGFP and EGFP x guRNA progeny (i) and YFP and YFP x guRNA progeny (j). a-d,g-j Bars represent geometric mean ± 95% C.I. and dots indicate repeats. e,f Bars represent mean ± s.e.m. and dots indicate repeats. n for a-f, 3. n for g-j, 36-49. *, p < 0.05; **, p < 0.01; ***, p < 0.001, as determined by T-test.

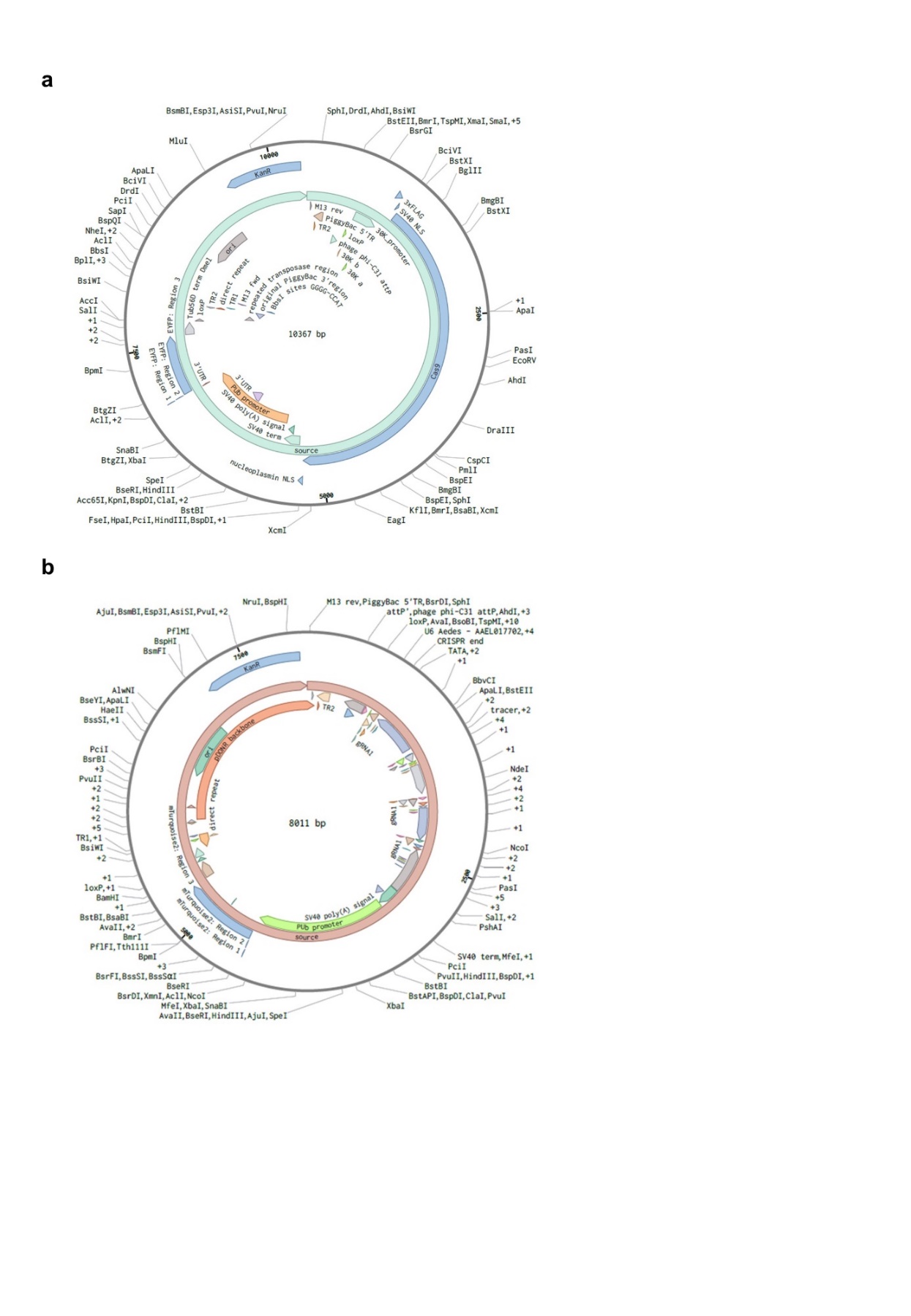

Fig. S27 | Representation of plasmids to generate transgenic mosquitoes, related to Fig. 5. **a** pB30K-Cas9inpJM0142 plasmid to generate Cas9 expressing mosquitoes. **b** pJM140-4gRNAAAEL005391 plasmid to generate guide RNA expressing mosquitoes.

### Table S1. Primer list.

| **Gene** | **ID** | **Forward or**  **reverse** | **Primers** | **Size, bp** |
| --- | --- | --- | --- | --- |
| **qPCR primers** | | | | |
| AeSyntenin | AAEL005391 | Forward | CACGATCCATAAAAACGCGTGA | 101 |
|  |  | Reverse | ATGCTTTTGCGGCAATTGGT |  |
| Actin | AAEL001928 | Forward | GAACACCCAGTCCTGCTGACA | 66 |
|  |  | Reverse | TGCGTCATCTTCTCACGGTTAG |  |
| RPS7 | AAEL009496 | Forward | TCAGTGTACAAGAAGCTGACCGGA | 128 |
|  |  | Reverse | TTCCGCGCGCGCTCACTTATTAGATT |  |
| DENV gRNA |  | Forward | CTCCCTGAGTGGAGTGGAAG | 177 |
|  |  | Reverse | ACACGCACCACCTTGTTTTG |  |
| DENV non-mutated 3’UTR/sfRNA |  | Forward | GTGAGCCCCGTCCAAGG | 170 |
|  |  | Reverse | GCTGCGATTTGTAAGGG |  |
| DENV Pk1 3’UTR/sfRNA |  | Forward | AACCATGGAAGCTGTACGCA | 101 |
|  |  | Reverse | AGCTTCATCTCACCTTGGGC |  |
| DENV Ctl. RNA |  | Forward | TGGCAGTGGTGTCCGTTTCC | 187 |
|  |  | Reverse | ATGCTCACCATCCCGACTGC |  |
| WNV gRNA |  | Forward | ATTCGGGAGGAGACGTGGTA |  |
|  |  | Reverse | CAGCCGCCAACATCAACAAA |  |
| WNV 3’UTR/sfRNA |  | Forward | AGTTGAGTAGACGGTGCTGC | 149 |
|  |  | Reverse | CCGTAGCGTGGTCTGACATT |  |
| ZIKV gRNA |  | Forward | TTGGTCATGATACTGCTGATTGC |  |
|  |  | Reverse | CCTTCCACAAAGTCCCTATTGC |  |
| ZIKV 3’UTR/sfRNA |  | Forward | GCTGGGAAAGACCAGAGACT |  |
|  |  | Reverse | CTATTCGGCGATCTGTGCCT |  |
| **T7 primers** | | | | |
| 3’UTR + tobramycin |  | Forward | ATATAATACGACTCACTATAGGGGAGAAGACGACCGACCAGAATCATGC | 482 |
|  |  | Reverse | AGAACCTGTTGATTCAACAGCAC |  |
| Ctl RNA + tobramycin |  | Forward | ATATAATACGACTCACTATAGGGGAGAAGACGACCGACCAGAATCATGC | 482 |
|  |  | Reverse | GGTCCTGTCATGGGAATGTC |  |
| ∆SL fragment |  | Forward | CGGGTATGTGCGTCTGGATCCTATCCACCTGAGAAGGTGTAAAAAATC | 287 |
|  |  | Reverse | AGAACCTGTTGATTCAACAGCAC |  |
| ∆DB fragment |  | Forward | CGGGTATGTGCGTCTGGATCCTATCAGGTCGGATTA C | 216 |
|  |  | Reverse | AGATTTTTTACACCTTCTCAGGTGGAGC |  |
| ∆3’SL + tobramycin |  | Forward | ATATAATACGACTCACTATAGGGGAGAAGACGACCGACCAGAATCATGC | 401 |
|  |  | Reverse | CAGCACCATTCCATTTTCTGGCG |  |
| dsLacZ |  | Forward | TAATACGACTCACTATAGGGACACCAACGTGACCTATCCC | 318 |
|  |  | Reverse | CCGCCACATATCCTGATCTT |  |
| dsAeSyntenin | AAEL005391 | Forward | TAATACGACTCACTATAGGGGAGACAGCCAAAAAGCAGGA | 243 |
|  |  | Reverse | CAACTCTCGAATTCCGTTGG |  |
| DENV gRNA qPCR template |  | Forward | TAATACGACTCACTATAGGGGGCTCCCTGAGTGGAGTGGAAG | 234 |
|  |  | Reverse | ACACGCACCACCTTGTTTTG |  |
| DENV ∆Pk1 sfRNA qPCR template |  | Forward | TAATACGACTCACTATAGGGAACCATGGAAGCTGTACGCA |  |
|  |  | Reverse | AGAACCTGTTGATTCAACAGCAC |  |
| DENV Ctl. RNA qPCR template |  | Forward | TAATACGACTCACTATAGGGGCAGCTGGACTACTCTTGAG | 482 |
|  |  | Reverse | GGTCCTGTCATGGGAATGTC |  |
| DENV in vitro transcribed sfRNA |  | Forward | TAATACGACTCACTATAGGGGAAGTCAGGTCGGATTAAGC | 403 |
|  |  | Reverse | AGAACCTGTTGATTCAACAGCAC |  |
| DENV in vitro transcribed Ctl. RNA |  | Forward | TAATACGACTCACTATAGGGGCAGCTGGACTACTCTTGAG | 483 |
|  |  | Reverse | GGTCCTGTCATGGGAATGTC |  |

DataSet S1. Analyzed MS data for RNA-affinity chromatography, related to Fig. 3.

DataSet S2. Analyzed MS data for co-IP AeSyntein, related to Fig. 4.

DataSet S3. pB30K-Cas9inpJM0142 plasmid sequence, related to Fig. 5.

DataSet S4. pJM140-4gRNAAAEL005391 plasmid sequence, related to Fig. 5.
